# Discovery of Covalent Wild-Type Isocitrate Dehydrogenase 1 (IDH1) Inhibitors Targeting Cys269

**DOI:** 10.64898/2026.09.24.753951

**Authors:** Charles. P. Brown, Elliot. O. Harmer, Ariba Azam, Charlie Sharp, Sam Horrell, James. A. Bull, Alan Armstrong, David. J. Mann

**Affiliations:** Department of Chemistry, Imperial College London, Molecular Sciences Research Hub, White City Campus, Wood Lane, London, W12 0BZ (UK); Department of Life Sciences, Imperial College London, South Kensington Campus, London SW7 2AZ (UK)

## Abstract

Isocitrate dehydrogenase 1 (IDH1) catalyses the interconversion of isocitrate and α-ketoglutarate and is overexpressed in several cancers, supporting tumour survival and treatment resistance. Inhibitors targeting the oncogenic mutant forms of IDH1 also inhibit the wild-type (WT) protein, but lose substantial potency due to greater competition with native substrates (isocitrate and Mg^2+^). We hypothesised that a covalent inhibitor may be more effective at overcoming the higher substrate affinity of wild-type IDH1. Here, a covalent fragment-based approach was used to develop IDH1-C269 selective inhibitors which prevent the key regulatory segment from forming an α-helix required for catalytic competency. Following the identification of C269-selective fragments, we used X-ray crystallography to characterise IDH1-fragment bound complexes and guide structure-based design efforts. Inspired by the unexpected detection of bound isocitrate molecule in an X-ray crystal structure, fragment expansion yielded a series of compounds exploiting both an adjacent site and a unique water-mediated hydrogen bonding network. This series achieved strong potency (reaching IC_50_ = 47.5 nM, a 155-fold improvement in potency relative to the original fragment), retained activity in the presence of competing Mg^2+^, showed selectivity against IDH2 and reduced NADPH concentrations in a relevant PDAC cell model. Together, these findings present a mechanistic rationale for the covalent targeting of wild-type IDH1 and provide structurally validated inhibitors for further development.

**Graphical Abstract:** 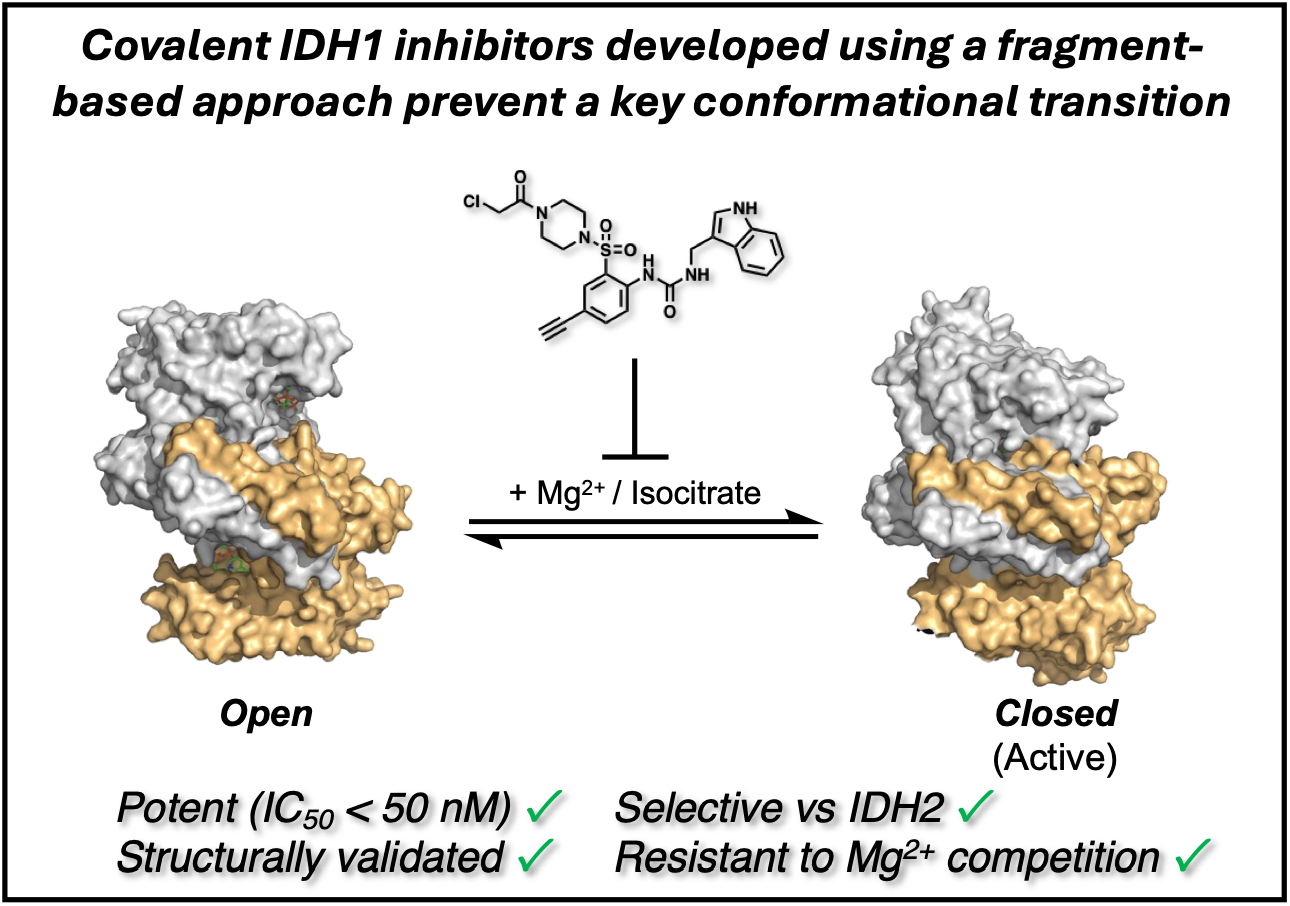

## Introduction

Isocitrate dehydrogenase 1 (IDH1) is a cytosolic, homodimeric protein that catalyses the NADP(H) and Mg^2+^-dependent interconversion of isocitrate and α-ketoglutarate.^1^ IDH1 has historically received significant interest due to the enrichment of mutant forms in gliomas and acute myeloid leukaemia. Mutations substituting R132 for histidine, cysteine, valine or glutamine cause IDH1 to catalyse a neomorphic activity whereby α-ketoglutarate is reduced to form the oncometabolite (*R*)-2-hydroxyglutarate.^2-4^ Mutant IDH1 inhibitors have been developed to inhibit this neomorphic activity, resulting in three FDA-approved therapies (ivosidenib, vorosidenib and olutasidenib) for the treatment of mutant cancers.^5-7^

Interest in targeting wild-type IDH1 has grown in recent years following reports that it is often overexpressed in cancer (including in cancers with significant unmet clinical need such as pancreatic ductal adenocarcinoma (PDAC) and glioblastoma) and that overexpression correlates with poorer survival outcomes.^8-11^ The role IDH1 plays in cancer is likely multifaceted. Early work showed it functions in reductive glutaminolysis – a process by which TCA cycle intermediates are replenished from glutamine.^12-15^ Overexpression of IDH1, and the resulting increase in flux through its oxidative reaction, can raise NADPH levels. This helps buffer elevated ROS and supports the heightened demand for macromolecular synthesis.^8, 9, 16^ Consistent with this function, genetic or pharmacological inhibition of IDH1 decreases cytosolic NADPH concentrations in several tumour types. By increasing NADPH production, IDH1 overexpression may also contribute to therapeutic resistance. For example, IDH1 was identified as a mediator of resistance to 5-fluorouracil in PDAC and radiation sensitivity in glioblastoma.9, ^17^

Although IDH1 has a well-validated function in several cancers, it is logical to be concerned about the possible toxicity implications of targeting a TCA cycle enzyme. Fortunately, increasing evidence suggests the majority of TCA cycle flux is catalysed by the mitochondrial isoform IDH3, and instead IDH1 and 2 likely predominantly function to regulate isocitrate/α-ketoglutarate levels and maintain reductive balance.^18^ Hence *Idh3α*^−/−^ mice are non-viable, whilst *Idh1*^−/−^ mice are healthy and fertile, but show increased vulnerability to oxidative insults.^19, 20^ *Idh2*^−/−^ mice display more severely affected phenotypes, including hearing loss and cardiac issues.^21, 22^

Combining the role IDH1 plays in cancer, evidence of therapeutic benefit resulting from inhibition, and the healthy phenotypes of *Idh1*^−/−^ mice, IDH1 represents an interesting, yet underexplored drug target. Motivated by these findings, a clinical trial is currently ongoing investigating the use of the mutant IDH1 inhibitor ivosidenib in combination with FOLFIRINOX in PDAC (NCT05209074). However, mutant IDH1 inhibitors are highly selective against the wild-type isoform, primarily as a result of the much tighter competing substrate affinity relative to the mutant.^23, 24^ For example, although ivosidenib binds to both WT and IDH1^R132H^ with approximately equal affinity, it is 400-fold less potent against the wild-type isoform in the presence of endogenous substrates. *In cellulo* studies in MIA PaCa-2 cells also found that ivosidenib potency increases substantially at lower Mg^2+^ concentrations.^16, 23^ Mutant IDH1 inhibitors will therefore likely only be suitable for targeting the wild-type form in low Mg^2+^ environments.

Covalent strategies of modulating protein activity have found considerable success in drugging challenging targets, owing to distinct advantages over traditional non-covalent approaches.^25^ Notably, covalent inhibitors have previously demonstrated the ability to better outcompete tightly binding endogenous substrates.^26^ We therefore hypothesised that a covalent strategy may be preferable when targeting wild-type IDH1 (Fig. 1).

**Figure 1.**
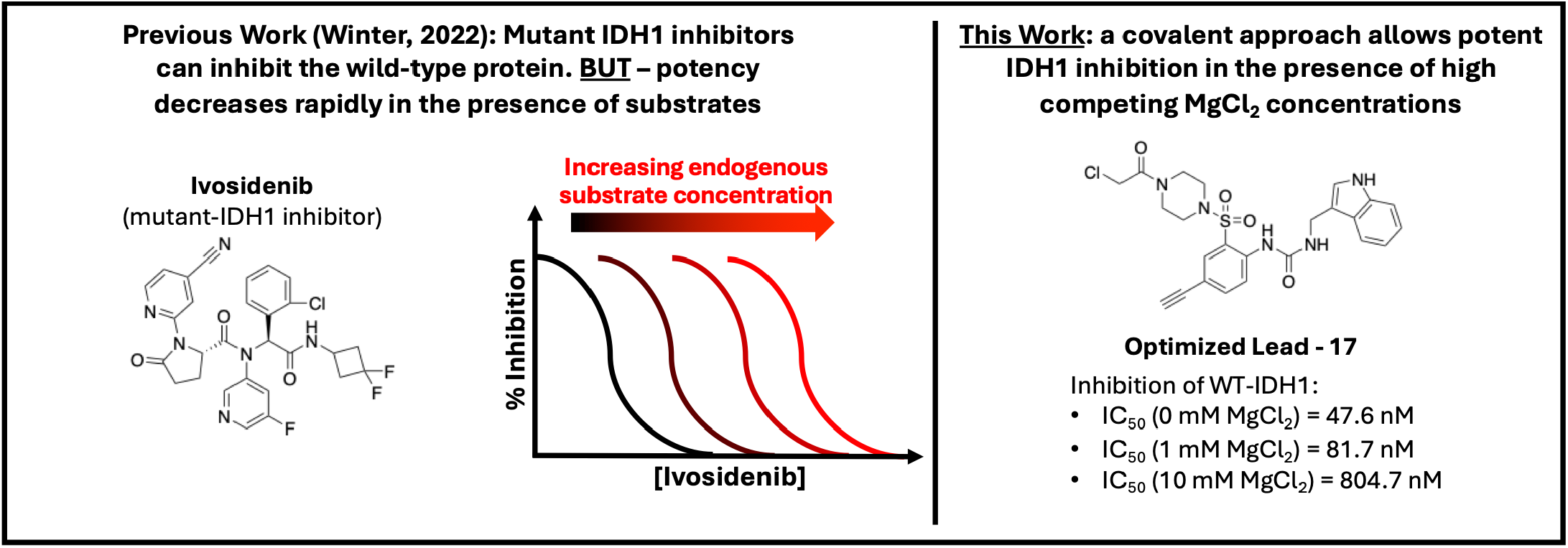
A conceptual overview of this work.

Here, we report the application of a structure-guided covalent-fragment approach that yielded a potent covalent IDH1 inhibitor (IC_50_ = 47.6 nM) with demonstrated proteomic target engagement, cell activity, and strong selectivity against the IDH2 isoform. These findings provide both starting points for the further development of covalent IDH1 inhibitors and an example of the successful application of an electrophilic fragment-based ligand discovery approach. We suggest that the strong potency achieved by the covalent inhibitor reported here, despite the presence of high competing Mg^2+^ concentrations, demonstrates the advantage of a covalent approach towards IDH1 inhibition.

## Results and Discussion

### Probing the Potential for Allosteric Covalent Inhibition

*Copasi* was used to model IDH1 occupancy for both non-covalent and covalent inhibitors of drug-like potency, both in the absence and presence of substrates.^27^ Aligned to our hypothesis, our modelling predicted that a covalent inhibitor should retain far greater levels of target engagement in the presence of substrates than a drug-like non-covalent equivalent, consistent with previously reported experimental data (Fig. S1).^16^

Confident that the modelling provided a theoretical rationale for a covalent approach, we subsequently implemented site-directed mutagenesis to probe the potential for inhibition through covalent IDH1 engagement. C269 is evolutionarily well-conserved (Fig. S2) and is located in the allosteric pocket occupied by several mutant IDH1 inhibitors, so was prioritised for exploration.^6, 28^ An IDH1^C269W^ construct was generated for use in simulating inhibitor-conjugated IDH1. Pleasingly, IDH1^C269W^ was catalytically inactive (Fig 2A), indicating the tethering of a bulky ligand to this position will likely inhibit IDH1 activity.

**Figure 2.**
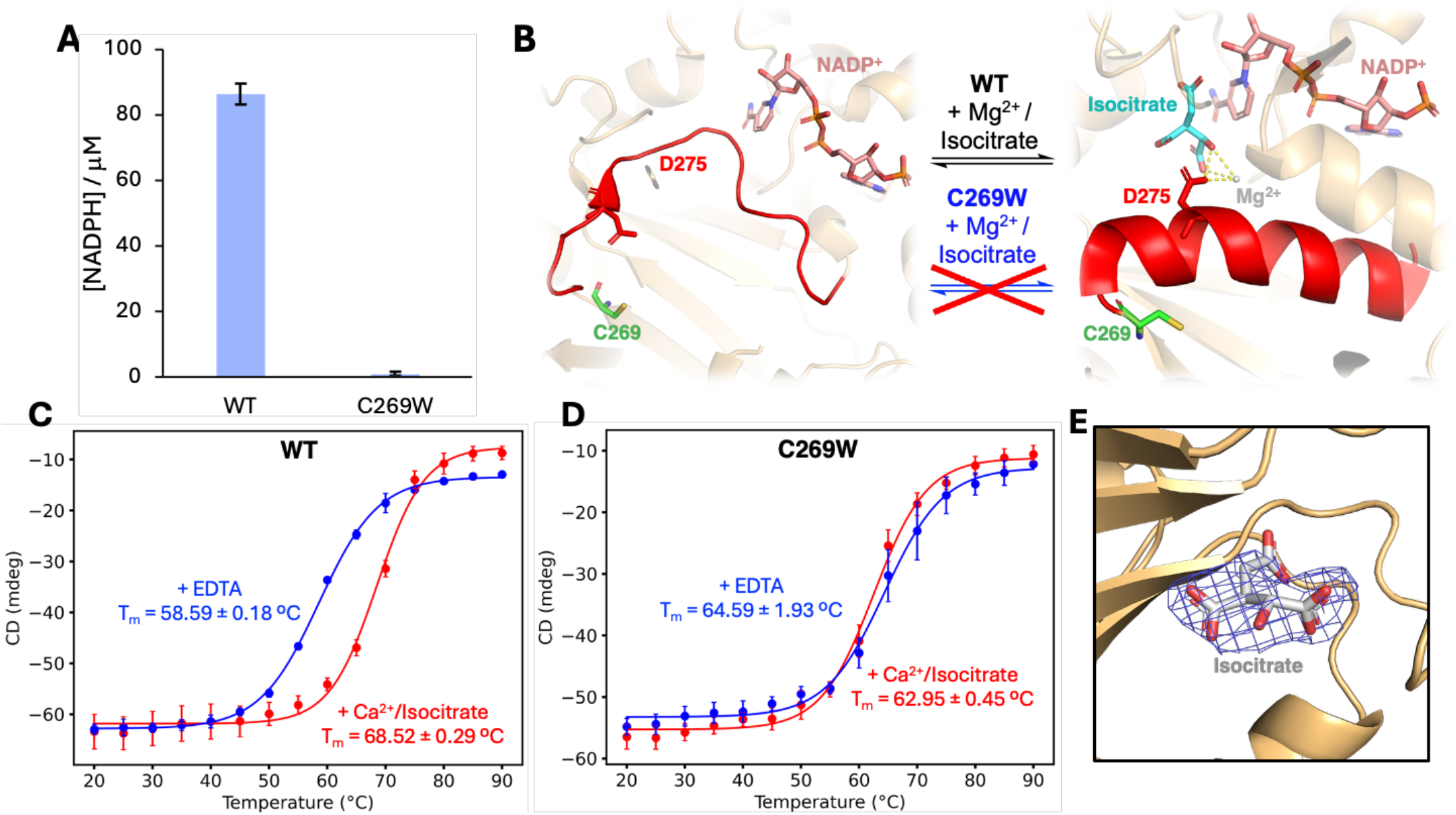
(**A**) NADPH generated by IDH1 and IDH1^269W^ (10 nM) following 60 mins incubation with NADP^+^ (250 µM), isocitrate (100 µM) and MgCl_2_ (10 mM) as determined by monitoring absorbance at 340 nm (N = 4), showing the mutant form is inactive. (**B**) A schematic illustrating the proposed mechanism by which the C269W mutation prevents the transition between the open and closed conformations by sterically blocking the formation of the regulatory segment α-helix (PDB-IDs = 1T0L and 1T09).^1^ (**C/D**) Melting curves for (**C**) IDH1 (**D**) and IDH1^C269W^ as determined via temperature-dependent circular dichroism (N = 3). (**E**) Isocitrate (grey) resolved bound to IDH1 (2F_o_-F_c_ density is displayed in blue at 0.8 σ).

We next sought to characterise the mechanism by which the C269W mutation inhibited catalysis, which we speculated may relate to the high conformational plasticity of the proximal ‘regulatory segment’ (residues 271-286). IDH1 switches from an inactive ‘open’ to an active ‘closed’ conformation on exposure to substrates (Mg^2+^ and isocitrate), during which the regulatory segment switches between an unstructured loop and α-helix.^1^ Formation of the α-helix is essential for catalysis as it completes the active site architecture by forming the bottom of the substrate-binding pocket. This includes positioning the key D275 residue for coordination with Mg^2+^ and isocitrate. Due to the close proximity between C269 and the regulatory segment, we hypothesised that the C269W mutation would inhibit catalysis by sterically blocking the regulatory segment from adopting the α-helical conformation (Fig 2B).

As the α-helical regulatory segment reduces protein flexibility and enhances hydrophobic packing in the IDH1 core, its formation is associated with significant thermal stabilisation (Fig 2C, measured to be 9.96°C here via temperature-dependent circular dichroism). This thermal stabilisation was abolished in the IDH1^C269W^ protein (Fig 2D), consistent with the mutation preventing conformational switching. We further confirmed this mechanism using X-ray crystallography, where we co-crystallised IDH1^C269W^ in the presence of isocitrate (250 µM)/CaCl_2_ (1 mM). Ca^2+^ is used here in place of Mg^2+^ to generate catalytically inactive IDH1 in the closed conformation.^1^ The resulting structure demonstrated that IDH1^C269W^ was restricted to the catalytically inactive open conformation, despite the presence of endogenous substrates.

To our surprise, in the IDH1^C269W^ X-ray crystal structure we resolved electron density consistent with an isocitrate molecule bound in a novel position within the allosteric pocket distinct from its traditional substrate binding site (Fig 2E). This allosteric site has been previously exploited by other groups in the development of reversible, non-covalent mutant-selective inhibitors (such as by a 2-pyridone motif in the FDA-approved olutasidenib).^29^ It has been speculated that native substrates may bind to and modulate IDH1 activity^30^, and a recent report revealed that IDH1^R132H^ activity can be regulated through this pocket via the autopalmitoylation of C269.^31^ We suggest that the presence of isocitrate in this position may indicate that isocitrate (or other TCA cycle intermediates) may regulate IDH1 activity by binding to this pocket. For example, previous work has found that α-ketoglutarate inhibits IDH1 at high concentrations – possibly it exerts this activity through this pocket.^32^

In all, the combined computational modelling and mutagenesis experiments described suggested that covalent IDH1 inhibition via C269 engagement would be an effective method of overcoming the tight competition with native substrates.

### An Irreversible Tethering Screen Identified C269-Targeting Covalent Fragments

We next performed a screen of cysteine-reactive covalent fragments to identify IDH1-C269 selective electrophiles for further optimisation. We screened a 1,600-member covalent fragment library, consisting of both commercial and in-house synthesised fragments, and utilising acrylamide, chloroacetamide, vinyl sulfonamide and other warhead chemistries.

We used the quantitative irreversible tethering (qIT) assay here as a primary screening method.^33^ The qIT assay is a fluorescence-based assay whereby a fragment’s reactivity with both the target cysteine and glutathione (GSH, an unstructured tripeptide control used here as a proxy for intrinsic electrophile reactivity) are measured and used for hit selection. Fragments which conjugate the target cysteine at an enhanced rate relative to GSH are presumed to engage in productive non-covalent interactions, so are designated ‘hits’ and pursued further, whilst those lacking any rate acceleration are excluded. By considering the rate of GSH reactivity, pan-reactive electrophiles can be excluded, allowing the selection of fragments which display genuine molecular recognition-accelerated reactivity with a target cysteine residue.

Here, we ranked fragments using the rate-enhancement factor (REF, a metric calculated by dividing the rate of reaction with IDH1-C269 by that with glutathione) and selected those with a REF > 3 for further exploration. To ensure we were measuring fragment reactivity solely at C269, we performed the screen using a single cysteine IDH1 construct (IDH1^C73S, C114S, C297S, C379S^). We then orthogonally validated any hits using LC-MS (see Fig. S3-9), by selecting those which caused a single shift of the correct molecular weight. From a screen of 1,600 fragments, we identified 7 validated hits using this sequence (Fig. 3A), including molecules bearing acrylamide, vinyl sulfonamide and chloroacetamide warhead chemistries. Notably **1** was previously reported as a hit against IDH1-C269 from a proteomic screen, but to our knowledge was not further explored.^34^ An IDH1-binding chloroacetamide (**2**) reported from the same proteomic screen was also characterised by qIT (REF = 5.3) and was included in our pool of hits.

**Figure 3.**
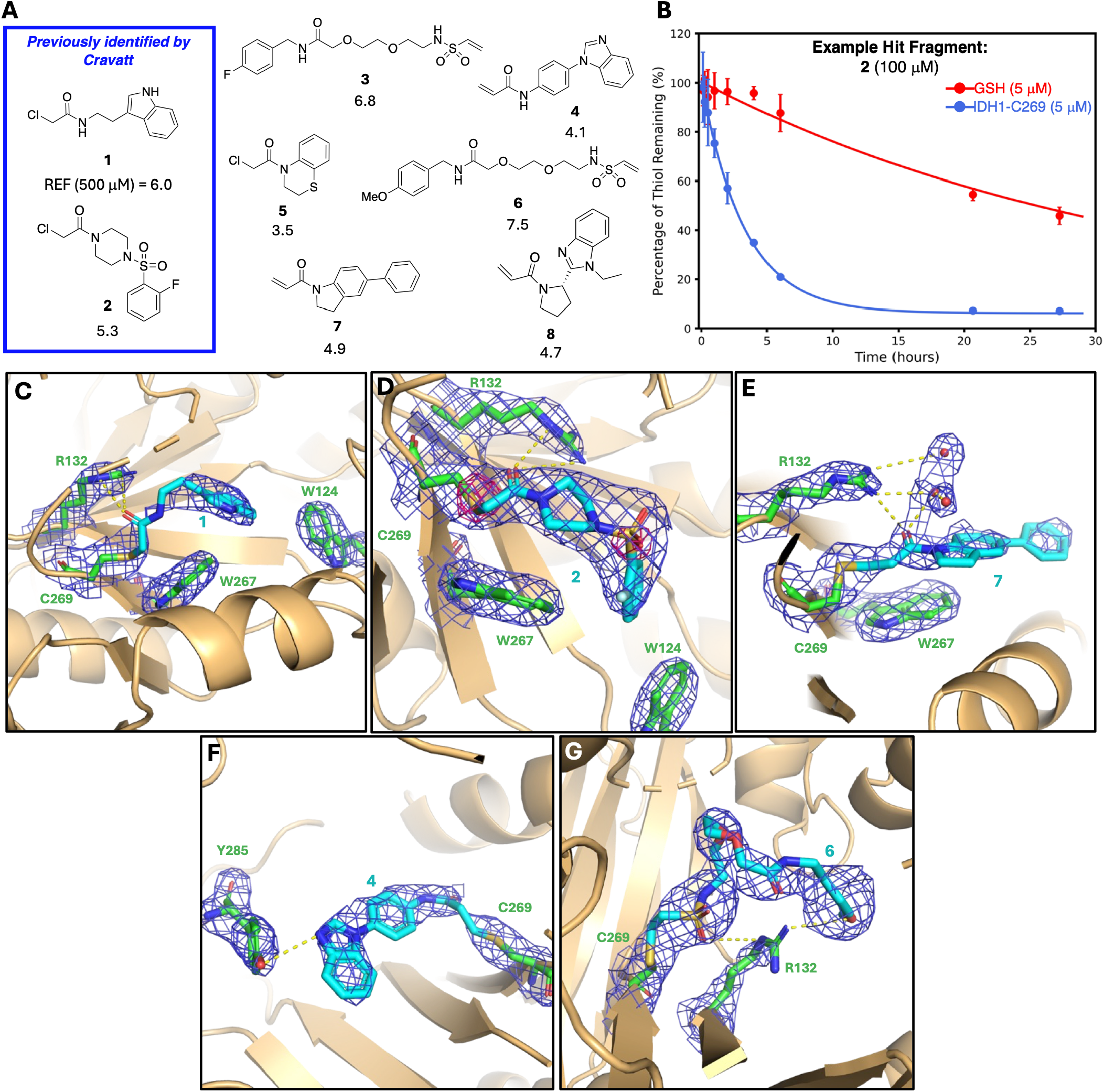
(**A**) IDH1-selective covalent hit fragments and measured REF values. (**B**) An example qIT-assay measurement for hit fragment **2**, showing the measured reactivity of **2** with GSH and IDH1-C269. (**C**-**G**) IDH1-fragment cocrystal structures focusing on interactions speculated to be key for fragment-binding: in each case IDH1 is displayed with cartoon representations (light orange) with the bound ligand (cyan) and key residues (green) represented as sticks. The 2F_O_ - F_c_ density is displayed as dark blue webbing at 1σ, and anomalous difference Fourier map (**D**) in pink at 4 σ.

We next sought to ensure that the mutations introduced into the single cysteine IDH1 screening construct did not cause false positive results by investigating whether hit fragments conjugate C269 in IDH1 when all cysteine residues are present. Previous reports have utilised intact-protein LC-MS as a primary covalent-fragment screening method, utilising the presence of a predominantly single-labelled species as an indication of accelerated reactivity at a single cysteine.^35^ Here, we adopted a similar approach, and incubated IDH1 (5 µM) with fragments **1, 6** and **8** (500 µM) for 180 mins, then investigated protein labelling via intact-protein LC-MS (Fig. S10 – 12). Fragments **1** and **8** each caused several protein modifications, although IDH1 bearing a single fragment-label was the predominant species in each case. Fragment **6** only caused a single modification (despite the presence of 5 other competing cysteines), consistent with accelerated reactivity at C269 relative to the other cysteines.

We next used a similar approach to investigate whether fragments compete with substrates for IDH1 occupancy (Fig. S10 – 12). Previous reports using hydrogen-deuterium exchange mass spectrometry found that IDH1 forms a complex with Mg^2+^, causing a partial folding of the regulatory segment.^36^ We therefore anticipated that C269-selective fragments may compete with Mg^2+^ for target occupancy. Here, the single labelling character was abolished for **1, 6** and **8** when incubated with IDH1 in the presence of MgCl_2_ (10 mM); instead, the unlabelled intact protein was the predominant species in each case. The loss of the single-labelling pattern provides strong evidence for the C269-conjugating fragments competing with Mg^2+^ for target occupancy. This data serves as an indication of target engagement resulting from genuine molecular recognition, instead of non-specific reactivity.

Finally, we investigated whether labelling of a single constituent of the IDH1 dimer caused total activity inhibition. Previous work has found that IDH1 employs a half-site mechanism, whereby activity alternates between the two monomers during catalysis. We hypothesised that this mechanism could result in conjugation of one monomer inhibiting the total activity of the dimer. To test this, **2** was incubated with IDH1 (100 µM,120 mins), before substrates were added and residual activity measured. Despite only conjugating approximately 50% of the C269 population (as determined via qIT assay profiling) **2** caused almost complete inhibition of IDH1 activity (Fig. S13), indicating conjugation of one side of the dimer likely inhibits total activity. Utilizing the same fragment concentration and incubation time, **2** failed to inhibit IDH1^C269S^ activity, proving inhibition occurs through C269 conjugation.

### Fragment Co-Crystallisation Reveals Key Interactions with IDH1

An IDH1-fragment co-crystallisation campaign was enacted to inform the downstream medicinal chemistry expansion of fragments. To this end, we established a procedure whereby each fragment was incubated with IDH1 for sufficient time to obtain >60% target occupancy, before the MBP purification tag was removed via incubation with precision protease, yielding IDH1-fragment complexes for the construction of crystallisation screens. Initial screens with apoIDH1 identified conditions (2M (NH_4_)_2_SO_4_, 0.1 M Hepes pH 7.5, 2% *v/v* PEG400) which yielded diffraction quality protein crystals with a large cubic morphology. These conditions were reproducible for IDH1 conjugated with hit fragments, yielding diffraction data to a maximal resolution of 2.15 Å.

From this campaign, 5 IDH1-fragment structures were successfully solved (Fig 3C-G). In each case, electron density was visible protruding from C269 consistent with the relevant fragment’s shape and size, allowing intermolecular interactions dictating molecular recognition to be rationalised. The structure of non-fragment-bound IDH1 was also solved for comparison.

For fragment **1** (Fig. 3C), in monomer A the indole ring system fills a hydrophobic pocket lined by V255, W124, I128, W267, I130 and V281 (likely forming CH-π interactions with W267 and π-π interactions with W124), whilst the warhead carbonyl forms a hydrogen bond with the R132 side chain. The latter interaction is hypothesised to be key by activating the electrophile towards reaction in the pre-conjugation complex via a combination of polarising and optimally positioning the chloroacetamide. In contrast, **1** was less well resolved in monomer B, and adopted a flipped conformation (which we assumed not to be biologically relevant).

Fragment **2** (Fig. 3D) adopted a comparable pose to **1**: it filled the same hydrophobic pocket; formed CH-π interactions with W267 and π-π interactions with W124; and formed activating interactions between the acetamide carbonyl and R132 side chain. However, unlike **1**, fragment **2** retained largely the same binding pose in both monomers. The binding mode of **2** was further validated using anomalous diffraction experiments to confirm the positioning of the sulfur atoms in the ligand and cysteine.

Fragment **7** (Fig. 3E) was only resolved bound to monomer B, and also appeared to engage in π-π interactions with W267. Unexpectedly, a hydrogen bonding network involving the R132 side chain, two water molecules and the acetamide carbonyl of **7** was resolved. It could be speculated that these interactions again catalyse reaction with C269 by a combination of polarising the acrylamide, stabilising the oxyanion intermediate formed during conjugation, and optimally positioning the electrophile for reaction with C269.

Density consistent with fragment **4** (Fig 3F) was resolved bound to monomer A, and appears to form a hydrogen bond between the benzimidazole nitrogen and the Y285 phenol. Similarly, fragment **6** (Fig 3G) was only well resolved bound to monomer A, where the PEG linker forms a U-shape around R132, with the sulfonamide NH and methoxy groups interacting with the R132 side chain.

Comparing these co-crystal structures, interactions with R132 are clearly a critical driver of molecular recognition, as four of the five fragments form an interaction with this residue. Interactions such as these between covalent warheads and cationic residues have been previously noted as being key to catalysing conjugation, for example several KRAS^G12C^ inhibitors exploit an interaction with an adjacent lysine for site-specific activation and optimal warhead positioning.^37-39^ Both **1** and **2** also fill the same hydrophobic groove, indicating this section may be amenable to engagement with small molecule ligands. These interactions were therefore prioritised during later optimisation.

### Expansion Into Putative Isocitrate Binding Pocket Enhances Potency

Following the solving of the IDH1-fragment co-crystal structures, we next began a fragment expansion campaign aiming to develop more potent covalent inhibitors. Fragment **2** stood out as an obvious choice for further expansion as: (1) it interacts symmetrically with IDH1; (2) it exploits the activating interaction with R132; (3) the X-ray crystal structures provided a high degree of certainty about the interactions dictating the IDH1-**2** molecular recognition. Analogues of **2** were therefore prepared and tested for activity.

The benzene sulfonamide group was found to be key for filling the hydrophobic pocket and engaging in π-π interactions, and replacement with a methyl sulfonamide led to a near complete loss of REF (Fig. S14). We hypothesised that the 2-fluoro substituent engaged in hydrophobic interactions with V255, and indeed removal of the fluorine (**10**, Fig. 4B) reduced REF to 4.3 (see Fig 4A for the relative positioning of V255 and **2**). Expanding from this position with bulkier hydrophobic groups enhanced REF substantially, for example 2-methyl (**11**) and 2-*tert-*butyl (**12**) substituted analogues increased REF to 11.7 and 35.8 respectively (note that due to poor solubility these were performed at a compound concentration of 100 µM where **2** displayed a REF of 10.4).

**Figure 4.**
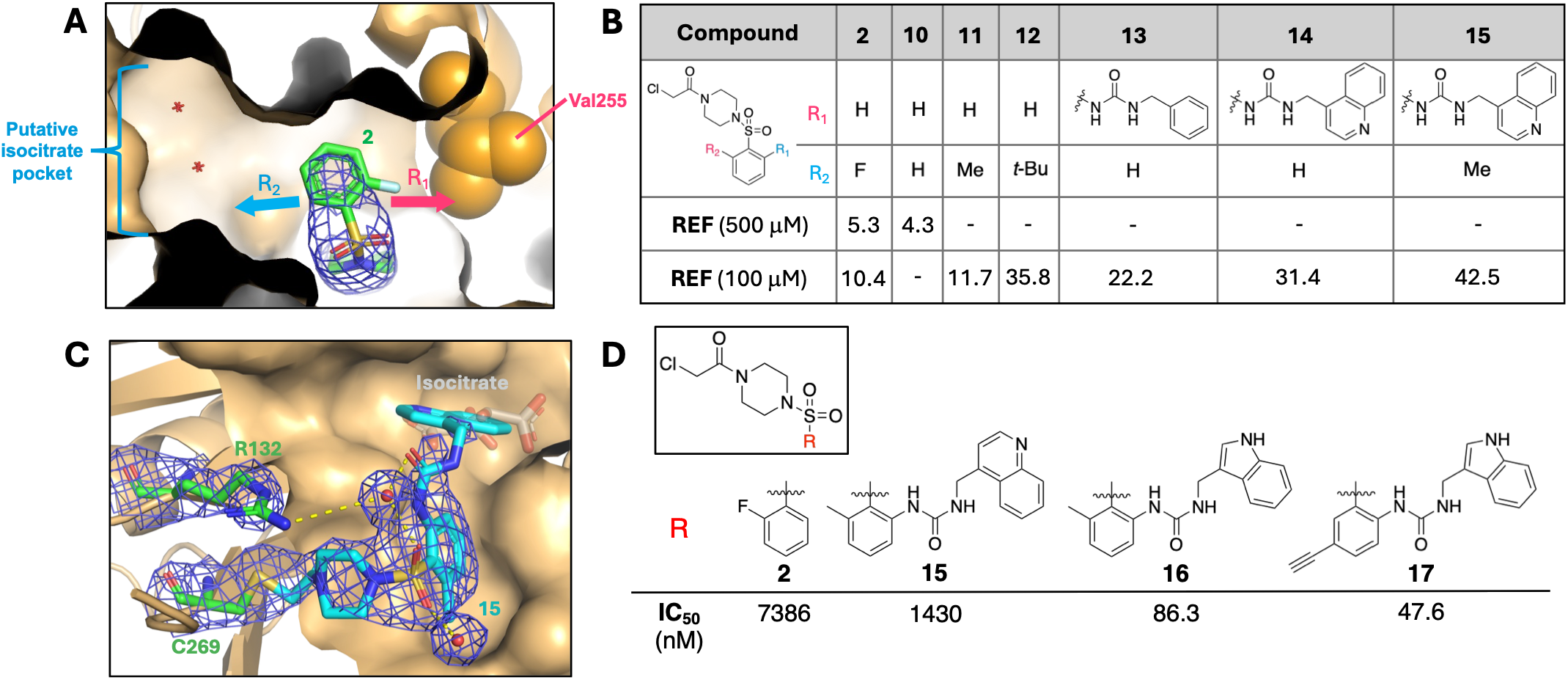
(**A**) IDH1-**2** crystal structure indicating the two vectors explored in analogues (**2** is displayed with a cyan stick representation, IDH1 is displayed with a light orange surface representation, and 2F_O_ - F_c_ density is displayed as dark blue webbing at 1σ). (**B**) Analogues of **2** with associated REF values against IDH1. (**C**) IDH1-**15** cocrystal structure, showing **15** (cyan), key residues (green), water molecules (red), and the relative positioning of isocitrate (grey) in the IDH1^C269W^ structure (2F_O_ - F_c_ density is displayed as dark blue webbing at 1 σ). (**D**) The chemical structures of key analogues **15, 16** and **17**, and measured IC_50_ values following a 2h incubation time with IDH1.

We next looked to expand **2** to engage other areas of the IDH1 allosteric pocket. Upon comparison of the IDH1-**2** and IDH1^C269W^ structures, we noticed that the putative isocitrate binding site previously identified was situated proximal to the site occupied by **2**. In consequence, we hypothesised that expansion to fill this section of the pocket may be an effective strategy to enhance potency (see Fig 4A for the relative positioning of the putative isocitrate binding site and **2**).

However, engaging this adjacent pocket proved to be a challenging analogue design problem, as the angle between the benzene sulfonamide ring and adjacent pocket section necessitated analogues with an appropriate pocket filling motif positioned approximately perpendicular to the original fragment. The docking software *Molmoda* was used here to predict whether hypothesised analogues were capable of filling this space.^40^ Methyl ureas extending from the ortho position of the benzenesulfonamide ring system were predicted to fulfil the required angle, and were thus prioritised in further analogues. Compound **13** was synthesised to test this hypothesis, and promisingly, displayed an enhanced REF (22.2) relative to **2**, despite lacking the V255 engaging hydrophobic group. Expansion of the aromatic pocket filling group to a quinoline (**14**) to enhance contacts within the pocket without compromising solubility further enhanced REF to 31.4. Finally, addition of a V255-engaging methyl group (**15**) further enhanced REF to 42.5.

Compound **15** was selected as an early lead inhibitor, and characterisation revealed it displayed an 11.5-fold enhanced *k*_*inact*_*/K*_*I*_ value relative to **2** (Fig. S21). This was mainly driven through enhanced *K*_*I*_ (32.7-fold tighter), despite a reduced *k*_*inact*_ (2.8-fold reduced). **15** inhibited IDH1 with a half-maximal inhibitory constant (IC_50_) of 1.43 µM (as determined by measuring residual activity following incubation with IDH1 for 2 h in the absence of substrates), representing a 5-fold improvement in inhibitory potency relative to fragment **2** (Fig. 4D).

The IDH1-**15** cocrystal structure was subsequently solved to understand the factors governing molecular recognition (Fig.4C). When bound to monomer A, electron density was visible protruding from C269 consistent **15**. Continuous density suggested the methyl urea linker adopted the hypothesised perpendicular angle, positioning the quinoline ring system in the pocket occupied by isocitrate in the IDH1^C269W^ structure. Unexpectedly, the urea linker, sulfonamide and R132 side chain engaged in a water-mediated hydrogen bonding network, which is speculated to stabilise the urea in the bioactive conformation. Coordination between the acetamide carbonyl and R132 side chain appears to be lost in **15**, likely reducing warhead polarisation in the pre-reaction IDH1-**15** complex, and possibly explaining the reduced *k*_*inact*_ noted relative to **2**. In contrast, weaker electron density was visible bound to monomer B, with **15** not appearing to fill a binding pocket and adopting a pose markedly different to that of **2**. This discrepancy is likely indicative of **15** engaging IDH1 asymmetrically - rapidly conjugating monomer A then more slowly conjugating monomer B.

Further medicinal chemistry efforts aimed to improve potency by replacing the quinoline with other heterocycles and optimising hydrophobic interactions. Pleasingly, we found that an indole-containing analogue (**16**) enhanced potency a further 16.6-fold, and a final analogue bearing a meta-substituted ethynyl substitute (**17**) yielded even greater enhancements in potency (IC_50_ = 47.6 nM), likely due to additional hydrophobic interactions with A258, W124 and I128. In total, through engaging the putative isocitrate-binding pocket and optimising interactions with adjacent hydrophobic residues, a 155-fold improvement in inhibitory potency was achieved between fragment **2** and lead compound **17** (Fig. 4D).

### Lead Compound 17 is Resistant to Mg^2+^ Competition, Selective Against IDH2 and Engages IDH1 in HEK-293 Lysates

We next aimed to characterise the properties of lead compound **17** to evaluate the advantages of a covalent approach. When incubated with recombinant IDH1 (1 µM), **17** (2 µM) caused a single adduct (Fig. S22), consistent with **17** acting as a C269-selective inhibitor.

Owing to the cardiac issues noted in *Idh2*^−/−^ mice, we anticipated that selective inhibition of IDH1 over IDH2 would be an advantageous property. However, isoform selectivity was expected to be challenging as all the residues identified as being key for the IDH1-**2** interaction (C269, W124, R132, V255 and W267) are conserved in IDH2.^21^ Pleasingly, **17** inhibited IDH1 41.2-fold more potently than IDH2 (Fig 5A). We anticipate that this selectivity window results from conformational differences between the IDH1 and IDH2 regulatory segments, as previously reported IDH2 crystal structures (albeit of mutant forms of the enzyme) include an α-helical regulatory segment, even in the absence of substrates (where IDH1 has a disordered regulatory segment which allows access to C269).^30^ This would preclude access to C308 (the equivalently positioned cysteine to C269) if present in solution (see Fig. 5B). The IDH1/2 selectivity window varied greatly between inhibitor analogues (for example the selectivity window decreased to 17.5-fold for **15**, but increased to 184-fold for **16**), indicating a complex structure-activity relationship (Fig. S23).

**Figure 5.**
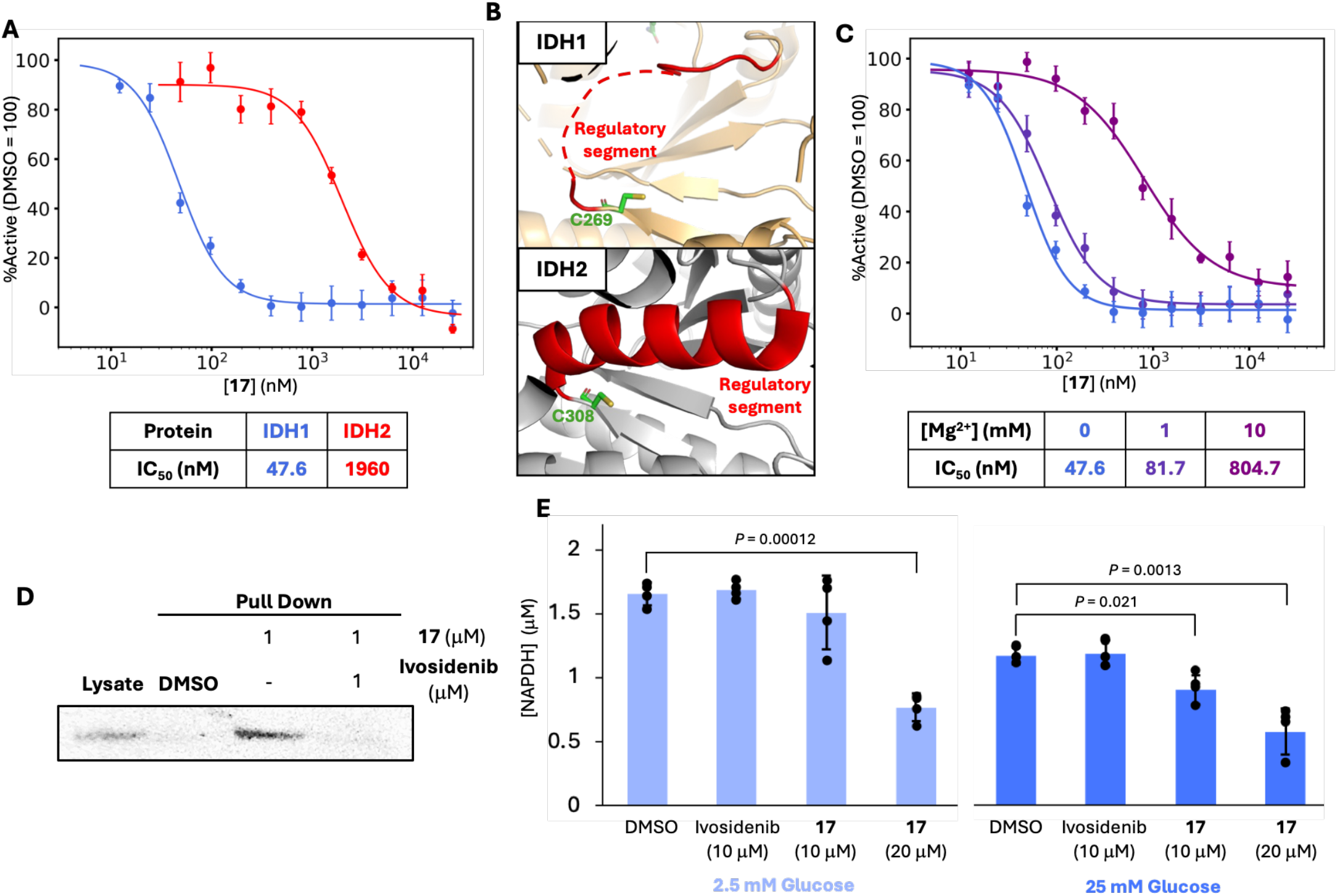
(**A**) Activity assays showing residual IDH1 and IDH2 activity following a 2 h preincubation with **17** (error = 1 standard deviation, N = 4). (**B**) A comparison between regulatory segment conformations in structures of IDH1 and IDH2 (PDB-ID = 5SVN)^30^, providing an explanation for target cysteine accessibility. **(C)** Activity assay curves showing residual IDH1 activity following a 2 h preincubation with **17** in the presence of increasing concentrations of Mg^2+^ (error = 1 standard deviation, N = 4). (**D**) Pull-down assay to assess target engagement: **17** (1 µM) was incubated with HEK-293 cell lysate in the presence or absence of equimolar competing ivosidenib and conjugated to Biotin-N_3_, before neutravidin pull-down was performed and western blotting used to assess the presence of IDH1. (**E**) [NADPH] concentrations following treatment of MIA PaCa-2 cells with **17**.

We next investigated the capacity of **17** to compete with Mg^2+^ for target occupancy. IDH1 natively binds to Mg^2+^, and reversible inhibitors lose much of their activity at enhanced Mg^2+^ concentrations. However, **17** demonstrated a strong capacity to outcompete bound Mg^2+^, and retained potency within 2-fold (IC_50_ = 81.7 nM) at a MgCl_2_ concentration of 1 mM, and sub-micromolar potency (IC_50_ = 804.7 nM) at very high competing MgCl_2_ concentrations (10 mM), (Fig 5C).

Lastly, we investigated the IDH1-**17** interaction in a biological context. Conveniently, the presence of an alkyne handle meant **17** could be used to probe target engagement using copper-catalysed azide alkyne cycloaddition (CuAAC) chemistry to conjugate labelled proteins with TAMRA- or biotin-N_3_ (allowing target engagement to be assessed via in-gel fluorescence or pull-down experiments respectively). **17** (100 nM) was incubated with HEK-293 native cell lysate, then conjugated to TAMRA-N_3_ using Copper-catalysed azide alkyne cycloaddition (CuAAC) chemistry. Unsurprisingly due to the relatively high electrophilicity of the chloroacetamide warhead, numerous fluorescent bands were visualised following probe incubation, including a strong band at approximately the correct molecular weight (~47 kDa) which we hypothesised to be IDH1 (marked with a red arrow in Fig. S24). To confirm this band corresponded to IDH1-C269 labelling, we coincubated HEK-293 cell lysate with both **17** and the FDA-approved mutant-IDH1 inhibitor ivosidenib (which is known to also display activity against wild-type IDH1). In the IDH1^R132H^-ivosidenib crystal structure (PDB-ID = 8T7O), ivosidenib fills an overlapping pocket with that exploited by **17**, so would compete for target occupancy.^28^ Pleasingly, ivosidenib outcompeted **17** for IDH1 allosteric pocket occupancy, as visualised by the loss of the putative IDH1 band in fluorescence gels (Fig. S25). Following a similar protocol, but instead conjugating labelled proteins to biotin-N_3_ and enriching using biotin-streptavidin affinity, **17** (1 µM) pulled-down IDH1 (as visualised via western-blotting, Fig 5D). Coincubation with an equimolar concentration of ivosidenib prior to conjugation to biotin-N_3_ prevented IDH1 pull-down – providing evidence for the proteomic engagement of IDH1 through C269.

Content that **17** engages IDH1 in cell lysates, we next investigated whether the resulting IDH1 inhibition causes a therapeutically desirable effect in a clinically relevant cell line. Previous work has noted that IDH1 is often overexpressed in pancreatic-ductal adenocarcinoma (PDAC) cell lines, especially under the low glucose conditions present in the tumour microenvironment.^9, 16^ This is likely with the purpose of generating additional NADPH equivalents to mitigate the resulting increase in free radical generation. As a result, PDAC cell lines and xenograft models have demonstrated sensitivity to IDH1 inhibition under low glucose conditions. Consistent with past findings, we found that NADPH concentrations were elevated in MIA PaCa-2 cells incubated in low glucose media.^16^ Further, consistent with reversible inhibitors losing potency in the presence of native Mg^2+^ concentrations, ivosidenib (10 µM) failed to reduce NADPH concentrations under either high or low glucose conditions. Pleasingly, treatment with **17** (at 10 and 20 µM) caused a dose dependent decrease in NADPH concentrations in MIA PaCa-2 cells under both high (25 mM) and low (2.5 mM) glucose conditions (Fig. 5E).

These findings suggest that covalent IDH1 inhibition may be an effective method of modulating NADPH levels in tissues overexpressing wild-type IDH1, and could therefore be represent a possible therapy for vulnerable tissues.

## Conclusion

Wild-type IDH1 shows promise as a valuable drug target as it is frequently overexpressed in cancer, functions in anaplerosis and is responsible for generating much of the pool of cytosolic NADPH used to detoxify therapeutic agents. However, owing to the tight competing native substrate affinity, currently marketed mutant IDH1 inhibitors are generally poor wild-type inhibitors under physiological conditions.

We hypothesised that a covalent strategy could be effective at inhibiting IDH1, and using site-directed mutagenesis showed that C269 could be a promising target residue. Covalent fragment screening, followed by crystallographic structure determination and analogue synthesis were then used to create a potent covalent IDH1 inhibitor. Fragment hits representing diverse chemotypes and warhead chemistries were identified and found to engage C269 by interacting with several sections of the surrounding allosteric pocket. These may represent interesting starting points for future optimisation campaigns.

Fragment **2** filled an adjacent hydrophobic groove and was expanded to both more effectively engage adjacent hydrophobic residues and occupy the newly identified putative isocitrate binding pocket, eventually yielding molecules with greater potency. Notably, from X-ray crystal structures, **15** exploited coordination with a conserved water molecule, possibly to maintain the inhibitor in the bioactive conformation. Lead compound **17** was selective against IDH2, maintained potency in the presence of competing Mg^2+^, engaged IDH1 in HEK-293 native lysates and potently inhibited IDH1 *in vivo*.

The large potency improvements observed here attest to the high ligandability of this pocket and suggest a larger medicinal chemistry effort could create inhibitors of drug-like potency. Future work aiming to translate these molecules into more selective probes or drug molecules would likely aim to substitute the chloroacetamide warhead with a less reactive alternative.

Combined, these findings exemplify the three main advantages of targeting IDH1 through covalent engagement of C269: 1) the high ligandability of this pocket means strong potency is achievable; 2) differential regulatory segment dynamics create a large selectivity window over IDH2; 3) potency is retained even in the presence of competing Mg^2+^. We hope that these advantages highlight the potential for covalent IDH1 inhibition and will lead to further exploration of this as a therapeutic strategy.

## Supplementary Materials

Supplementary figures (S1-25), material and methods, and X-ray crystallography data collection and refinement statistics are available in the supplementary materials.

## Acknowledgements

For financial support, we gratefully acknowledge the UK Engineering and Physical Sciences Research Council (EP/S023518/1, UKRIK3018). We also thank Dr. Sofia Caria for her help with the X-ray crystallography data processing.

