## Supplemental Figures, Data, and Methods for "Discovery of Covalent Wild-Type Isocitrate Dehydrogenase 1 (IDH1) Inhibitors Targeting Cys269"

**Supplementary Materials Contents**

1. **Supplementary Figures**

Figure S1: Computational modelling of covalent and non-covalent IDH1 inhibitors
Figure S2: Investigating the evolutionary conservation of C269
Figure S3 – S9: Triaging hit molecules
Figure S10 - S12: Investigating $Mg^{2+}$ competition via LC-MS
Figure S13: Investigating inhibition by **2** through conjugating one constituent of the IDH1 dimer
Figure S14 – S20: qIT reaction curves for analogues 10-15
Figure S21: $k_{inact}/K_I$ determination of **2** and **15**
Figure S22: Intact-protein LC-MS analysis of IDH1-**17**
Figure S23: Gel-based chemoproteomic profiling of **17** in HEK293 cell lysates
Figure S24: Gel-based chemoproteomic profiling of **17** in HEK293 cell lysates with competing ivosidenib

2. **Materials and Methods**

2.1 Biological Materials and Methods
2.2 X-ray Crystallography Methods and Statistics
2.3 Chemistry Materials and Methods
2.4 Analytical Data for Reported Compounds

#### 3. Supplementary References

#### Figure S1 – Computational Modelling of Covalent and Non-Covalent IDH1 Inhibitors

(**A**) **Without substrates:**

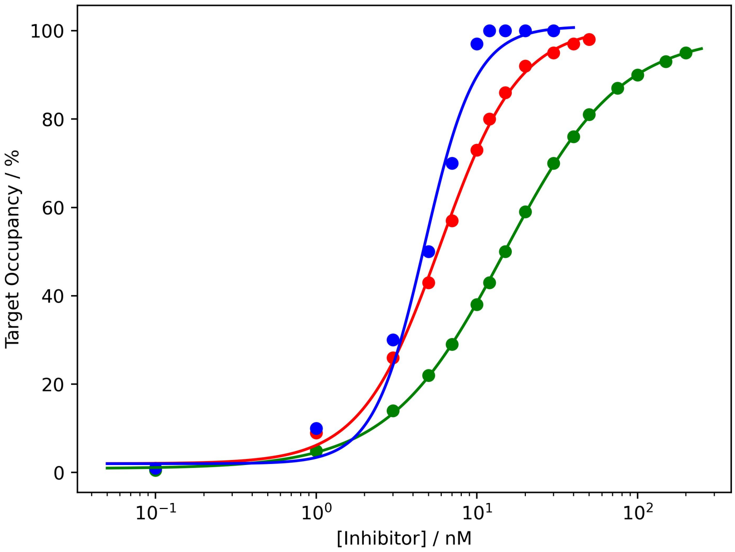


(**B**) **+ 100 μM $MgCl_2$-isocitrate:**

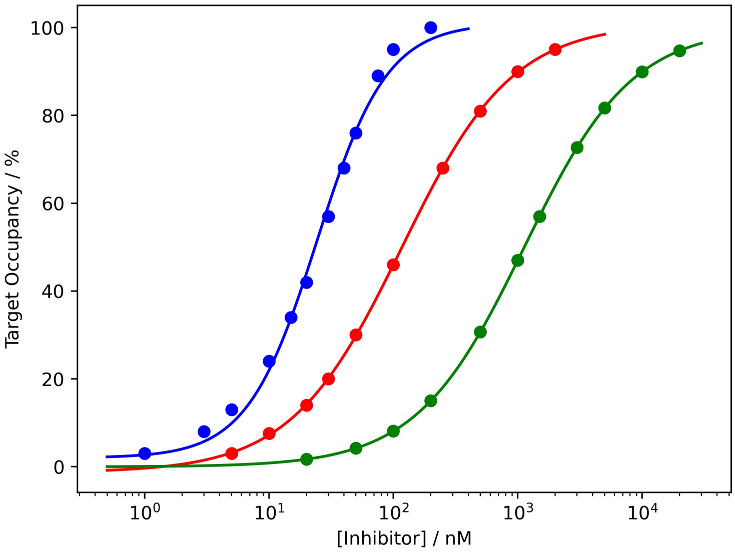


**Figure S1:** Target occupancy vs [Inhibitor] curves for a covalent inhibitor ($k_{inact}/K_I = 1 \times 10^6$ $M^{-1}$ $s^{-1}$, blue) and for non-covalent inhibitors ($K_D = 1$ *nM*, red; $K_D = 10$ *nM*, green) in the absence (**A**) and presence (**B**) of substrates.

#### Figure S2 – Investigating the Evolutionary Conservation of C269

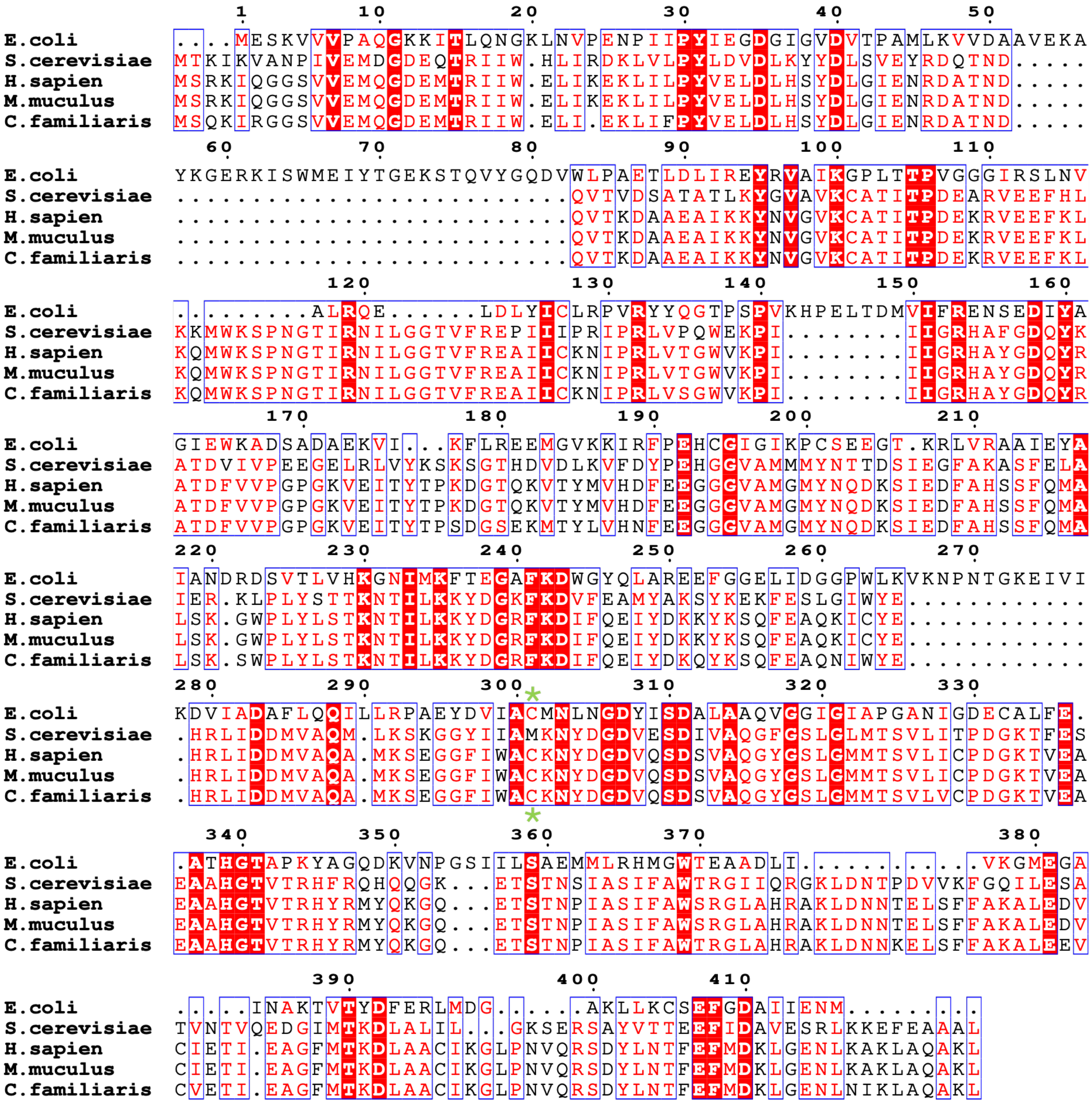

**Figure S2:** A sequence alignment between *Homo sapien, Escherichia coli , Saccharomyces cerevisiae, Muc muculus and Canis lupus familiaris* IDH1, with the key cysteine marked (*). Sequence alignment was generated using the EBI Clustal Omega tool[1], and visualisation generated using *Esprpit 3.2*[2].

#### Figure S3 – Triaging Hit Molecules

**A**

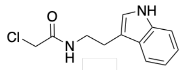


**1**

$k_{obs}/[I]$ (GSH) = 0.0175 $M^{-1}$ $s^{-1}$
$k_{obs}/[I]$ (IDH1) = 0.105 $M^{-1}$ $s^{-1}$
REF = 6.00

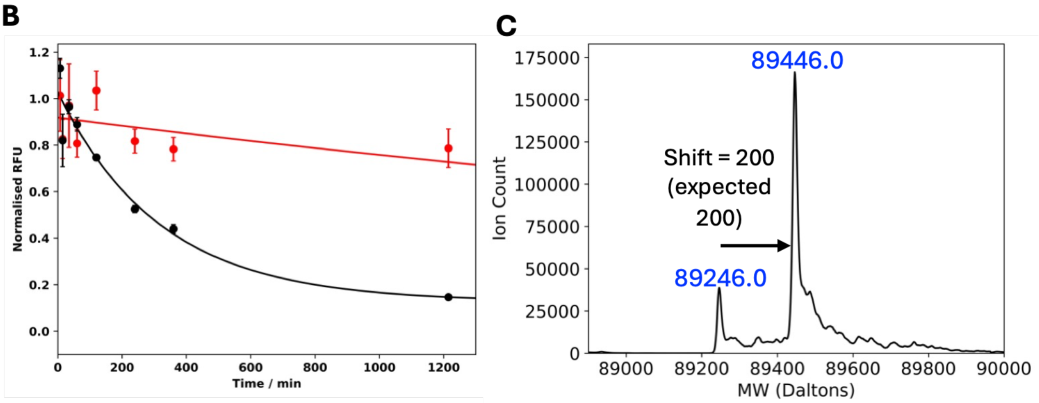


**Figure S3:** Triaging fragment **1**. (**A**) The chemical structure of fragment **1** and calculated rate constants for the reaction with MBP-IDH1$^{Cys-}$ (black) and GSH (red) (N = 3). (**B**) Triplicate qIT repeats show **1** reproducibly reacts at an enhanced rate with MBP-IDH1$^{Cys-}$ relative to with GSH (error = 1 standard deviation, N=3). (**C**) Intact-protein LC-MS analysis of MBP-IDH1$^{Cys-}$ (5 $\mu$M) following pre-incubation with fragment **1** (360 mins).

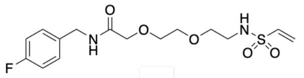


**3**

$k_{obs}/[I]$ (GSH) = 0.0227 $M^{-1}$ $s^{-1}$
$k_{obs}/[I]$ (IDH1) = 0.154 $M^{-1}$ $s^{-1}$
REF = 6.77

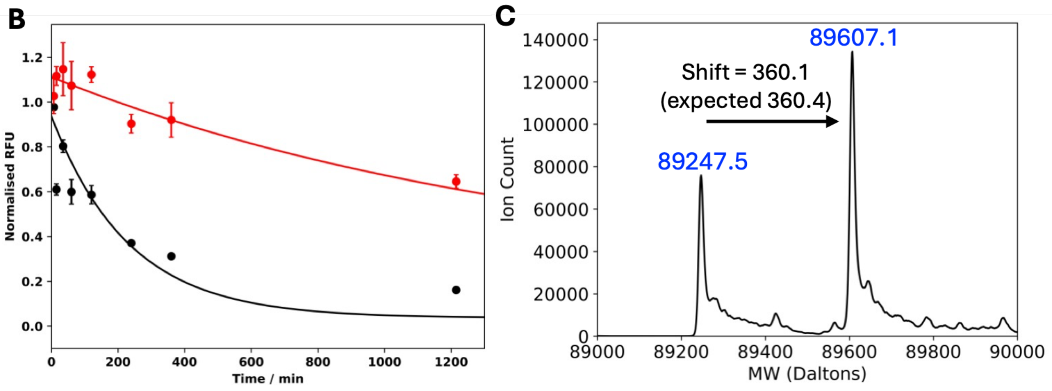


**Figure S4:** Triaging fragment **3**. (**A**) The chemical structure of fragment **3** and calculated rate constants for the reaction with MBP-IDH1$^{Cys-}$ (black) and GSH (red) (N = 3). (**B**) Triplicate qIT repeats show **3** reproducibly reacts at an enhanced rate with MBP-IDH1$^{Cys-}$ relative to with GSH (error = 1 standard deviation, N=3). (**C**) Intact-protein LC-MS analysis of MBP-IDH1$^{Cys-}$ (5 $\mu$M) following pre-incubation with fragment **3** (360 mins).

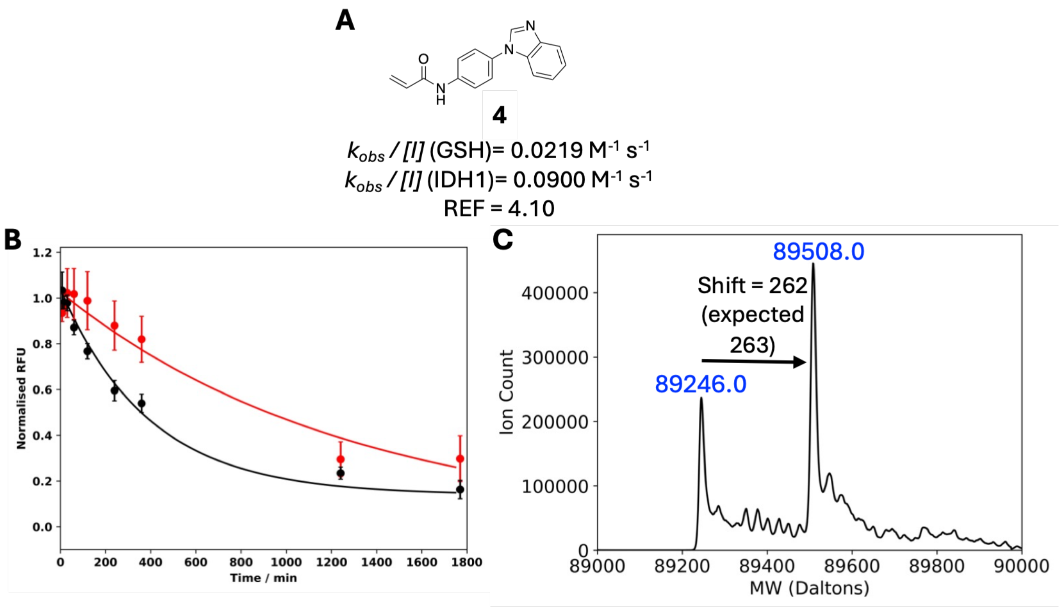


**Figure S5:** Triaging fragment **4**. (**A**) The chemical structure of fragment **4** and calculated rate constants for the reaction with MBP-IDH1$^{Cys-}$ (black) and GSH (red) (N = 3). (**B**) Triplicate qIT repeats show **4** reproducibly reacts at an enhanced rate with MBP-IDH1$^{Cys-}$ relative to with GSH (error = 1 standard deviation, N=3). (**C**) Intact-protein LC-MS analysis of MBP-IDH1$^{Cys-}$ (5 $\mu$M) following pre-incubation with fragment **4** (360 mins).

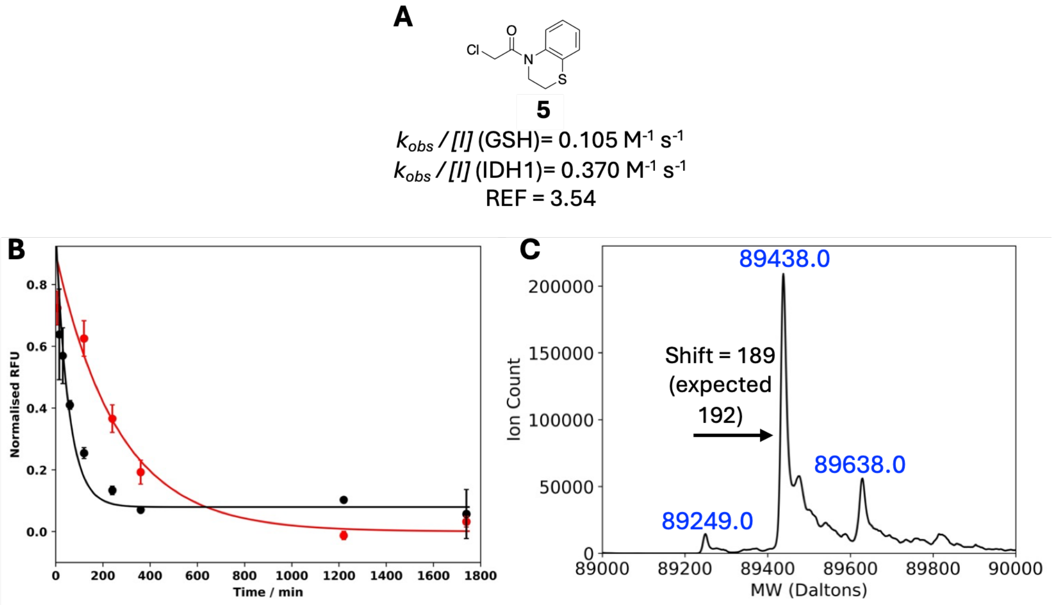


**Figure S6:** Triaging fragment **5**. (**A**) The chemical structure of fragment **5** and calculated rate constants for the reaction with MBP-IDH1$^{Cys-}$ (black) and GSH (red) (N = 3). (**B**) Triplicate qIT repeats show **5** reproducibly reacts at an enhanced rate with MBP-IDH1$^{Cys-}$ relative to with GSH (error = 1 standard deviation, N=3). (**C**) Intact-protein LC-MS analysis of MBP-IDH1$^{Cys-}$ (5 $\mu$M) following pre-incubation with fragment **5** (360 mins).

**A**

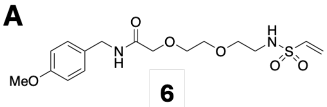


**6**

$k_{obs}$ / *[I]* (GSH) = 0.0189 $M^{-1}$ $s^{-1}$
$k_{obs}$ / *[I]* (IDH1) = 0.141 $M^{-1}$ $s^{-1}$
REF = 7.48

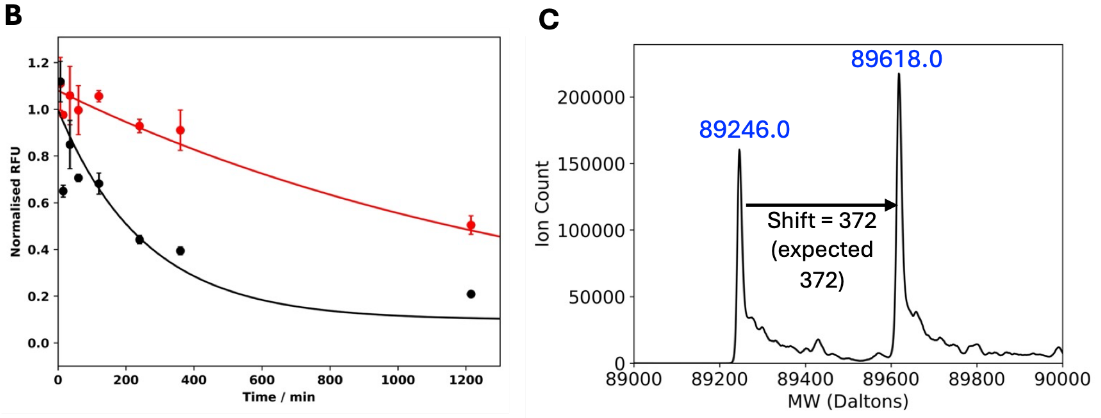


**Figure S7:** Triaging fragment **6**. (**A**) The chemical structure of fragment **6** and calculated rate constants for the reaction with MBP-IDH1$^{Cys-}$ (black) and GSH (red) (N = 3). (**B**) Triplicate qIT repeats show **6** reproducibly reacts at an enhanced rate with MBP-IDH1$^{Cys-}$ relative to with GSH (error = 1 standard deviation, N=3). (**C**) Intact-protein LC-MS analysis of MBP-IDH1$^{Cys-}$ (5 μM) following pre-incubation with fragment **6** (360 mins).

**A**

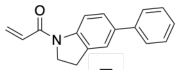


**7**

$k_{obs}$ / *[I]* (GSH) = 0.0331 $M^{-1}$ $s^{-1}$
$k_{obs}$ / *[I]* (IDH1) = 0.161 $M^{-1}$ $s^{-1}$
REF = 4.86

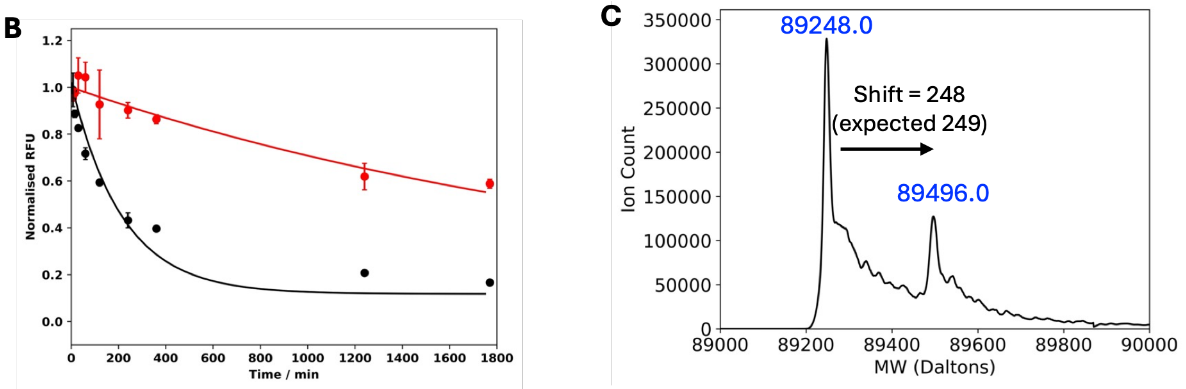


**Figure S8:** Triaging fragment **7**. (**A**) The chemical structure of fragment **5** and calculated rate constants for the reaction with MBP-IDH1$^{Cys-}$ (black) and GSH (red) (N = 3). (**B**) Triplicate qIT repeats show **7** reproducibly reacts at an enhanced rate with MBP-IDH1$^{Cys-}$ relative to with GSH (error = 1 standard deviation, N=3). (**C**) Intact-protein LC-MS analysis of MBP-IDH1$^{Cys-}$ (5 μM) following pre-incubation with fragment **7** (360 mins).

**A**

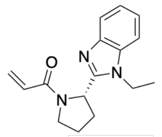


**8**

$k_{obs}$ / [I] (GSH) = 0.119 $M^{-1}$ $s^{-1}$

$k_{obs}$ / [I] (IDH1) = 0.562 $M^{-1}$ $s^{-1}$

REF = 4.72

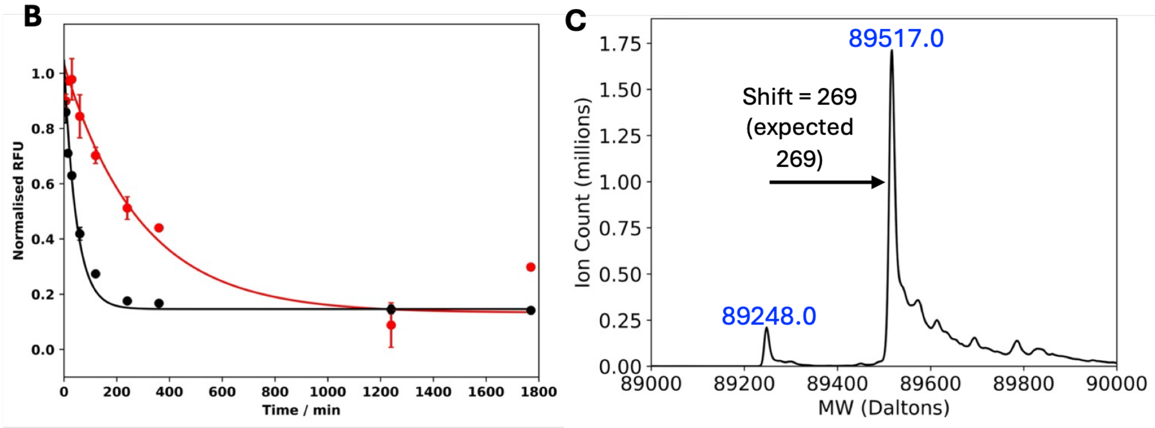


**Figure S9:** Triaging fragment **8**. (**A**) The chemical structure of fragment **8** and calculated rate constants for the reaction with MBP-IDH1$^{Cys-}$ (black) and GSH (red) (N = 3). (**B**) Triplicate qIT repeats show **8** reproducibly reacts at an enhanced rate with MBP-IDH1$^{Cys-}$ relative to with GSH (error = 1 standard deviation, N=3). (**C**) Intact-protein LC-MS analysis of MBP-IDH1$^{Cys-}$ (5 μM) following pre-incubation with fragment **8** (360 mins).

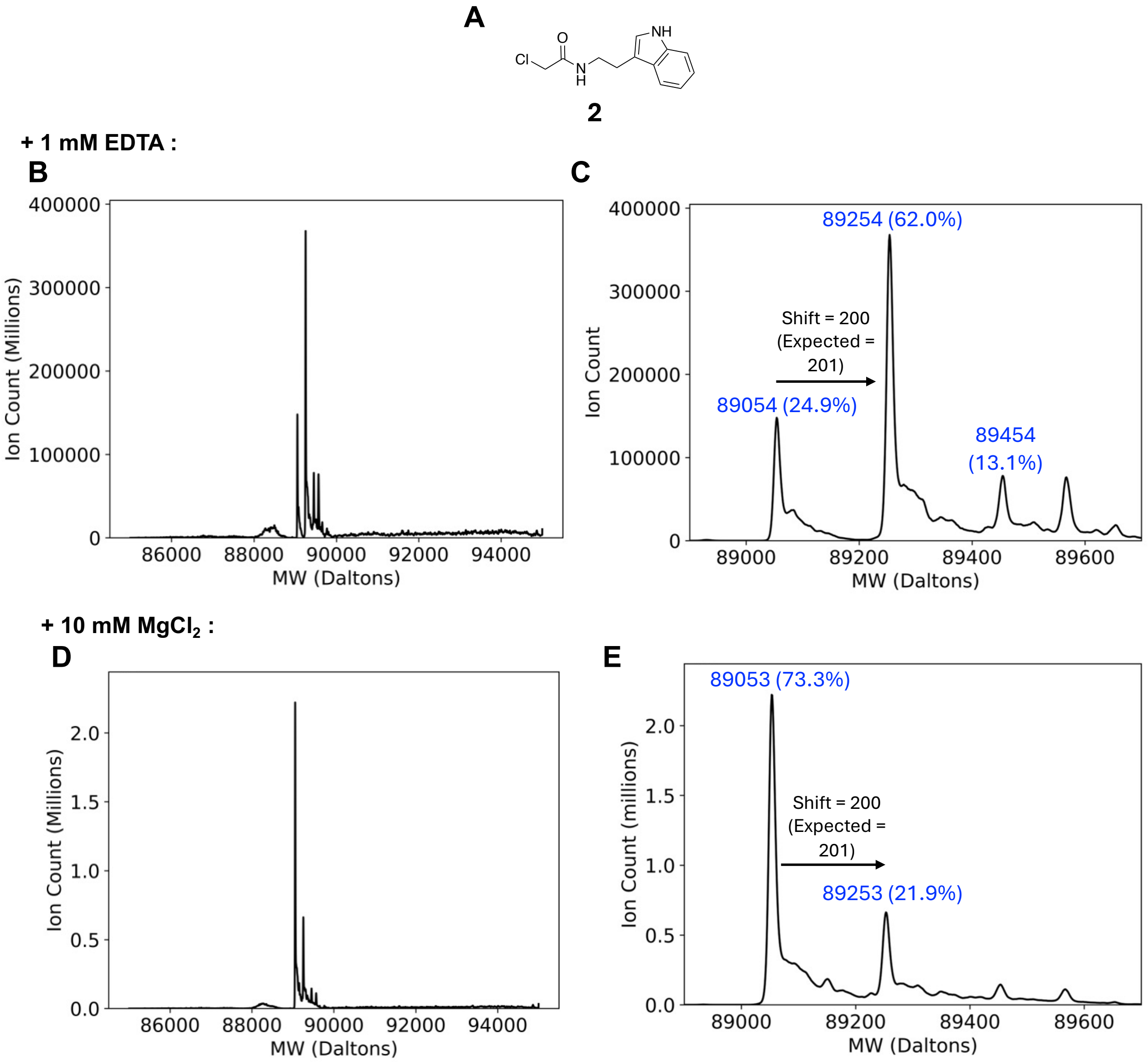


**Figure S10:** Deconvoluted spectra from intact-protein LC-MS analysis of MBP-IDH1 (5 µM) following 180 mins pre-incubation with fragment **2** (**A**, 500 µM) in buffer containing either 1 mM EDTA (**B/C**) or 10 mM $MgCl_2$ (**D/E**). The molecular weight and relative percentage abundance of each peak is displayed in blue.

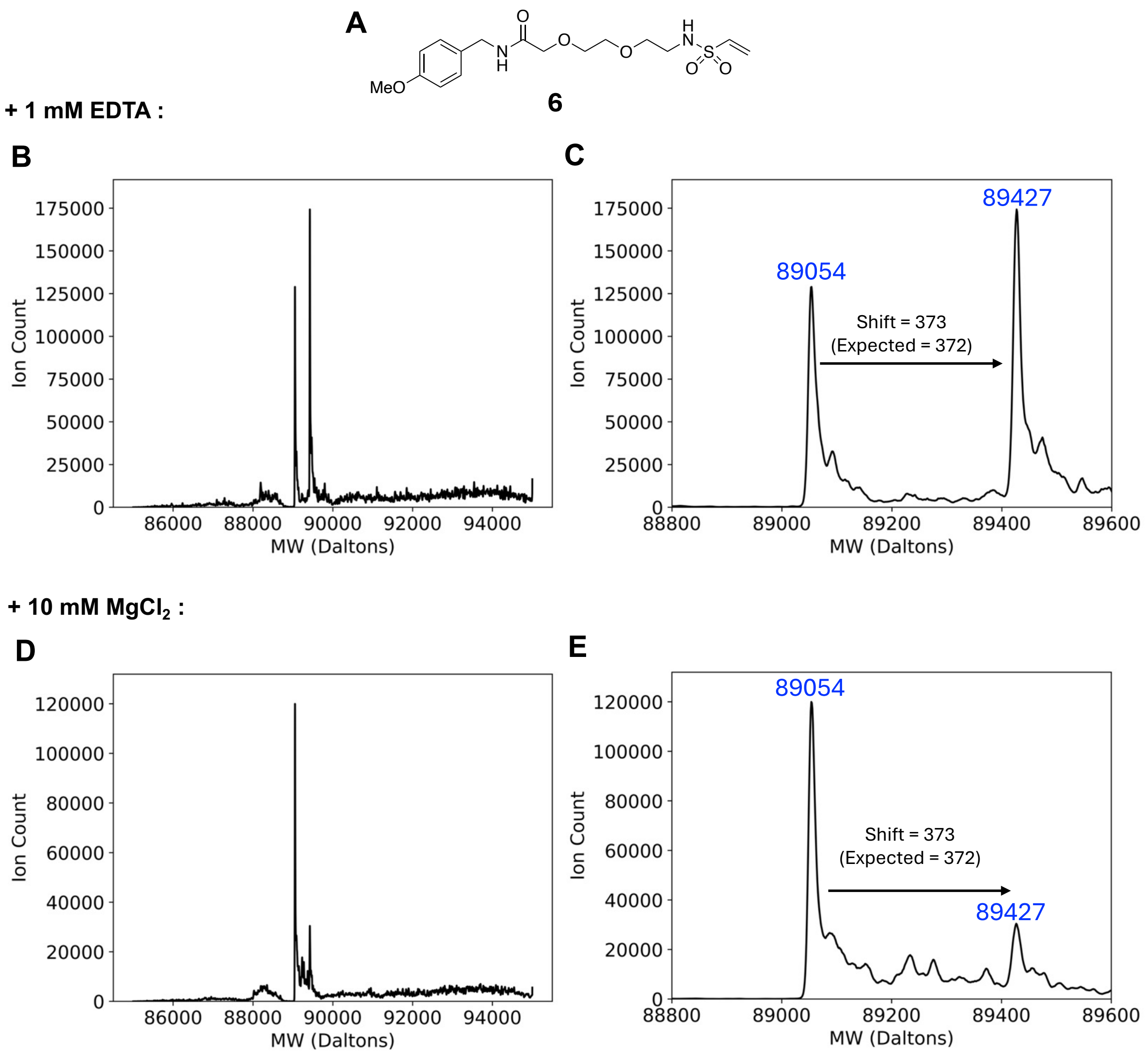


**Figure S11:** Deconvoluted spectra from intact-protein LC-MS analysis of MBP-IDH1 (5 μM) following 180 mins pre-incubation with fragment **6** (**A**, 500 μM) in buffer containing either 1 mM EDTA (**B/C**) or 10 mM $MgCl_2$ (**D/E**). The molecular weight and relative percentage abundance of each peak is displayed in blue.

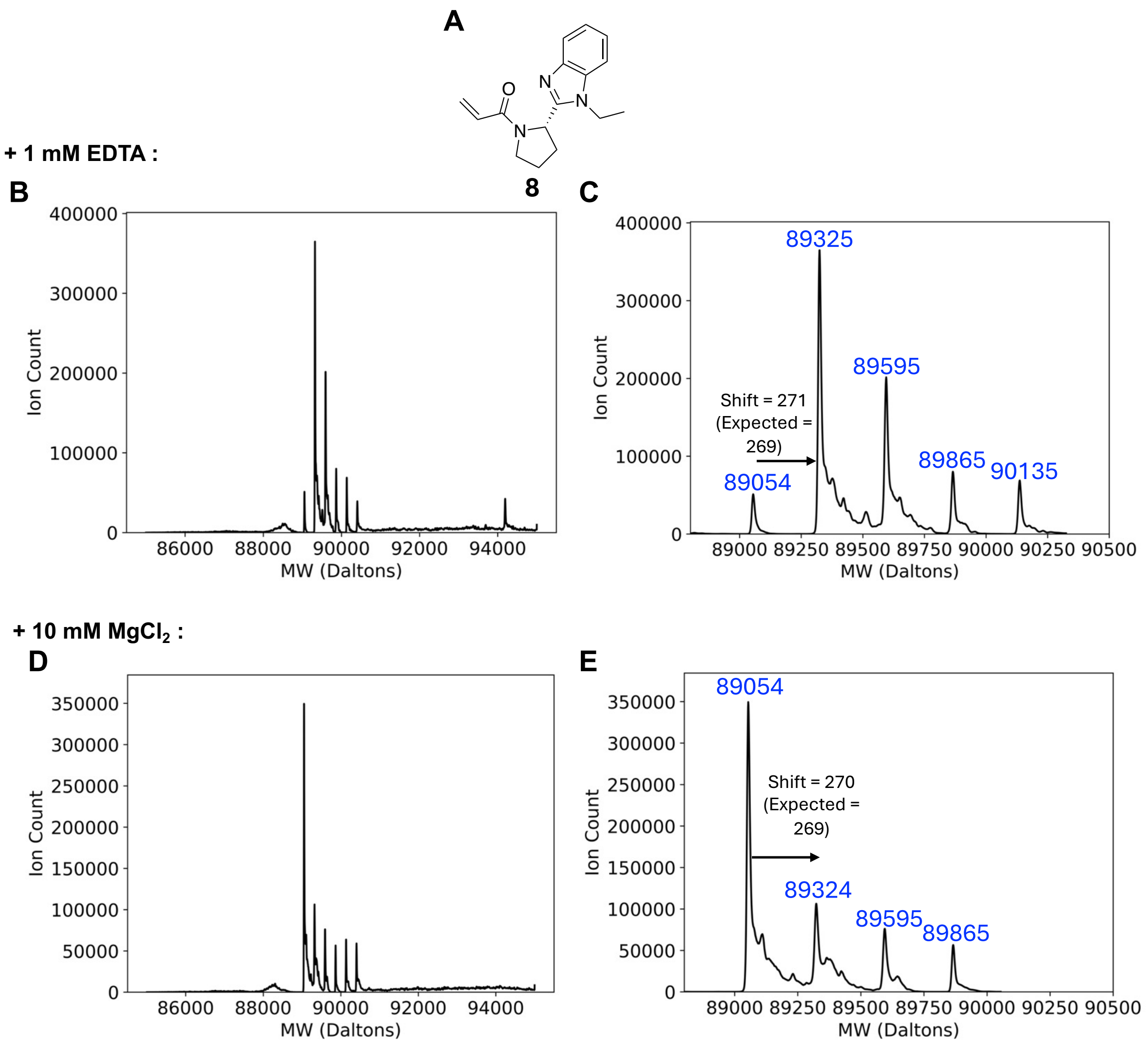


**Figure S12:** Deconvoluted spectra from intact-protein LC-MS analysis of MBP-IDH1 (5 μM) following 180 mins pre-incubation with fragment **8** (**A**, 500 μM) in buffer containing either 1 mM EDTA (**B/C**) or 10 mM $MgCl_2$ (**D/E**). The molecular weight and relative percentage abundance of each peak is displayed in blue.

**Figure S13 – Investigating Inhibition by 2 Through Conjugating One Side of the IDH1 Dimer**

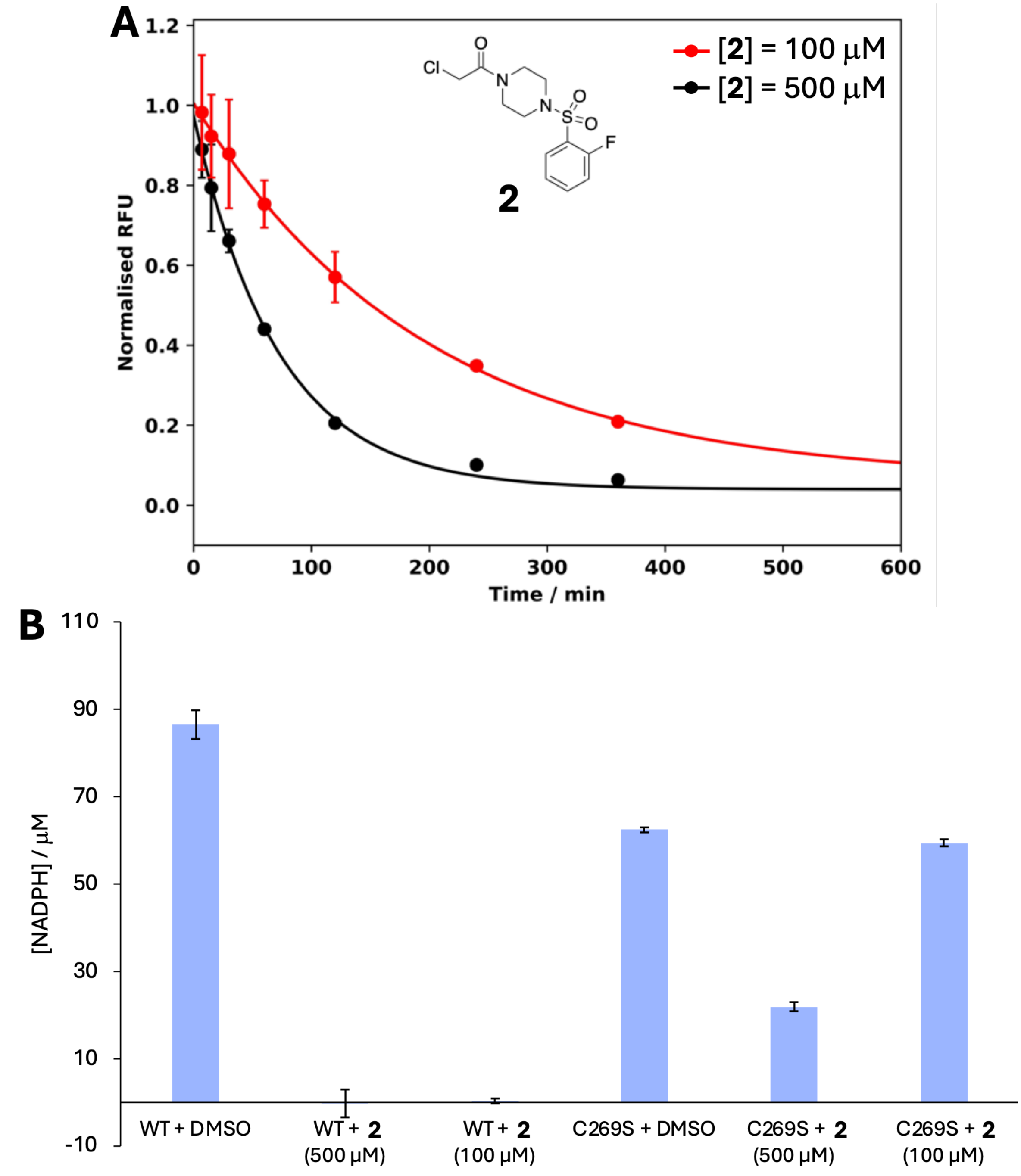


**Figure S13:** Evidence for conjugation of a single constituent of the IDH1 dimer inhibiting total activity. (**A**) qIT reaction curves for **2** when incubated at either 100 μM or 500 μM (N = 3). (**B**) Residual activity when either WT or C269S-mutant IDH1 is incubated with **2** for 2 h at a concentration of either 100 μM or 500 μM (N = 4).

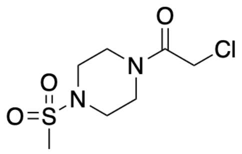


**18** (500 μM)

$k_{obs}$ / *[I]* (GSH) = 0.0535 $M^{-1}$ $s^{-1}$

$k_{obs}$ / *[I]* (IDH1) = 0.0509 $M^{-1}$ $s^{-1}$

REF = 0.95

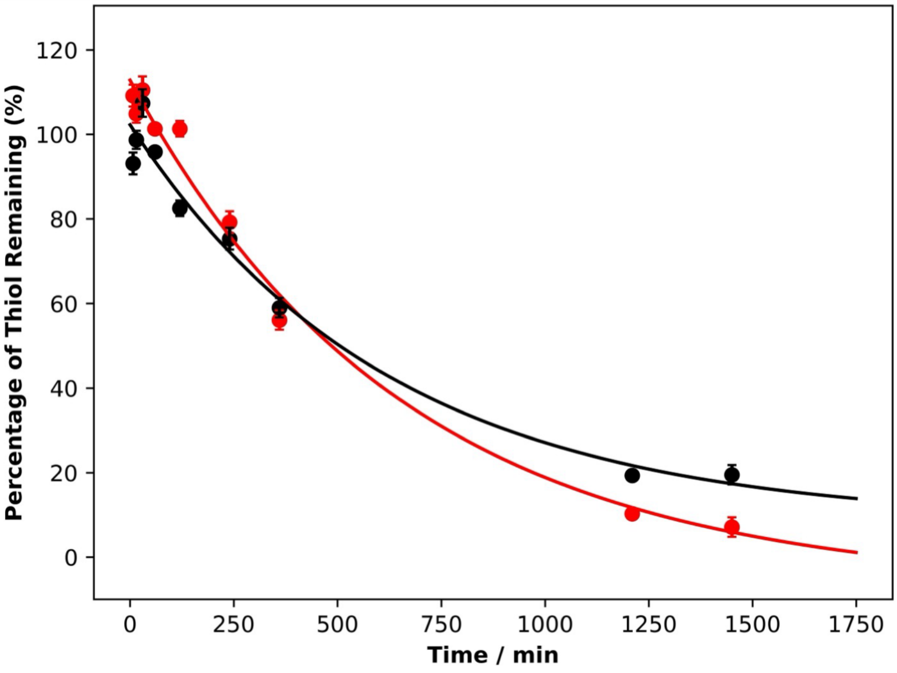


**Figure S14:** The chemical structure of fragment **18**, calculated rate constants for the reaction with MBP-IDH1$^{Cys-}$ and GSH, and triplicate qIT repeats measuring the reactions between **18** and MBP-IDH1$^{Cys-}$ (black) or GSH (red). Error = 1 standard deviation (N=3).

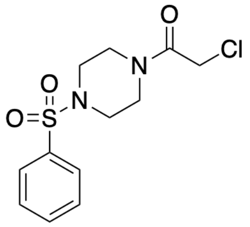


**10** (500 μM)

$k_{obs}$ / [I] (GSH) = 0.0474 $M^{-1}$ $s^{-1}$

$k_{obs}$ / [I] (IDH1) = 0.206 $M^{-1}$ $s^{-1}$

REF = 4.3

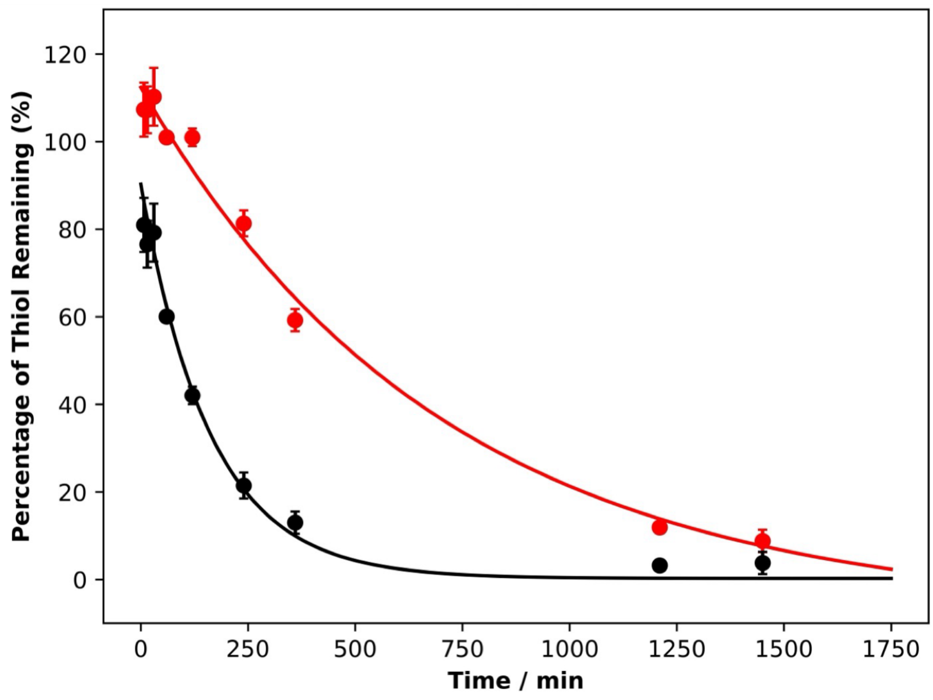


**Figure S15:** The chemical structure of fragment **10**, calculated rate constants for the reaction with MBP-IDH1$^{Cys-}$ and GSH, and triplicate qIT repeats measuring the reactions between **10** and MBP-IDH1$^{Cys-}$ (black) or GSH (red). Error = 1 standard deviation (N=3).

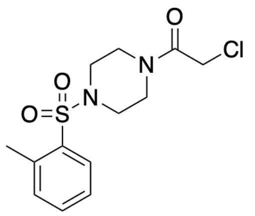


**11** (100 μM)
$k_{obs}$ / *[I]* (GSH) = 0.0937 $M^{-1}$ $s^{-1}$
$k_{obs}$ / *[I]* (IDH1) = 1.10 $M^{-1}$ $s^{-1}$
REF = 11.7

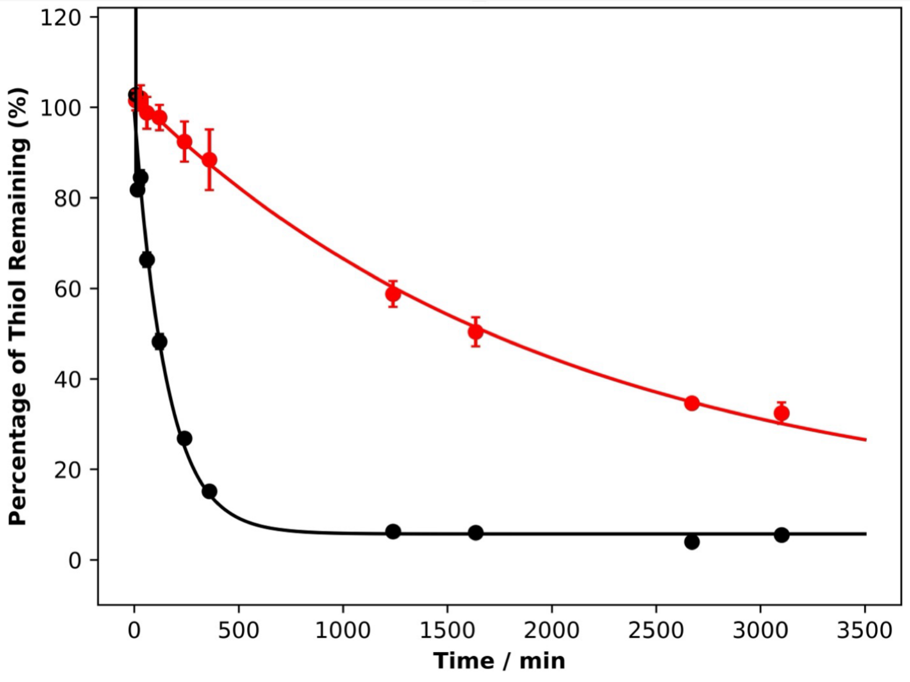


**Figure S16:** The chemical structure of fragment **11**, calculated rate constants for the reaction with MBP-IDH1$^{Cys-}$ and GSH, and triplicate qIT repeats measuring the reactions between **11** and MBP-IDH1$^{Cys-}$ (black) or GSH (red). Error = 1 standard deviation (N=3).

**A**

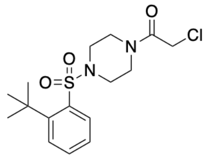


**12** (100 μM)
$k_{obs}$ / *[I]* (GSH) = 0.0791 $M^{-1}$ $s^{-1}$
$k_{obs}$ / *[I]* (IDH1) = 2.82 $M^{-1}$ $s^{-1}$
REF = 35.6

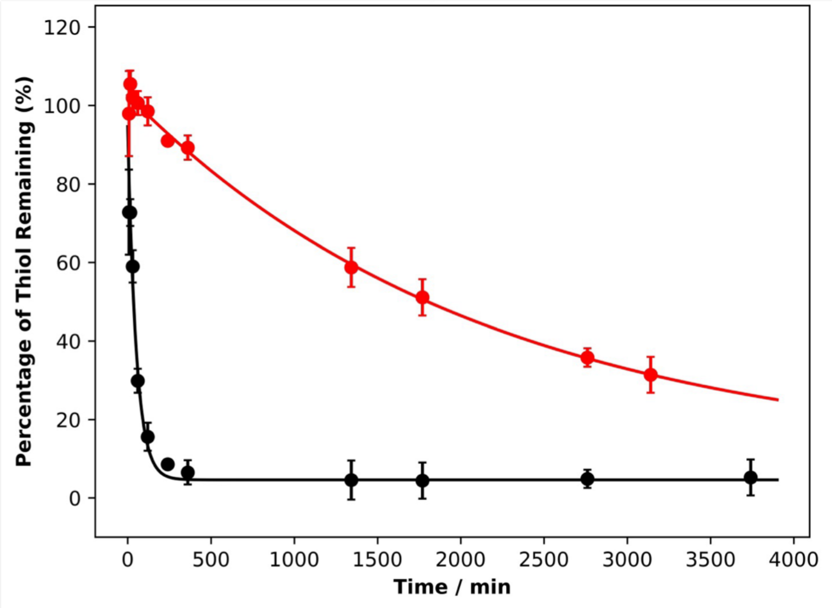


**Figure S17:** The chemical structure of fragment **12**, calculated rate constants for the reaction with MBP-IDH1$^{Cys-}$ and GSH, and triplicate qIT repeats measuring the reactions between **12** and MBP-IDH1$^{Cys-}$ (black) or GSH (red). Error = 1 standard deviation (N=3).

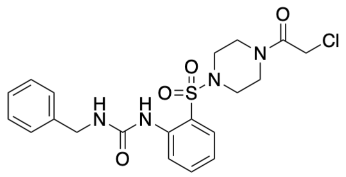

**13** (100 μM)

$k_{obs}$ / *[I]* (GSH) = 0.0679 $M^{-1}$ $s^{-1}$

$k_{obs}$ / *[I]* (IDH1) = 1.51 $M^{-1}$ $s^{-1}$

REF = 22.2

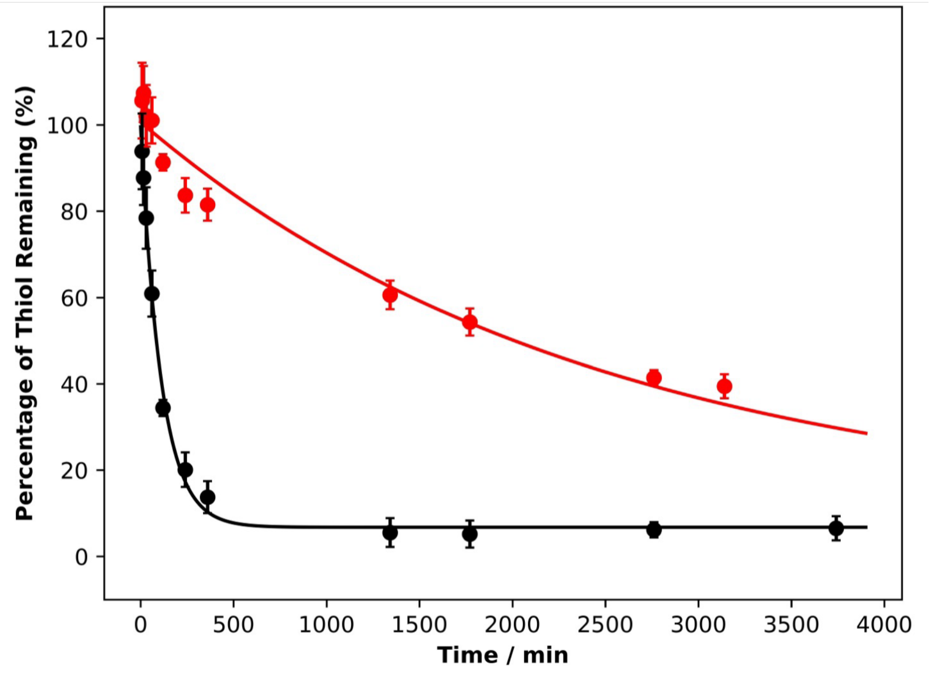


**Figure S18:** The chemical structure of fragment **13**, calculated rate constants for the reaction with MBP-IDH1$^{Cys-}$ and GSH, and triplicate qIT repeats measuring the reactions between **13** and MBP-IDH1$^{Cys-}$ (black) or GSH (red). Error = 1 standard deviation (N=3).

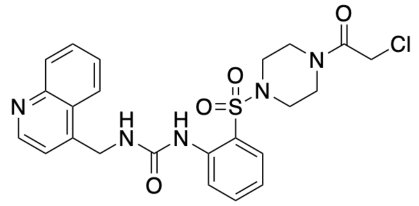


**14** (100 μM)

$k_{obs}$ / *[I]* (GSH) = 0.951 $M^{-1}$ $s^{-1}$

$k_{obs}$ / *[I]* (IDH1) = 0.0302 $M^{-1}$ $s^{-1}$

REF = 31.4

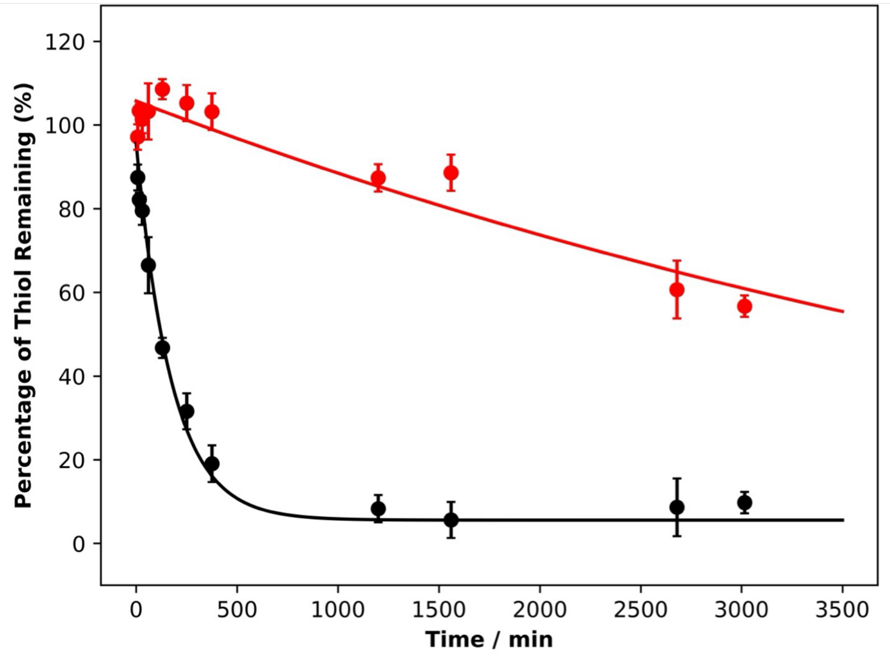


**Figure S19:** The chemical structure of fragment **14**, calculated rate constants for the reaction with MBP-IDH1$^{Cys-}$ and GSH, and triplicate qIT repeats measuring the reactions between **14** and MBP-IDH1$^{Cys-}$ (black) or GSH (red). Error = 1 standard deviation (N=3).

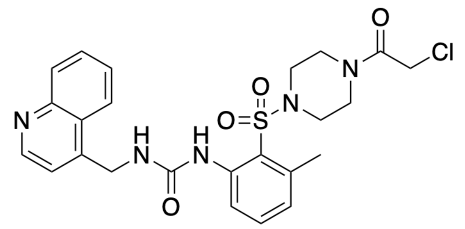

**15** (100 μM)

$k_{obs}$ / *[I]* (GSH) = 2.15 $M^{-1}$ $s^{-1}$

$k_{obs}$ / *[I]* (IDH1) = 0.0506 $M^{-1}$ $s^{-1}$

REF = 42.5

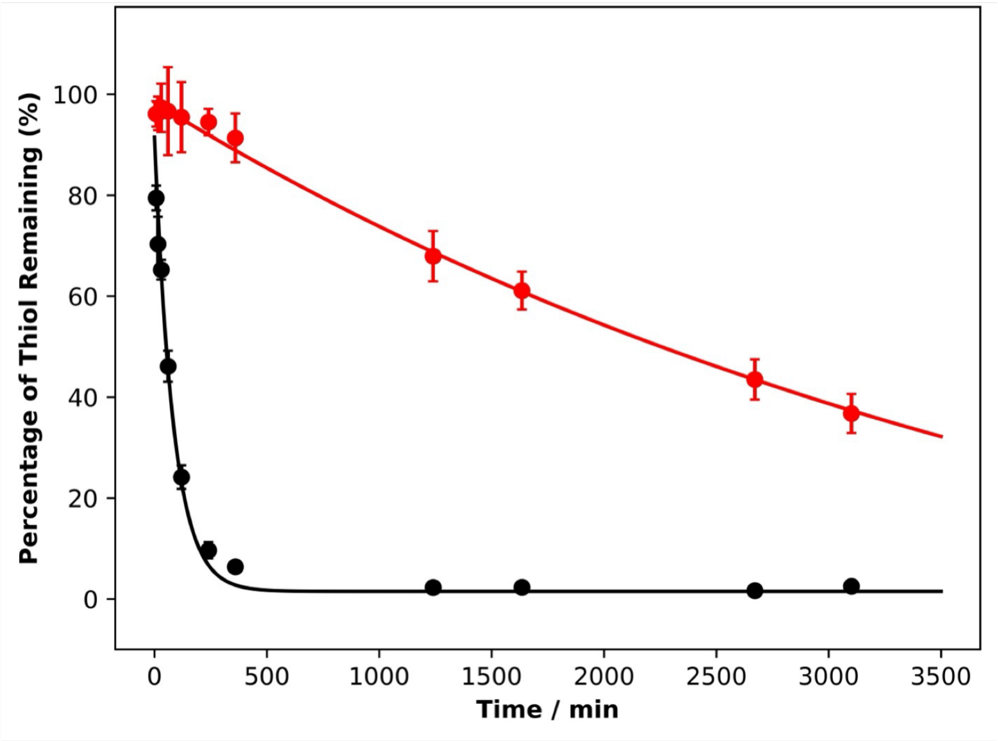


**Figure S20:** The chemical structure of fragment **15**, calculated rate constants for the reaction with MBP-IDH1$^{Cys-}$ and GSH, and triplicate qIT repeats measuring the reactions between **15** and MBP-IDH1$^{Cys-}$ (black) or GSH (red). Error = 1 standard deviation (N=3).

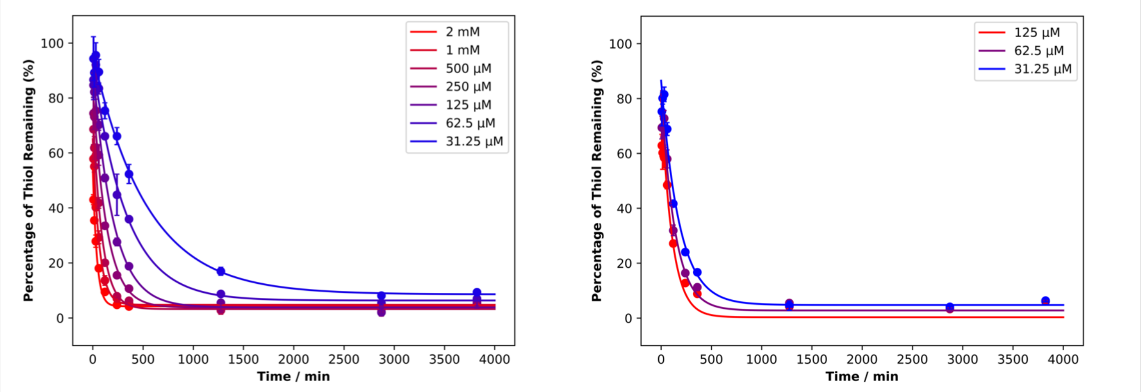


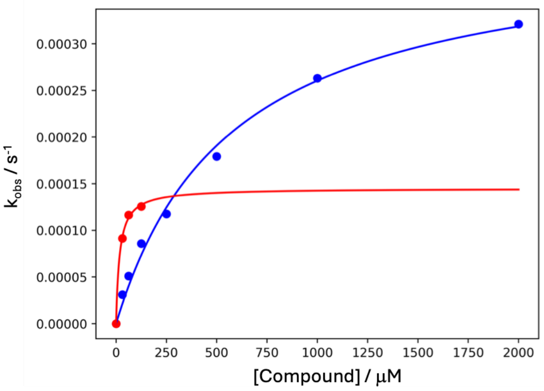


D

| Compound | $k_{inact}$ ($s^{-1}$) | $K_I$ (µM) | $k_{inact}$ / $K_I$ ($M^{-1}$ $s^{-1}$) |
|---|---|---|---|
| **2** | 0.000410 | 573 | 0.715 |
| **15** | 0.000145 | 17.5 | 8.27 |

**Figure S21:** $k_{inact}/K_I$ determination for compounds **2** and **15**: (**A**) qIT profiling measuring the rate of reaction between **2** and MBP-IDH1$^{Cys-}$ at increasing concentrations (error bars = 1 standard deviation, N = 3). (**B**) qIT profiling measuring the rate of reaction between **15** and MBP-IDH1$^{Cys-}$ at increasing concentrations (error bars = 1 standard deviation, N = 3). (**C**) A comparison between the rate of IDH1-**15** and IDH1-**2** reactions at increasing concentrations. (**D**) Estimated $k_{inact}$, $K_I$ and $k_{inact}/K_I$ values for **2** and **15**.

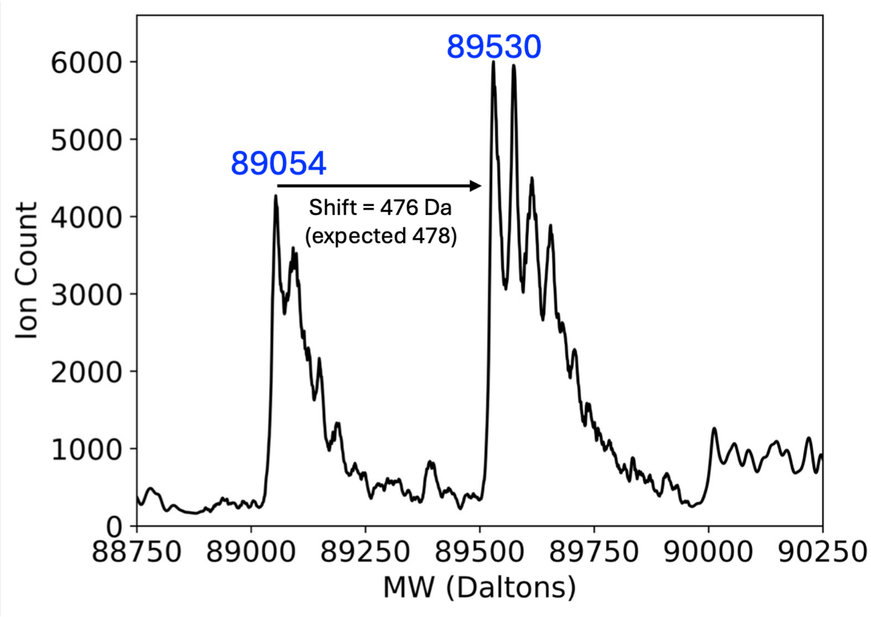


**Figure S22:** Deconvoluted spectra from intact-protein LC-MS analysis of MBP-IDH1 (1 μM) following incubation with fragment **17** (2 μM, 120 mins)**.**

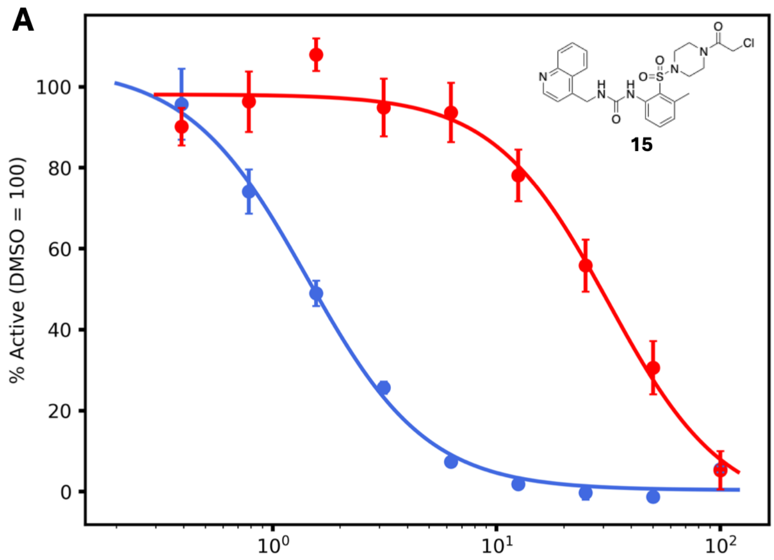


| Protein | IDH1 | IDH2 |
|---|---|---|
| $IC_{50}$ (μM) | 1.35 | 23.7 |

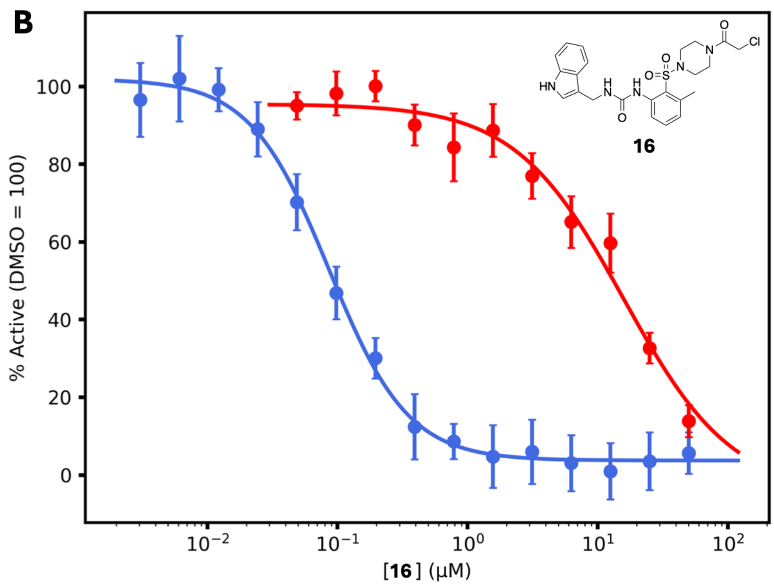


| Protein | IDH1 | IDH2 |
|---|---|---|
| $IC_{50}$ (μM) | 0.0863 | 15.9 |

**Figure S23:** Activity assay results showing inhibitory potency of **15** and **16** against IDH1 and IDH2.

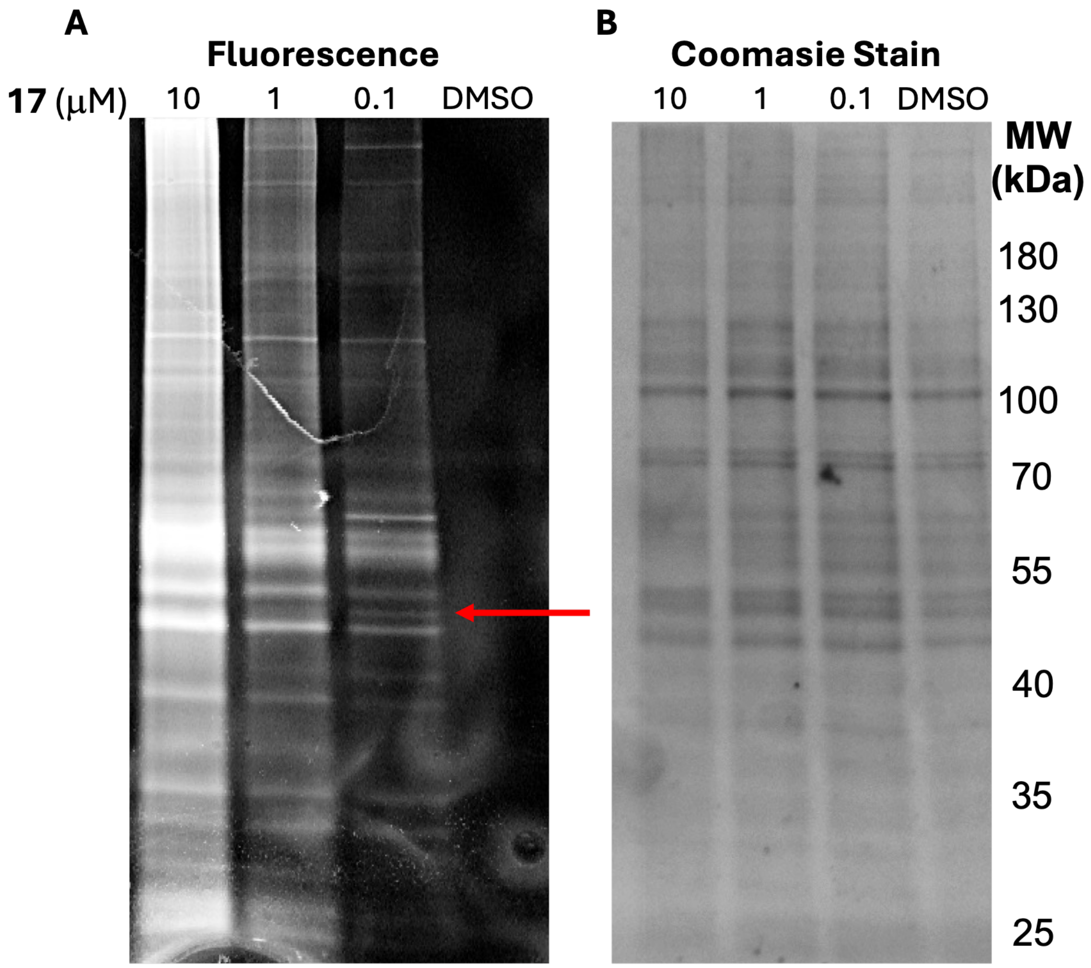


**Figure S24:** Gel-based chemoproteomic analysis of HEK293 cell lysates treated with **17** (at increasing concentrations) or DMSO (1% *v/v*), followed by conjugation of labelled proteins to TAMRA-azide. (**A**) Labelled bands as visualised by fluorescence imaging (λex / λabs = 532 / 575 nm), with the putative IDH1 band labelled by a red arrow. (**B**) Visualisation of total protein content via Coomasie blue staining.

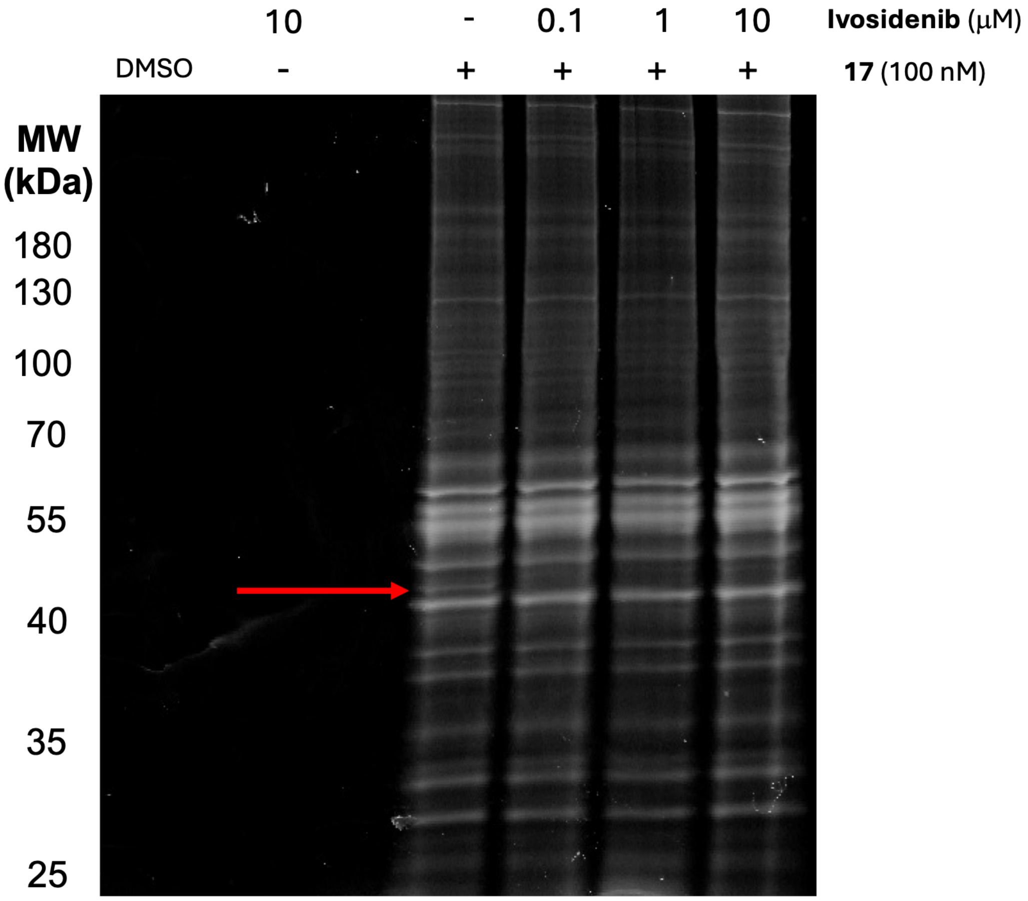


**Figure S25:** Gel-based chemoproteomic competition assay between **17** and increasing concentrations of ivosidenib, showing the results of incubation of relevant compounds with HEK293 cell lysates, followed by conjugation of labelled proteins to TAMRA-azide. The putative IDH1 band labelled by a red arrow.

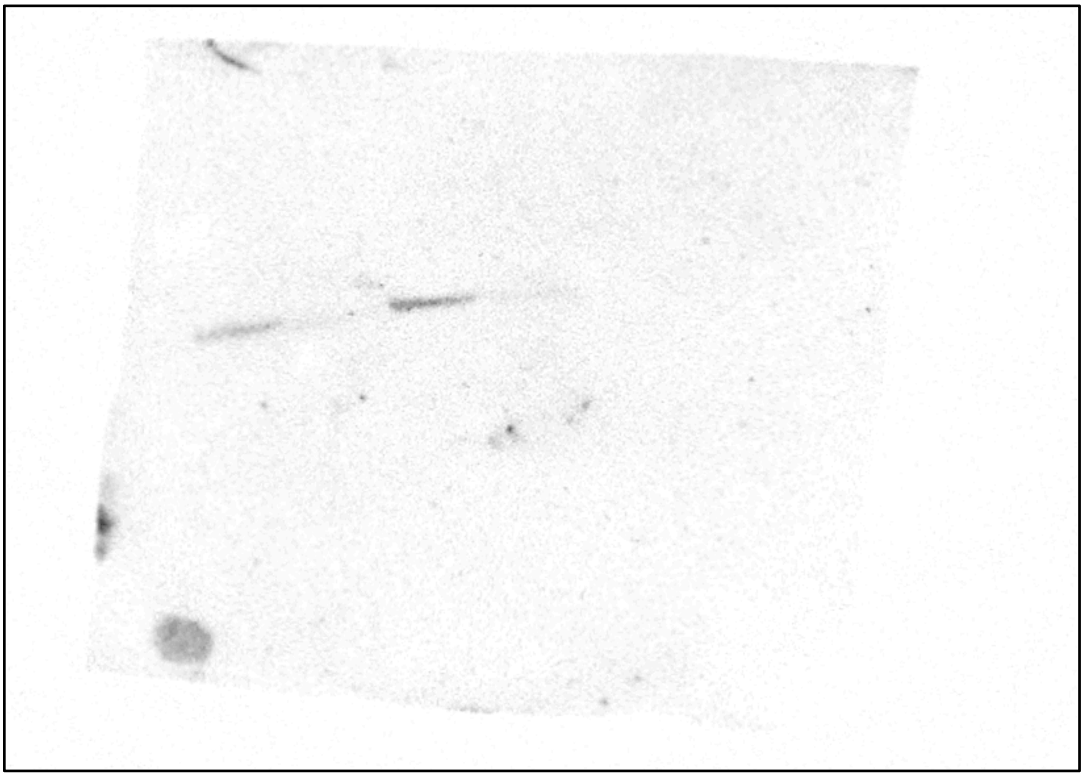

**Figure S26:** Uncropped western-blot visualisation of biotin-streptavidin pulldown by **17** in HEK293-lysates (cropped version in Fig. 5E).

### 2. Materials and Methods

#### 2.1 Biological Materials and Methods

##### *2.1.1 General Biological Materials*

**Media**

Terrific broth (TB): tryptone (12 g/L) yeast extract (25 g/L), glycerol (4 mL), $KH_2PO_4$ (0.017 M), $K_2HPO_4$ (0.072 M).

Luria-Bertani broth (LB): tryptone (10 g/L), yeast extract (5 g/L), NaCl (10 g/L).

**Buffers**

- Lysis buffer: NaCl (500 mM), Tris-HCl pH 8 (50 mM), DTT (1 mM), DNase (5 $\mu$g / mL).
- Wash buffer: NaCl (150 mM), Tris-HCl pH 8 (50 mM), DTT (1 mM).
- Elution buffer: NaCl (150 mM), Tris-HCl pH 8 (50 mM), DTT (1 mM), Maltose (10 mM)
- qIT assay buffer: NaCl (150 mM), Hepes pH 8 (25 mM), $MgCl_2$ (10 mM).
- qIT quench buffer: NaCl (150 mM), Hepes pH 7.4 (25 mM), $MgCl_2$ (10 mM).
- Activity assay buffer 1: NaCl (150 mM), Hepes pH 8 (50 mM), DTT (100 $\mu$M).
- Activity assay buffer 2: NaCl (150 mM), Hepes pH 8 (50 mM), DTT (100 $\mu$M), $MgCl_2$ (20 mM), isocitrate (200 $\mu$M), $NADP^+$ (500 $\mu$M).
- CD buffer: phosphate pH 7.4 (20 mM), DTT (1 mM)

##### *2.1.2 Computational Modelling of IDH1-Substrate and Compound Equilibria*

To estimate IDH1 inhibitor occupancy in the absence and presence of substrates, a model was built using the systems biology modelling software package (Copasi).[3] Isocitrate-$Mg^{2+}$ was treated as a single substrate (consistent with past findings that isocitrate-$Mg^{2+}$ typically binds as a single substrate)[4] at a concentration of 100 $\mu$M (previous work has estimated cellular isocitrate concentrations to be in the double-digit mM range and $Mg^{2+}$ concentrations to be much higher)[5-7], with a binding affinity ($K_D$) for IDH1 of 0.9 $\mu$M (previously reported by *Liu et al)*[4], whilst the IDH1 concentration was estimated at 10 nM (this was kept low to model a large excess of substrate and inhibitor). Reversible inhibitors were modelled with $K_D$ values of 1 and 10 nM, whilst an irreversible inhibitor was modelled with a $k_{inact}/K_I$ value of $1 \times 10^6$ $M^{-1}s^{-1}$. All reversible steps were modelled assuming fast exchange kinetics.

##### *2.1.3 MBP-IDH1 Transformation, Expression and Purification*

(Note: this procedure was used for the expression and purification of all MBP-IDH1 constructs and MBP-IDH2)

***Transformation:*** Aliquots (50 μL) of electrocompetent *E. coli* BL21 (DE3) cells were thawed on ice for 15 min, then MBP-IDH1-containing pET-21A plasmid DNA (1 μL) was added, and the mixture was subjected to electroporation (resistance = 200 Ohms, Capacitance (ex) = 250 uFD, Gene pulser = 2.2, Capacitance = 25 uFD). LB media (1 mL) was immediately added, and the mixture spread over LB agar containing ampicillin (10 μg / mL), which was incubated overnight at 37˚C.

***Recombinant Expression and Purification:*** Approximately 50 colonies of *E. coli* BL21 (DE3) pre-transformed with MBP-IDH1-encoding plasmids were transferred into 2 L of terrific broth media containing ampicillin (50 μg / mL) and allowed to grow under shaking (200 RPM) at 37°C until $OD_{600}$ reached 0.7 – 0.9. The culture was then cooled to 20°C for 60 mins, before recombinant MBP-IDH1 expression was induced via the addition of IPTG (0.5 mM) and the culture subsequently left shaking (200 RPM) at 20°C overnight. Cells were then pelleted via centrifugation (10 min, 6,000 RPM) and used for MBP-IDH1 purification.

Harvested cells were suspended in ice cold lysis buffer via agitation using a vortex, then passed through a 50 mL syringe to ensure complete suspension. Cells were then lysed using a cell disrupter (*Constant systems*) at a pressure of 27 kpsi, and the lysate pelleted via centrifugation (18,000 RPM, 45 mins, 4°C). The resulting supernatant was passed through an MBPTrapHP column (*Cytiva*) at 2.5 mL / min, which was subsequently washed with wash buffer (10 column volumes, 5 mL / min) and bound proteins eluted using elution buffer (2 column volumes, 2.5 mL / min). As lysates typically overwhelmed the MBP-binding capacity of the column, MBPTrap flowthroughs were reapplied to the column, washed and bound proteins eluted 2-3 times.

Fractions containing MBP-IDH1 (as judged using a nanodrop) were pooled, concentrated using centrifugal spin filters (*Amicron*) and further purified via size-exclusion chromatography. An AKTA pure (*General Electric*) protein purification system with a 200 pg SEC column was pre-equilibrated with wash buffer, then concentrated MBP-IDH1 containing solution (2 mL) was applied to the column, and proteins eluted at 1 mL / min. Elutions were assessed for MBP-IDH1 purity via SDS-PAGE gel electrophoresis. Fractions containing purified MBP-IDH1 were pooled, concentrated to ~10 mg / mL using a centrifugal spin filter, flash frozen on dry ice and stored at -80°C until required.

##### 2.1.4 qIT Assay Procedure

***Assay setup:*** MBP-IDH1$^{Cys-}$ aliquots were thawed on ice, then thoroughly exchanged into qIT assay buffer until [DTT] < 0.5 μM and diluted to a final protein concentration of 15 μM. A GSH solution was simultaneously prepared by diluting GSH in qIT assay buffer to a concentration of 15 μM. Pierce's immobilized TCEP disulfide reducing gel (*Thermo Scientific*) was thoroughly washed with qIT assay buffer, and diluted to make a 1.5% *v/v* solution. In a black 384-well flat bottom assay plate (*Corning*, product number 3575) 20 μL of TCEP-gel solution was combined with either 20 μL of MBP-IDH1$^{Cys-}$ solution (termed the *Reaction Plate*), 20 μL of GSH solution (termed the *GSH Plate)* or buffer alone (in negative control columns) and incubated at 23°C for 60 mins. Fragments (50 mM stock solutions in DMSO, or DMSO alone for control wells) were thawed, then diluted to a concentration of 1.5 mM in qIT assay buffer. The assay was started by adding 20 μL of fragment solution to the relevant wells in the protein and GSH assay plates using a liquid handling robot (*Analytik Jena*), then was incubated at 23°C and the proportion of thiol remaining in each plate quantified via quenching into CPM at regular time points. The final assay plate – termed the *Reaction plate* - included either MBP-IDH1 or GSH (5 μM), compounds (500 μM) and TCEP-reducing gel (0.5% *v/v*) in qIT assay buffer.

***CPM Quenches:*** At regular time points (typically 7, 15, 30, 60, 120, 240, 360, ~1200 and ~1600 mins) a solution of CPM (1.39 μM) in qIT quench buffer was prepared, and 27 μL was dispensed into each well of a 384-well flat bottom assay plate – termed a *Quench plate*. Using the liquid handling robot, a 3 μL sample was removed from each well of the protein and GSH *Reaction plates* and dispensed into the *Quench plate,* which was subsequently mixed via 3 x 20 μL aspiration and dispension operations. This was incubated at 23°C for 60 mins, then fluorescence (excited at 384 nm and read at 470 nm) measured for each well using a Clariostar Plus Microplate Reader (*BMG Labtech*).

***Data analysis:*** Using a custom-built python-script, fluorescence measurements for each fragment-containing well were normalised, then a one-phase exponential decay curve was fitted to the data via non-linear regression using the *optimize.curve_fit* function from the *Scipy* module. REF values were determined for each compound by dividing the rate constant for the reaction with MBP-IDH1$^{Cys-}$ by the rate constant for the reaction with GSH, and were used to select and rank hits.

##### 2.1.5 Intact Protein LC-MS

***Sample preparation:*** Protein was diluted in qIT assay buffer to a final concentration of 5 μM, then DTT (100 μM) added and the sample incubated at 23°C for 60 mins. Fragments were added to the desired concentration (either 1 – 500 μM depending on the experiment) in protein solution so that the final DMSO concentration reached 1% *v/v*, then the reaction was incubated for the desired period of time (as described with each result). Samples were desalted and excess fragment removed via sequential rounds of buffer exchange into mass spectrometry buffer (5 x 1 in 5 dilutions) into 10 mM ammonium bicarbonate solution (in LC-MS grade water), then diluted to a final concentration of 1 μM, flash frozen, and stored at -80°C.

***Data collection:*** A 6545XT AdvanceBio LC/QTOF (*Agilent*), coupled to an Aquity UPLC 2.1 mm x 50 mm, 2.7 μm protein BEH C4 column (*Waters*) were used for these experiments, with gradients of buffer A (water + 0.1% formic acid) to buffer B (acetonitrile + 0.1% formic acid) used to elute proteins bound to the column. A flow rate of 0.2 ml / min was maintained for all experiments. All spectra were obtained using electrospray ionisation in the positive ion mode (see **Table 16** for ionisation parameters) in positive ion mode, acquiring spectra in the 500 – 2500 *m/z* range. The samples were kept at 4°C and column at 40°C during experiments. The column was preequilibrated with 96.3% buffer A, then 1 μL of sample injected. The column was washed with buffer A for 1 min, then bound material was eluted by a linear gradient to 96.3% buffer B over the course of 4 minutes, then the column held at 96.3% buffer B for a further 1 minute. Resulting mass spectra were deconvoluted using the Masshunter software package (*Agilent*) on maximum entropy mode, and plotted using a custom-built python script.

| **Ionisation Parameters:** | |
|---|---|
| Drying gas temperature (mL / min) | 290 |
| Drying gas flow rate (L / min) | 14 |
| Nebuliser pressure (PSI) | 30 |
| Sheath gas temperature (°C) | 400 |

**Table S1:** LC-MS Ionisation Parameters.

| | |
|---|---|
| Sheath gas flow rate (L / min) | 12 |
| Capillary voltage (V) | 5000 |
| Nozzle voltage (V) | 2000 |

##### *2.1.6 Activity Assays*

***Activity Assay General Procedure***: MBP-IDH1 (or relevant mutants) were diluted to a concentration of 20 nM in *activity assay buffer 1,* then 20 μL aliquoted in triplicate into the wells of a low volume 384-well black, clear flat bottom microplate (*Corning*, product number = 3540). Simultaneously, *activity assay buffer 2* was aliquoted into the wells of a 384-well transfer plate, before 20 μL was transferred from each well into the protein-containing plate using a liquid handling robot, so that substrates were introduced to all the MBP-IDH1-containing wells at the same time. The assay plate was next centrifuged at 500 RPM for 15 seconds, then absorbance read at 340 nm continuously for 60 mins at 23°C.

***MBP-IDH1 Mutant Comparison:*** MBP-IDH1, MBP-IDH1$^{C269W}$ and MBP-IDH1$^{C269S}$ were each diluted in *activity assay buffer 1* to a final concentration of 20 nM, then MBP-IDH1 and MBP-IDH1$^{C269S}$ were both incubated with either **2** (at a final concentration of 500 or 100 μM, with DMSO at 1% *v/v*) or DMSO (1% *v/v*), whilst MBP-IDH1$^{C269W}$ was incubated with DMSO alone (1% *v/v*), at 23°C for 2 h. Remaining activity was assessed using the *Activity Assay General Procedure.*

***$IC_{50}$ Determination:*** MBP-IDH1 was diluted to a concentration of 20 nM in *activity assay buffer 1*, then preincubated with compound at halving concentrations (1% *v/v* DMSO) at 23°C for 2 h. Remaining activity was assessed using the *Activity Assay General Procedure.*

##### *2.1.7 Fragment Labelled IDH1 Preparation Procedure*

***Fragment preincubation:*** MBP-IDH1 was thoroughly exchanged into qIT assay buffer (without $MgCl_2$) via sequential rounds of buffer exchange using centrifugal spin filters (*Amicron*) and diluted to a final concentration of 5 μM. Immobilized TCEP disulfide reducing gel was washed with qIT assay buffer (with $MgCl_2$ removed), then added to the MBP-IDH1 solution to a final concentration of 0.5% *v/v* and incubated at 23°C for 60 mins. The relevant fragment was added (at either 500 or 100 μM with 1% *v/v* DMSO) and incubated at 23°C for the period of time indicated in the results section. Excess compound was next removed via sequential rounds of buffer exchange using centrifugal spin filters (*Amicron*) into qIT assay buffer (without $MgCl_2$, but with the addition of 10 mM DTT). GST-precision protease was added in a 1:20 (protease:MBP-IDH1) ratio, and incubated at 4°C overnight.

***Fragment Labelled IDH1 Purification:*** GST (*Pierce*) and MBP (*Abcam*)-affinity resins (100 μL of each / 10 mg of MBP-IDH1) were washed with *wash buffer* via sequential rounds of dilution (10 mL) and centrifugation (5000 RPM, 1 min), then were incubated with the cleavage reaction mixture at 4°C for 2 h. Resin was then pelleted via centrifugation (5000 RPM, 1 min) and the supernatant further purified via size-exclusion chromatography. An AKTA pure (*General Electric*) protein purification system with a 200 pg SEC column was pre-equilibrated with wash buffer, then IDH1 containing solution (2 mL) was applied to the column, and proteins eluted at 1 mL / min. Elutions were assessed for IDH1 purity via SDS-PAGE gel electrophoresis. Fractions containing purified IDH1 were pooled, concentrated to ~10 mg / ml using a centrifugal spin filter, then either used immediately or flash frozen on dry ice and stored at -80°C until required.

##### *2.1.8 Circular Dichroism*

**Circular dichroism data collection:** Fragment-labelled IDH1 (obtained using the *Fragment Labelled IDH1 Preparation Procedure*) or apo-IDH1 (obtained using the *Fragment Labelled IDH1 Preparation Procedure* but with the fragment pre-incubation step removed) were diluted in CD buffer to a concentration of 10 μM. For substrate-bound experiments, additional $Ca^{2+}$ (1 mM) and isocitrate (250 μM) were added, otherwise EDTA (1 mM) was added. IDH1 solution (300 μL) was placed in a 0.1 cm path length quartz cuvette (*Hellma*), then ellipticity measured every 0.5 nm at wavelengths from 190-280 nm. For temperature dependent measurements, spectra were obtained every 5°C at increasing temperatures from 20 - 95°C. All measurements were performed using a Chirascan spectrophotometer (*Applied Photophysics*). The melting temperature was determined by plotting the average ellipticity between 215 - 220 nm against temperature, then fitting to a sigmoidal curve using a custom-built python script. In each case, the melting temperature was determined from three repeats.

##### *2.1.9 Assessing Target-Engagement via Gel-Based ABPP*

HEK293 cell lysates (1 mg / ml) were incubated with **17** (at concentrations of 0.1 - 10 μM as indicated in relevant figures, in 1% *v/v* DMSO) or a DMSO control (1% *v/v*), with or without ivosidenib (at concentrations of 0.1 – 10 μM) at room temperature for 2h. Proteins were then precipitated via methanol-chloroform extraction and resolubilised in PBS + 1% SDS. CuAAC was performed via sequential addition of TAMRA-Azide (100 μM), THTBA (500 μM), $CuSO_4$ (1 mM) and TCEP (1 mM), followed by incubation for 60 mins at RT, and was quenched via EDTA (5 mM) addition. Proteins were again purified via methanol-chloroform extraction, then suspended in 2 x SDS-PAGE running buffer (50 μL), boiled at 95°C for 15 mins, and separated on a 10% polyacrylamide gel. Fluorescent bands were visualised FUJIFILM FLA-5000 fluorescence imaging system (λex/λabs = 532/575 nm), then gels stained with Coomasie brilliant blue, and total protein content imaged.

##### *2.1.10 Assessing Target-Engagement via Pulldown*

**Probe treatment and pulldown:** HEK293 cell lysates (1 mg / ml) were incubated with **17** (1 μM, in 1% *v/v* DMSO) or a DMSO control (1% *v/v*), with or without ivosidenib (1 μM) at room temperature for 2h. Proteins were then precipitated via methanol-chloroform extraction and resolubilised in PBS + 1% SDS. CuAAC was performed via sequential addition of Biotin-$[PEG]_3$-Azide (100 μM), THTBA (500 μM), $CuSO_4$ (1 mM) and TCEP (1 mM), followed by incubation for 60 mins at RT, and was quenched via EDTA (5 mM) addition. Proteins were again purified via methanol-chloroform extraction, then suspended in PBS + 0.25% SDS and incubated at 4°C overnight under shaking with 7.5 μL of Pierce Streptavidin Magnetic Beads (*Thermo Scientific*). Magnetic beads were washed 4 x with PBS + 2% SDS, then suspended in 5 x SDS-PAGE running buffer (50 μL) and boiled at 95°C for 15 mins. Magnetic beads were subsequently pelleted, and eluted proteins separated on a 10% polyacrylamide gel.

**Immunoblotting:** Proteins were transferred from a polyacrylamide gel to a nitrocellulose membrane in transfer buffer (25 mM tris, 192 mM glycine, 20% *v/v* methanol) via a wet transfer procedure (400 mA, 60 mins), then incubated under rocking overnight at 4°C with an anti-IDH1 antibody (*Thermo Fisher*, GT1521) diluted 1 in 10,000 in 5% v/v skimmed milk solution. The nitrocellulose membrane was subsequently washed with 30 mL of PBS (1x) and PBST (3 x, each with 15 mins rocking), then incubated with Goat Horse-radish peroxidase-conjugated anti mouse IgG antibodies (*Thermo Fisher*, 31430) diluted 1 in 25,000 in 5% v/v skimmed milk solution. The nitrocellulose membrane was subsequently washed with 30 mL of PBS (1x) and

PBST (3 x, each with 15 mins rocking), then antibody-bound bands visualised via treatment with ECL reagent (*Biorad*), as per manufacturer's instructions.

##### *2.1.11 Cellular NADPH Concentration Assays*

**Culturing of MIA PaCa-2 cells:** MIA PaCa-2 cells were maintained at 37°C and 5% $CO_2$. Cells were maintained in DMEM media (*Gibco*) supplemented with 10% *v/v* FBS (*Thermo-Fischer)* and 1% *v/v* Pen-Strep (*Thermo-Fischer*). Cells were passaged once per week, and were divided approximately 1 in 20 from a near confluent culture. Cells were passaged at least 3 times following thawing prior to use in experiments.

**Probe Treatment and NADPH-Concentration Determination:** MIA PaCa-2 cells were plated in 96 well plates (*Corning)* to a density of 400,000 cells per well in either regular or low glucose DMEM (supplemented in each case with 10% *v/v* FBS and 1% *v/v* Pen-Strep), and incubated at 37°C for 20 hours. Media was removed from the cells in each condition and replaced with the equivalent media DMEM supplemented with either **17** (at concentrations of 10 or 20 μM), ivosidenib (10 μM) or DMSO (1% *v/v*). In each compound-treated condition, DMSO was kept at 1% (*v/v*). Cells were incubated with compounds at 37°C for 24 hours, then NADPH concentrations determined using the *Total NADP and NADPH Assay Kit* (*Abcam,* ab186033), following protocols provided by the manufacturer.

#### 2.2 X-Ray Crystallography Methods and Statistics

##### 2.2.1 X-Ray Crystallography Methods

***Crystallisation screen plate setup and crystal harvesting:*** For the fragment-bound IDH1 crystals, IDH1-fragment complexes were produced via the *Fragment Labelled IDH1 Preparation Procedure*, whilst for the apo-IDH1 structure, the fragment preincubation step was skipped. IDH1 was concentrated to approximately 110 - 150 μM, then was treated with $NADP^+$(1 mM) and EDTA (1 mM), before being used to construct crystallisation screens. Both sitting and hanging drop screens were used. For sitting drop screens, mother liquor well conditions were constructed in MRC two-well crystallisation plates (*SwissCl*) using a Dragonfly crystal robot (*sptlabtech*). IDH1 and mother liquor solutions were combined in 200 nL:100 nL and 100 nL:100 nL (IDH1:mother liquor) drops *using a* Mosquito LV crystal robot (*sptlabtech*), then sealed, incubated at 16°C, and inspected regularly for signs of crystallisation. For hanging drop screens, 400 μL of mother liquor solution (2M $(NH_4)_2SO_4$, 4.27% PEG 400, 0.1 M Hepes 7.4) was aliquoted into EasyXtal hanging drop plates (*Qiagen*), then 1 μL of mother liquor was combined with 1 μL of protein solution on the glass slide, sealed, incubated at 16°C, and inspected regularly for signs of crystallisation. Successful crystals were harvested using nylon loops (*Molecular Dimensions*), briefly swirled in cryoprotectant solution (80:20, mother liquor:ethylene glycol) then flash frozen in liquid nitrogen and shipped to the Diamond Light Source for data collection.

***Data collection and model solving***: Diffraction data were either collected at the Diamond Light Source I24 beamline at a temperature of 100 K using an Eiger CdTe 9M detector, or for anomalous diffraction experiments on the I23 beamline under a vacuum using a Pilatus 12M detector (*Dectris*). Data reduction was performed using Xia2[8]. For the apoIDH1 structure, molecular replacement was performed using MOLREP[9] with a previously identified IDH1 structure (PDB-ID = 1T09)[10], whilst for further structures apo-IDH1 was used for subsequent molecular replacement.[9] All model fitting and refinement was performed using CCP4I2[11] via iterative cycles of optimisation using the Coot[12] and REFMAC[13] packages. Ligands were built using AceDRG[14]. Final model and refinement statistics were obtained using Molprobility[15] and figures created using the PYMOL molecular graphics system, version 3.1.3.1 (*Schrodinger*).

#### 2.2.2 X-Ray Crystallography Statistics

#### Apo-IDH1 Crystal Structure

| Data Collection | | Data Refinement | |
|---|---|---|---|
| **Wavelength (Å)** | 0.6199 | **R-work** | 0.204 |
| **Resolution range (Å)** | 41.35 - 2.52 (2.62–2.52) | **R-free** | 0.274 |
| **Space group** | $P\ 4_3\ 2_1\ 2$ | **Number of non-hydrogen atoms** | 6229 |
| **Unit cell dimensions (a,b,c) (Å)** | 83.700, 82.700, 301.360 | • **Macromolecules** | 6026 |
| **Total reflections** | 484412 (54980) | **Protein residues** | 789 |
| **Unique reflections** | 36596 (4029) | **RMS (bonds)** | 0.0067 |
| **Multiplicity** | 13.2 (13.6) | **RMS (angles)** | 1.655 |
| **Completeness** | 100.0 (100.0) | **Ramachandran favored/outliers (%)** | 92.66 / 0.13 |
| **Mean I/σ (I)** | 11.7 (0.9) | **Rotamer outliers (%)** | 2.13 |
| **Wilson B-factor (Å²)** | 61.27 | **Clash score** | 4.99 |
| **R-merge** | 0.159 (3.068) | **Average B-factor** | 74.88 |
| **R-meas** | 0.171 (3.322) | • **Macromolecules** | 74.59 |
| **R-pim** | 0.047 (0.893) | • **Ligands** | 91.17 |
| **CC1/2** | 0.999 (0.359) | • **Solvent** | 58.12 |

#### IDH1-1 Crystal Structure

| Data Collection | | Data Refinement | |
|---|---|---|---|
| **Wavelength (Å)** | 0.6199 | **R-work** | 0.220 |
| **Resolution range (Å)** | 61.05 – 2.27 (2.34–2.27) | **R-free** | 0.264 |
| **Space group** | $P\ 4_3\ 2_1\ 2$ | **Number of non-hydrogen atoms** | 6269 |
| **Unit cell dimensions (a,b,c) (Å)** | 82.706, 82.706, 305.255 | • **Macromolecules** | 6040 |
| **Total reflections** | 1342226 (124530) | **Protein residues** | 791 |
| **Unique reflections** | 50358 (4564) | **RMS (bonds)** | 0.0062 |
| **Multiplicity** | 26.7 (27.3) | **RMS (angles)** | 1.485 |
| **Completeness** | 100.0 (100.0) | **Ramachandran favored/outliers (%)** | 94.61 / 0.26 |
| **Mean I/σ (I)** | 26.7 (27.3) | **Rotamer outliers (%)** | 1.96 |
| **Wilson B-factor (Å²)** | 46.14 | **Clash score** | 3.7 |
| **R-merge** | 0.171 (3.578) | **Average B-factor** | 68.80 |
| **R-meas** | 0.174 (3.645) | • **Macromolecules** | 68.17 |
| **R-pim** | 0.034 (0.692) | • **Ligands** | 92.35 |
| **CC1/2** | 0.999 (0.506) | • **Solvent** | 55.73 |

#### IDH1-6 Crystal Structure

| Data Collection | | Data Refinement | |
|---|---|---|---|
| **Wavelength (Å)** | 0.6199 | **R-work** | 0.206 |
| **Resolution range (Å)** | 50.65 – 2.50 (2.60–2.50) | **R-free** | 0.267 |
| **Space group** | P $4_3$ $2_1$ 2 | **Number of non-hydrogen atoms** | 6267 |
| **Unit cell dimensions (a,b,c) (Å)** | 82.450, 82.450, 306.800 | • **Macromolecules** | 6032 |
| **Total reflections** | 457514 (46003) | **Protein residues** | 791 |
| **Unique reflections** | 37892 (4157) | **RMS (bonds)** | 0.0066 |
| **Multiplicity** | 12.1 (11.1) | **RMS (angles)** | 1.652 |
| **Completeness** | 100.0 (100.0) | **Ramachandran favored/outliers (%)** | 92.17 / 0.64 |
| **Mean I/σ (I)** | 13.1 (1.0) | **Rotamer outliers (%)** | 1.97 |
| **Wilson B-factor (Å$^2$)** | 61.58 | **Clash score** | 5.47 |
| **R-merge** | 0.132 (2.429) | **Average B-factor** | 71.72 |
| **R-meas** | 0.138 (2.548) | • **Macromolecules** | 7124 |
| **R-pim** | 0.039 (0.762) | • **Ligands** | 90.53 |
| **CC1/2** | 0.999 (0.339) | • **Solvent** | 60.18 |

#### IDH1-4 Crystal Structure

| Data Collection | | Data Refinement | |
|---|---|---|---|
| **Wavelength (Å)** | 0.6199 | **R-work** | 0.197 |
| **Resolution range (Å)** | 82.89 – 2.43 (2.52–2.43) | **R-free** | 0.225 |
| **Space group** | P $4_3$ $2_1$ 2 | **Number of non-hydrogen atoms** | 6250 |
| **Unit cell dimensions (a,b,c) (Å)** | 82.887, 82.887, 308.553 | • **Macromolecules** | 5976 |
| **Total reflections** | 749980 (66680) | **Protein residues** | 788 |
| **Unique reflections** | 41861 (4281) | **RMS (bonds)** | 0.0065 |
| **Multiplicity** | 17.9 (15.6) | **RMS (angles)** | 1.683 |
| **Completeness** | 100.0 (100.0) | **Ramachandran favored/outliers (%)** | 95.22 / 0.26 |
| **Mean I/σ (I)** | 16.5 (2.2) | **Rotamer outliers (%)** | 2.85 |
| **Wilson B-factor (Å$^2$)** | 56.63 | **Clash score** | 4.01 |
| **R-merge** | 0.100 (1.183) | **Average B-factor** | 64.58 |
| **R-meas** | 0.103 (1.222) | • **Macromolecules** | 63.87 |
| **R-pim** | 0.024 (0.307) | • **Ligands** | 90.25 |
| **CC1/2** | 0.997 (0.787) | • **Solvent** | 60.43 |

#### DH1-7 Crystal Structure

| Data Collection | | Data Refinement | |
|---|---|---|---|
| **Wavelength (Å)** | 0.6199 | **R-work** | 0.208 |
| **Resolution range (Å)** | 82.54 – 2.34 (2.42–2.34) | **R-free** | 0.265 |
| **Space group** | P $4_3$ $2_1$ 2 | **Number of non-hydrogen atoms** | 6411 |
| **Unit cell dimensions (a,b,c) (Å)** | 82.570, 82.570, 304.315 | • **Macromolecules** | 6146 |
| **Total reflections** | 1983837 (103060) | **Protein residues** | 789 |
| **Unique reflections** | 45762 (4393) | **RMS (bonds)** | 0.008 |
| **Multiplicity** | 43.4 (23.5) | **RMS (angles)** | 1.785 |
| **Completeness** | 100.0 (100.0) | **Ramachandran favored/outliers (%)** | 94.21 / 0.13 |
| **Mean I/σ (I)** | 17.5 (1.2) | **Rotamer outliers (%)** | 1.72 |
| **Wilson B-factor (Å²)** | 49.51 | **Clash score** | 2.7 |
| **R-merge** | 0.169 (3.307) | **Average B-factor** | 62.53 |
| **R-meas** | 0.171 (3.378) | • **Macromolecules** | 62.23 |
| **R-pim** | 0.025 (0.683) | • **Ligands** | 77.97 |
| **CC1/2** | 0.999 (0.535) | • **Solvent** | 55.29 |

#### IDH1-2 Anomalous Diffraction Crystal Structure

| Data Collection | | Data Refinement | |
|---|---|---|---|
| **Wavelength (Å)** | 2.7752 | **R-work** | 0.196 |
| **Resolution range (Å)** | 151.67 – 2.40 (2.49–2.40) | **R-free** | 0.254 |
| **Space group** | P $4_3$ $2_1$ 2 | **Number of non-hydrogen atoms** | 6319 |
| **Unit cell dimensions (a,b,c) (Å)** | 82.976, 82.976, 303.331 | • **Macromolecules** | 6076 |
| **Total reflections** | 4020889 (325728) | **Protein residues** | 801 |
| **Unique reflections** | 42776 (4378) | **RMS (bonds)** | 0.0074 |
| **Multiplicity** | 94.0 (74.4) | **RMS (angles)** | 1.767 |
| **Completeness** | 100.0 (100.0) | **Ramachandran favored/outliers (%)** | 92.21 / 0.00 |
| **Mean I/σ (I)** | 31.5 (1.7) | **Rotamer outliers (%)** | 1.48 |
| **Wilson B-factor (Å²)** | 74.48 | **Clash score** | 4.44 |
| **R-merge** | 0.124 (3.396) | **Average B-factor** | 79.91 |
| **R-meas** | 0.124 (3.419) | • **Macromolecules** | 79.60 |
| **R-pim** | 0.012 (0.388) | • **Ligands** | 95.39 |
| **CC1/2** | 1.000 (0.426) | • **Solvent** | 64.43 |

#### IDH1$^{C269W}$ Crystal Structure

| Data Collection | | Data Refinement | |
|---|---|---|---|
| **Wavelength (Å)** | 0.6199 | **R-work** | 0.203 |
| **Resolution range (Å)** | 50.43 – 2.30 (2.38–2.30) | **R-free** | 0.255 |
| **Space group** | P $4_3$ $2_1$ 2 | **Number of non-hydrogen atoms** | 6181 |
| **Unit cell dimensions (a,b,c) (Å)** | 82.420, 82.420, 302.610 | • **Macromolecules** | 5881 |
| **Total reflections** | 1272339 (126030) | **Protein residues** | 777 |
| **Unique reflections** | 47692 (4568) | **RMS (bonds)** | 0.0068 |
| **Multiplicity** | 26.7 (27.6) | **RMS (angles)** | 1.680 |
| **Completeness** | 100.0 (100.0) | **Ramachandran favored/outliers (%)** | 93.33 / 0.78 |
| **Mean I/σ (I)** | 17.8 (0.8) | **Rotamer outliers (%)** | 3.09 |
| **Wilson B-factor (Å$^2$)** | 59.39 | **Clash score** | 5.33 |
| **R-merge** | 0.141 (5.677) | **Average B-factor** | 69.64 |
| **R-meas** | 0.139 (5.544) | • **Macromolecules** | 69.45 |
| **R-pim** | 0.028 (1.094) | • **Ligands** | 81.23 |
| **CC1/2** | 0.999 (0.336) | • **Solvent** | 63.19 |

#### IDH1-15 Crystal Structure

| Data Collection | | Data Refinement | |
|---|---|---|---|
| **Wavelength (Å)** | 0.6199 | **R-work** | 0.207 |
| **Resolution range (Å)** | 51.30 – 2.40 (2.49–2.40) | **R-free** | 0.266 |
| **Space group** | P $4_3$ $2_1$ 2 | **Number of non-hydrogen atoms** | 6319 |
| **Unit cell dimensions (a,b,c) (Å)** | 82.922, 82.922, 307.811 | • **Macromolecules** | 6091 |
| **Total reflections** | 6198734 (360484) | **Protein residues** | 794 |
| **Unique reflections** | 50418 (4536) | **RMS (bonds)** | 0.0071 |
| **Multiplicity** | 122.9 (79.5) | **RMS (angles)** | 1.734 |
| **Completeness** | 100.0 (99.8) | **Ramachandran favored/outliers (%)** | 95.38 / 0.00 |
| **Mean I/σ (I)** | 24.5 (0.8) | **Rotamer outliers (%)** | 2.42 |
| **Wilson B-factor (Å$^2$)** | 59.16 | **Clash score** | 5.31 |
| **R-merge** | 0.188 (5.082) | **Average B-factor** | 70.10 |
| **R-meas** | 0.188 (5.112) | • **Macromolecules** | 69.13 |
| **R-pim** | 0.018 (0.548) | • **Ligands** | 101.84 |
| **CC1/2** | 1.000 (0.596) | • **Solvent** | 62.37 |

#### 2.3 Chemistry Materials and Methods

##### *2.3.1 General Chemical Considerations*

All reactions were performed in flame-dried glassware using anhydrous solvents and reagents from commercial sources. Flash column chromatography was either performed manually, using 0.2 – 0.3 mg average particle size silica (*Sigma Aldrich*) or using isolera columns (*Biotage*) with a Isolera One flash column chromatography system (*Biotage*). $R_f$ values were determined via thin-layer chromatography using aluminium-backed TLC plates (*Sigma Aldrich*) and visualised using UV absorbance and either potassium permanganate or ninhydrin staining.

All NMR experiments were performed using AV400 400 MHz spectrometers (*Bruker*), and chemical shifts reported in parts per million (ppm) relative to the peak of tetramethylsilane. Peak multiplicity is reported as follows: s = singlet, d = doublet, t = triplet, q = quartet, dd = doublet of doublets, dt = doublet of triplets, td = triplet of doublets, m = multiplet. In each case, the $CDCl_3$ peak ($\delta$ = 7.27 ppm) is used as the internal standard.

All high-resolution mass spectra (HRMS) were collected on the authors behalf by Malgorzata Puchnarewicz from the Imperial College London Department of Chemistry Mass Spectrometry service. Ionisation was performed by electrospray ionisation in positive ion mode using a LCT Premier ESI-ToF (*Walters*).

##### *2.3.2 Synthesis of Analogues of 2*

**Standard Procedure 1 – Synthesis of Sulfonamide**

Piperazine (6 equiv) was dissolved in dry DCM (0.25 M) and cooled in an ice-water bath. The corresponding sulfonyl chloride (1 equiv) and $NEt_3$ (1.5 equiv) were added dropwise and the reaction allowed to warm to room temperature, then left stirring until all sulfonyl chloride was consumed (as determined by TLC). The crude reaction mixture was washed with saturated aqueous 1M $NaHCO_3$ solution (50 mL), and brine (50 mL), then dried over $MgSO_4$, solvent removed *in vacuo*, and purified via column chromatography to yield the desired piperazine sulfonamide.

**Standard Procedure 2 – Synthesis of Chloroacetamides**

Using a procedure adapted from that reported by Allen et al.[16] Piperazine sulfonamide (1.0 equiv) was dissolved in dry DCM (0.3 M), then chroroacetyl chloride (1.1 equiv) added dropwise under stirring. The reaction mixture was stirred at RT for 10 mins, then amberlyst resin OH form (270 mg / 1 mmol of amine) added and stirred at RT for a further 10 mins. The resulting mixture was filtered through celite, solvent removed in vacuo, and the product purified via column chromatography.

**Synthesis of** 1-(methylsulfonyl)piperazine[17]

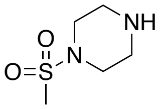


The desired product was synthesised from methylsulfonyl chloride (0.32 g, 2.8 mmol, 1 equiv), piperazine (1.45 g, 16.8 mmol, 6 equiv) and $NEt_3$ (585 μL, 4.2 mmol, 1.5 equiv) in DCM (11.2 mL, 0.25 M) according to standard procedure 1. The desired product was purified via flash column chromatography (gradient of 100% DCM to 20% MeOH in DCM) to yield 1-(methylsulfonyl) piperazine (270 mg, 1.65 mmol, 58%), $R_f$ = 0.2 (25% MeOH in DCM).
$^1$H NMR ($CDCl_3$, 400 MHz) δ 3.17 (4H, t, J = 4.5 Hz, SNC*H₂*) 2.94 (4H, t, J = 4.9 Hz, C$\underline{H_2}$NH), 2.75 (3H, s, C$\underline{H_3}$$SO_2$N), 1.65 (1H, s, N$\underline{H}$).
$^{13}$C NMR ($CDCl_3$, 101 MHz) δ 50.6, 46.7, 45.9, 45.6, 34.0. Analytical data were consistent with a previously reported synthesis.[17]

**Synthesis of** 2-chloro-1-(4-(methylsulfonyl)piperazin-1-yl)ethan-1-one (**18**)

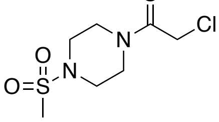


The desired product was synthesised from 1-(methyl sulfonyl) piperazine (73 mg, 0.442 mmol), chloroacetyl chloride (54.9 mg, 0.486 mmol) and amberlyst resin OH form (119 mg) in DCM (1.47 mL) according to standard procedure 2. The desired product was purified via flash column chromatography (50-100% EtOAc in hexane gradient), yielding 2-chloro-1-(4-(methylsulfonyl)piperazin-1-yl)ethan-1-one (**18**, 61.4 mg, 0.251 mmol, 57%) as a white solid. $R_f$ = 0.60 (100% EtOAc).

[1]H NMR ($CDCl_3$, 400 MHz) δ 4.08 (2H, s, $C\underline{H_2}Cl$), 3.75 (2H, t, J = 4.7 Hz, $C\underline{H_2}NCO$), 3.65 (2H, t, J = 4.9 Hz, $C\underline{H_2}NCO$), 3.31 (2H, t, J = 4.5 Hz, $SO_2NC\underline{H_2}$), 3.25 (2H, t, J = 5.1 Hz, $SO_2NC\underline{H_2}$), 2.81 (3H, s, $C\underline{H_3}SO_2N$).
[13]C NMR ($CDCl_3$, 101 MHz) δ 165.1, 45.8, 45.2, 41.5, 40.4, 34.8.
HRMS ($ESI^+$) $C_7H_{14}N_2O_3SCl^+$ $[M + H]^+$ calculated mass = 241.0406, found 241.0414 (Δ = 3.3 ppm).

**Synthesis of** 1-(phenylsulfonyl)piperazine

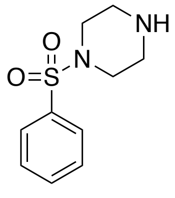

The desired product was synthesised from benzenesulfonyl chloride (0.5 g, 2.8 mmol, 1 eq), piperazine (1.45 g, 16.8 mmol, 6 eq) and $NEt_3$ (585 μL, 4.2 mmol) in DCM (11.2 mL) according to standard procedure 1. The desired product was purified via flash column chromatography (gradient of 100% DCM to 20% MeOH in DCM) to yield 1-(benzenesulfonyl) piperazine (493 mg, 2.156 mmol, 77%) as a white solid. $R_f$ = 0.65 (20% MeOH in DCM).
[1]H NMR ($CDCl_3$, 400 MHz) δ 7.75 (2H, d, J = 7.6 Hz, Ar-$\underline{H}$), 7.60 (1H, t, J = 7.0 Hz, Ar-$\underline{H}$), 7.53 (2H, t, J = 7.7 Hz, Ar-$\underline{H}$), 2.98 (4H, m, $SO_2NC\underline{H_2}$), 2.93 (4H, m, $C\underline{H_2}NH$), 1.48 (1H, s, $N\underline{H}$).
[13]C NMR ($CDCl_3$, 101 MHz) δ 135.4, 132.8, 129.0, 127.8, 46.9, 45.3.
Analytical data were consistent with a previously reported synthesis.[17]

**Synthesis of** 2-chloro-1-(4-(phenylsulfonyl)piperazin-1-yl)ethan-1-one (**10**)

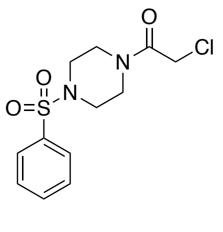

The desired product (**10**) was synthesised from 1-(benzenesulfonyl) piperazine (100 mg, 0.442 mmol) and chloroacetyl chloride (54.9 mg, 0.486 mmol, 1.1 equiv) in DCM (1.47 mL, 0.3 M) with amberlyst resin OH form (119 mg) according to standard procedure 2. The desired product was purified via flash column chromatography (gradient from 50% EtOAc in hexane to 100%), yielding 2-chloro-1-(4-(methylsulfonyl)piperazin-1-yl)ethan-1-one (**10**, 107.9 mg, 0.358 mmol, 81.1%) as a white solid. $R_f$ = 0.8 (100% EtOAc).

[1]H NMR ($CDCl_3$, 400 MHz) δ 7.70 (2H, d, J = 7.9 Hz, Ar-*H*), 7.60 (1H, t, J = 7.3 Hz, Ar-*H*), 7.52 (2H, t, J = 7.8 Hz, Ar-*H*), 3.98 (2H, s, C*H*2Cl), 3.66 (2H, t, J = 5.4 Hz, C*H*2NCO), 3.57 (2H, t, J = 6.6 Hz, C*H*2NCO), 3.04 (2H, t, J = 4.9 Hz, $SO_2NCH_2$), 2.99 (2H, t, J = 4.9 Hz, $SO_2NCH_2$).
[13]C NMR ($CDCl_3$, 101 MHz) δ 165.1, 135.2, 133.4, 129.4, 127.6, 46.0, 45.7, 41.4, 40.7.
HRMS ($ESI^+$) $C_{12}H_{15}ClN_2O_3S^+$ $[M + H]^+$ calculated mass = 303.0570, found 303.0572 (Δ = 0.7 ppm).

**Synthesis of** 1-(*o*-toluenesulfonyl)piperazine

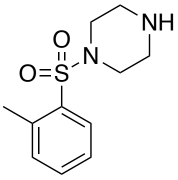

The desired product was synthesised from *o*-toluenesulfonyl chloride (0.53 g, 2.8 mmol, 1 eq), piperazine (1.45 g, 16.8 mmol, 6 eq) and $NEt_3$ (585 μL, 4.2 mmol) in DCM (11.2 mL) according to standard procedure 1. The desired product was purified via flash column chromatography (100% DCM to 10% MeOH in DCM) to yield 1-(*o*-toluenesulfonyl)piperazine (590 mg, 2.46 mmol, 88%). $R_f$ (100% EtOAc) = 0.4.
[1]H NMR ($CDCl_3$, 400 MHz) δ 7.84 (1H, m, J = 8.5 Hz, Ar-*H*), 7.42 (1H, t, J = 8.0 Hz, Ar-*H*), 7.29 – 7.28 (2H, m, Ar-*H*), 3.08 – 3.07 (4H, m, $SO_2NCH_2$), 2.86 – 2.85 (4H, m, $CH_2NH$), 2.60 (3H, s, Ar-$CH_3$), 1.58 (1H, s, N*H*).
[13]C NMR ($CDCl_3$, 101 MHz) δ 137.9, 135.2, 132.8, 130.2, 126.0, 46.1, 45.4, 20.7.
HRMS ($ESI^+$) $C_{11}H_{17}N_2O_2S$ $[M + H]^+$ calculated mass = 241.1011, found 241.1003 (Δ = 3.3 ppm).

**Synthesis of** 2-chloro-1-(4-(*o*-toluenesulfonyl)piperazin-1-yl)ethan-1-one (**11**)

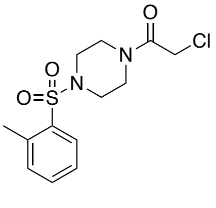

The desired product was synthesised from 1-(*o*-toluenesulfonyl)piperazine (106 mg, 0.442 mmol) and chloroacetyl chloride (54.9 mg, 0.486 mmol, 1.1 equiv) in DCM (1.47 mL, 0.3 M) with amberlyst resin OH form (119 mg) according to standard procedure 2. The desired product was purified via flash column chromatography (100% DCM), yielding 2-chloro-1-(4-

(*o*-toluenesulfonyl)piperazin-1-yl)ethan-1-one (**11**, 119.8 mg, 0.407 mmol, 92%) as a white solid. $R_f$ = 0.60 (25% EtOAc in DCM).

$^1$H NMR ($CDCl_3$, 400 MHz) δ 7.84 – 7.82 (1H, m, Ar-*H*), 7.45 (1H, t, J = 7.7 Hz, Ar-*H*), 7.30 (2H, m, Ar-*H*), 4.01 (2H, s, C*H*$_2$Cl), 3.64 (2H, t, J = 5.4 Hz, C*H*$_2$NCO), 3.55 (2H, t, J = 4.9 Hz, C*H*$_2$NCO), 3.20 (2H, t, J = 4.9 Hz, C*H*$_2$NCO), 3.14 (2H, t, J = 5.4 Hz, $SO_2$NC*H*$_2$), 2.58 (3H, s, Ar-C*H*$_3$).

$^{13}$C NMR ($CDCl_3$, 101 MHz) δ 165.2, 138.0, 135.0, 133.4, 133.1, 130.3, 126.4, 45.9, 45.2, 44.9, 41.7, 40.7, 20.8.

HRMS (ESI$^+$) $C_{13}H_{18}N_2O_2SCl$ [M + H]$^+$ calculated mass = 317.0727, found 317.0730 (Δ = 0.9 ppm).

**Synthesis of** 1-(4-((2-(*tert*-butyl)phenyl)sulfonyl)piperazin-1-yl)-2-chloroethan-1-one (**12**)

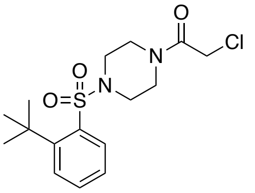


Piperazine (557 mg, 6.47 mmol, 6 equiv) and $NEt_3$ (225 μL, 163 mg, 9.71 mmol, 1.5 equiv) were dissolved in DCM (4.32 mL), cooled on ice, then 2-*ter-t*butyl benzene sulfonyl chloride (250 mg, 1.08 mmol, 1 equiv) added dropwise. The reaction mixture was stirred for 1 h, then washed with water (3 x 10 mL) and brine (10 mL), and dried over $MgSO_4$. The desired product was purified as a white solid (262 mg, 0.926 mmol, 86%), which was used in the next step without further characterisation.

1-((2-(*tert*-butyl)phenyl)sulfonyl)piperazine (125 mg, 0.442 mmol, 1 eq) was dissolved in DCM (1.25 mL, 0.3 M), then chloroacetyl chloride (54.9 mg, 0.489 mmol, 1.1 eq) added dropwise and stirred for 10 mins. Amberlyst resin OH form (299.2 mg) was added and the mixture stirred for a further 10 mins. The reaction mixture was filtered through celite and purified via flash column chromatography (40% EtOAc in hexane), yielding the desired product as a colourless oil (**12**, 137 mg, 87%). $R_f$ (DCM) = 0.5.

$^1$H NMR ($CDCl_3$, 400 MHz) δ 7.76 - 7.68 (2H, m, Ar-*H*), 7.49 (1H, m, Ar-*H*), 7.31 (1H, m, Ar-*H*), 4.07 (2H, s, C*H*$_2$Cl), 3.67 (2H, t, J = 5.2 Hz, NC*H*$_2$), 3.60 (2H, t, J = 4.6 Hz, C*H*$_2$NCO), 3.39 (2H, t, J = 5.4 Hz, $SO_2$NC*H*$_2$), 3.33 (2H, t, J = 5.4 Hz, $SO_2$NC*H*$_2$), 1.57 (9H, s, Ar-C(C*H*$_3$)$_3$).

$^{13}$C NMR ($CDCl_3$, 101 MHz) δ 165.1, 150.6, 138.2, 132.6, 130.2, 129.6, 126.0, 46.2, 45.4, 45.2, 41.8, 40.5, 37.1, 31.9.

HRMS (ESI$^+$) $C_{16}H_{23}ClN_2O_3S^+$ [M + H]$^+$ calculated mass = 359.1192, found 359.1196 (Δ = 1.1 ppm).

**Synthesis of** *tert*-butyl 4-((2-nitrophenyl)sulfonyl)piperazine-1-carboxylate

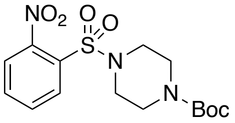

1-Boc piperazine (931 mg, 5.01 mmol, 1.1 equiv), and triethylamine (940 μL, 682.4 mg, 6.76 mmol, 1.5 equiv) were dissolved in DCM (45 mL, 0.1 M), cooled on ice, then 2-nitrobenzene sulfonyl chloride (1 g, 4.55 mmol, 1 equiv) added dropwise. The reaction was warmed to room temperature, then stirred for 30 mins. The reaction mixture was washed with water (20 mL) and brine (20 mL), then dried over $MgSO_4$. The solvent was removed *in vacuo,* yielding *tert*-butyl 4-((2-nitrophenyl)sulfonyl)piperazine-1-carboxylate (1.581 g, 4.26 mmol, 94%) as a colourless oil. $R_f$ = 0.5 (50% EtOAc in hexane).

$^1$H NMR (400 MHz, CDCl3) δ 7.97 (1H, dd, J = 7.3, 1.9 Hz, Ar-*H*), 7.75 – 7.67 (2H, m, Ar-*H*), 7.63 – 7.61 (1H, m, Ar-*H*), 3.52 – 3.50 (4H, m, C*H₂*NBoc), 3.28 – 3.25 (4H, m, $SO_2NCH_2$), 1.43 (9H, s, $C(CH_3)_3$)

$^{13}$C NMR (101 MHz, CDCl3) δ 154.3, 148.5, 134.1, 131.7,131.1, 131.1, 124.3, 80.6, 45.9, 28.4.

HRMS (ESI$^+$) $C_{15}H_{21}N_3O_6S^+$ $[M + H]^+$ calculated mass = 371.1157, found 371.1154 (Δ = 0.8 ppm).

**Synthesis of** *tert*-butyl 4-((2-aminophenyl)sulfonyl)piperazine-1-carboxylate

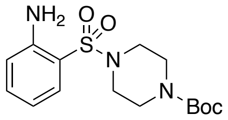

*Tert*-butyl 4-((2-nitrophenyl)sulfonyl)piperazine-1-carboxylate (1.581 g, 4.26 mmol, 1 equiv), Fe powder (1.19 g, 21.3 mmol, 5 equiv) and $NH_4Cl$ (1.82 g, 34.08 mmol, 8 equiv) were combined in EtOH (17.18 mL) / $H_2O$ (4.29 mL) then stirred at 80°C for 4 h. The reaction mixture was filtered, then extracted with DCM (2 x 50 mL). The combined organic extracts were washed with water (50 mL) and brine (50 mL), then dried over $MgSO_4$, yielding *tert*-butyl 4-((2-aminophenyl)sulfonyl)piperazine-1-carboxylate (1.3 g, 3.81 mmol, 89%) as an off white solid. $R_f$ = 0.6 (50% EtOAc in hexane).

$^1$H NMR (400 MHz, $CDCl_3$) δ 7.52 (1H, d, J = 7.2, Ar-*H*), 7.32 – 7.25 (1H, m, Ar-*H*), 6.78 – 6.70 (2H, m, Ar-*H*), 5.05 (2H, s, $Ar-NH_2$), 3.47 – 3.46 (4H, m, $CH_2$NBoc), 3.07 – 3.06 (4H, m, $SO_2NCH_2$), 1.41 (9H, s, $C(CH_3)_3$).

$^{13}$C NMR (101 MHz, $CDCl_3$) δ 154.3,146.5, 134.6, 130.5, 117.9, 117.5, 117.3, 80.5, 45.9, 28.5.

HRMS (ESI$^+$) $C_{15}H_{24}N_3O_4S^+$ $[M + H]^+$ calculated mass = 342.1488, found 342.1476 ($\Delta$ = 3.5 ppm).

**Synthesis of** 1-benzyl-3-(2-((4-(2-chloroacetyl)piperazin-1-yl)sulfonyl)phenyl)urea (**13**)

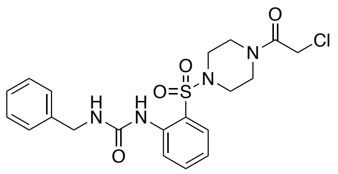

Triphosgene (5.8mg, 0.0217 mmol, 0.37 equiv) was dissolved in DCM (50 μL), cooled on ice, then *tert*-butyl 4-((2-aminophenyl)sulfonyl)piperazine-1-carboxylate (20 mg, 0.0587 mmol, 1 equiv) and pyridine (2.36 μL, 2.32 mg, 0.0587 mmol, 1 equiv) in DCM (150 μL) were added dropwise. The reaction was allowed to warm to room temperature and stirred for 2 h. The reaction was then cooled on ice, and benzylamine (6.29 mg, 0.0587 mmol, 1 equiv) and pyridine (2.36 μL, 2.32 mg, 0.0587 mmol, 1 equiv) in DCM (85 μL) was added dropwise. The reaction was allowed to warm to room temperature, then stirred overnight. The reaction was quenched via the addition of 1M $NaHCO_3$ solution (10 mL) which was subsequently stirred for 30 mins. The reaction was next diluted with DCM (5 mL), washed with $H_2O$ (10 mL), HCl (10 mL) and brine (20 mL), then dried over $MgSO_4$ and purified via flash column chromatography (40% EtoAc in hexane) to yield *tert*-butyl 4-((2-(3-benzylureido)phenyl)sulfonyl)piperazine-1-carboxylate (23 mg, 0.0485 mmol, 83%) as a white solid, which was proceeded to the next step without further purification.

*Tert*-butyl 4-((2-(3-benzylureido)phenyl)sulfonyl)piperazine-1-carboxylate (23 mg, 0.0485 mmol, 1 equiv) was dissolved in THF (230 mL), cooled on ice, then 5-6 M HCl in propanol (230 μL) added dropwise. The reaction was allowed to warm to room temperature and stirred for 3 h. $H_2O$ (5 mL) was next added, then basified via the dropwise addition of 3M NaOH solution until the pH reached 14. The aqueous mixture was then extracted with DCM (2 x 5 mL), and the combined organic extracts washed with brine (10 mL) and dried over $MgSO_4$. The resulting free amine was subsequently dissolved in DCM (162 μL, 0.3 M), then chloroacetyl chloride (6.5 mg, 0.0582 mmol, 1.2 equiv) added dropwise and the reaction stirred at room temperature for 10 mins. Amberlyst resin OH form was then added, and the reaction stirred for a further 10 mins. The reaction mixture was next filtered through celite, and purified via flash column chromatography (0-100% DCM in hexane gradient) to yield 1-benzyl-3-(2-((4-(2-chloroacetyl)piperazin-1-yl)sulfonyl)phenyl)urea (**13**, 8.5 mg, 0.0188 mmol, 39%) as a white solid. $R_f$ = 0.5 (50:50 EtOAc in DCM).

$^1$H NMR ($CDCl_3$, 400 MHz) δ 8.52 (1H, s (br), Ar-N*H*), 8.25 (1H, d, J = 8.4 Hz, Ar-*H*), 7.66 (1H, dd, J = 8.1,1.6 Hz, Ar-*H*), 7.55 – 7.51 (1H, m, Ar-*H*), 7.37 – 7.29 (5H, m, Ar-*H*), 7.14 – 7.09 (1H, m, Ar-*H*), 5.50 (1H, t, J = 5.8 Hz, $ArCH_2NH$), 4.44 (2H, d, J = 6.0 Hz, Ar-C*H2*), 3.96 (2H, s, C*H2*Cl), 3.57 (2H, t, J = 4.9 Hz, C*H2*NCO), 3.43 (2H, J = 4.6 Hz, C*H2*NCO), 3.00 - 2.99 (4H, m, $SO_2NCH_2$).

$^{13}$C NMR ($CDCl_3$, 101 MHz) δ 165.1, 154.3, 138.1, 134.7, 129.6, 128.8, 127.1, 123.2, 122.6, 122.2, 45.5, 44.5, 41.3, 40.6.

HRMS (ESI$^+$) $C_{20}H_{23}N_4O_4SCl$ [M + H]$^+$ calculated mass = 451.1207, found 451.1198 (Δ = 2.0 ppm).

**Synthesis of** *tert*-butyl 4-((2-(3-(quinolin-4-ylmethyl)ureido)phenyl)sulfonyl)piperazine-1-carboxylate

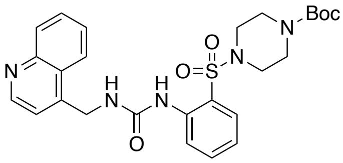

Triphosgene (60 mg, 0.202 mmol, 0.37 equiv) was dissolved in DCM (100 μL), cooled on ice, then *tert*-butyl 4-((2-aminophenyl)sulfonyl)piperazine-1-carboxylate (207 mg, 0.613 mmol, 1 equiv) and pyridine (24.5 μL, 23.5 mg, 0.613 mmol, 1 equiv) in DCM (3.05 mL) were added dropwise. The reaction was allowed to warm to room temperature and stirred for 2 h. The reaction was then cooled on ice, and 4-quinolinylmethyl)amine (154 mg, 0.975 mmol, 1.6 equiv) and pyridine (24.5 μL, 23.5 mg, 0.613 mmol, 1 equiv) in DCM (3.05 mL) was added dropwise. The reaction was allowed to warm to room temperature, then stirred overnight, before being quenched via the addition of 1M $NaHCO_3$ solution (10 mL) which was subsequently stirred for 30 mins. The reaction was next diluted with DCM (5 mL), washed with $H_2O$ (10 mL), HCl (10 mL) and brine (20 mL), then dried over $MgSO_4$ and purified via flash column chromatography (40% EtoAc in hexane) to yield *tert*-butyl 4-((2-(3-(quinolin-4-ylmethyl)ureido)phenyl)sulfonyl)piperazine-1-carboxylate (150 mg, 0.286 mmol, 47%) as a colourless oil. $R_f$ = 0.2 (50:50 EtOAc/hexane).

$^1$H NMR ($CDCl_3$, 400 MHz) δ 8.71 (1H, s, Ar-*H*), 8.70 (1H, s, Ar-N*H*), 8.25 (1H, d, J = 8.6 Hz, Ar-*H*), 8.03 (1H, d, J = 7.1, Ar-*H*), 7.98 (1H, d, J = 7.0 Hz, Ar-*H*), 7.65 (1H, t, J = 8.3 Hz, Ar-*H*), 7.58 (1H, dd, J = 8.1, 1.6 Hz, Ar-*H*), 7.50 (1H, t, J = 8.3, Ar-*H*), 7.45 (1H, t, J = 8.8 Hz, Ar-*H*), 6.57 (1H, J = 5.6 Hz, Ar-$CH_2NH$), 4.86 (2H, d, J = 5.7 Hz, Ar-C*H2*), 3.31 (4H, t, J = 4.6 Hz, C*H2*NCO), 2.89 (4H, s (br), $SO_2NCH_2$), 1.38 (9H, s, $C(CH_3)_3$).

$^{13}$C NMR ($CDCl_3$, 101 MHz) δ 154.2, 154.0, 150.9, 146.6, 137.4, 134.6, 134.5, 130.3, 129.9, 123.1, 122.7, 122.2, 117.8, 117.2, 116.9, 80.6, 80.3, 45.8, 28.3.

HRMS (ESI$^+$) $C_{26}H_{31}N_5O_5S$ [M + H]$^+$ calculated mass = 526.2124, found 526.2115 (Δ = 1.7 ppm).

**Synthesis of** 1-(2-((4-(2-chloroacetyl)piperazin-1-yl)sulfonyl)phenyl)-3-(quinolin-4-ylmethyl)urea (**14**)

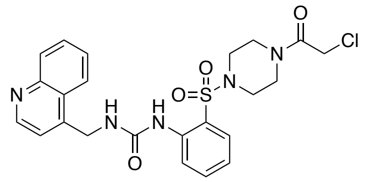

4-((2-(3-(Quinolin-4-ylmethyl)ureido)phenyl)sulfonyl)piperazine-1-carboxylate (85 mg, 0.184 mmol, 1 equiv) was add to 5% TFA (*v/v*) in DCM (1.84 mL, 0.1 M), then stirred at room temperature for 4 h. Water (10 mL) was added, then the solution basified via the dropwise addition of 3 M NaOH until it reached a pH 12. The free amine was then extracted with DCM (2 x 10 mL), washed with brine (10 mL), dried over $MgSO_4$ and used in the next step without characterisation.

1-(2-(piperazin-1-ylsulfonyl)phenyl)-3-(quinolin-3-ylmethyl)urea (46.9 mg, 0.170 mmol, 1 equiv) was dissolved in DCM (0.875 ml, 0.2 M) then chloroacetyl chloride added (14.9 μL, 21.1 mg, 186 mmol, 1.1 equiv) and the reaction stirred for 10 mins at room temperature. Amberlyst resin (50.5 mg) was subsequently added and the reaction stirred for a further 10 mins. The reaction mixture was filtered through celite, then purified via flash column chromatography (gradient of 80% -100% EtOAc in hexane) to yield 1-(2-((4-(2-chloroacetyl)piperazin-1-yl)sulfonyl)phenyl)-3-(quinolin-4-ylmethyl)urea (**14**, 27.6 mg, 0.0550 mmol, 32%) as a colourless oil. $R_f$ = 0.35 (100% EtOAc).

$^1$H NMR ($CDCl_3$, 400 MHz) δ 8.73 (1H, d, J = 4.4 Hz, Ar-*H*), 8.54 (1H, s (br), Ar-N*H*), 8.16 (1H, d, J = 8.3 Hz, Ar-*H*), 8.06 (1H, d, J = 8.2 Hz, Ar-*H*), 7.97 (1H, d, J = 8.1 Hz, Ar-*H*), 7.67 (1H, t, J = 7.0 Hz, Ar-*H*), 7.59 (1H, d, J = 6.4 Hz, Ar-*H*), 7.51 (1H, t, J = 7.1 Hz, Ar-*H*), 7.44 (1H, t, J = 7.1 Hz, Ar-*H*), 7.31 (1H, d, J = 4.5 Hz, Ar-*H*), 7.06 (1H, t, J = 7.5 Hz, Ar-*H*), 6.78 (1H, s (br), $CH_2$N*H*), 4.84 (2H, d, J = 5.9 Hz, ArC$H_2$NH), 3.93 (2H, s, C$H_2$Cl), 3.45 (2H, s (br), C$H_2$NCO), 3.30 (2H, s (br), C$H_2$NCO), 2.93 (4H, s (br), $SO_2$NC$H_2$).

$^{13}$C NMR ($CDCl_3$, 101 MHz) δ 157.0, 154.2, 149.8, 147.6, 144.2, 137.7, 129.6, 129.5, 126.9, 126.1, 123.1, 122.8, 122.4, 122.2, 119.5, 45.1, 41.0, 40.8, 40.5.

HRMS (ESI$^+$) $C_{23}H_{25}N_5O_5SCl$ [M + H]$^+$ calculated mass = 502.1316, found 502.1321 (Δ = 1.0 ppm).

**Synthesis of** ***tert*****-butyl 4-((2-methyl-6-nitrophenyl)sulfonyl)piperazine-1-carboxylate**

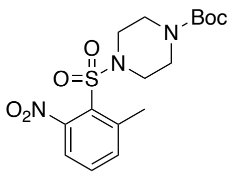


2-Methyl 6-nitro benzenesulfonyl chloride (500 mg, 2.13 mmol, 1 equiv) and DIPEA (543 μL, 425 mg, 3.3 mmol, 1.5 equiv) were dissolved in DCM (7 mL, 0.3 M) and cooled on ice. 1-boc piperazine (396.2 mg, 2.13 mmol, 1 equiv) was then added, and the reaction allowed to warm to room temperature and stirred for 30 mins. The reaction mixture was subsequently diluted with DCM, washed with 1 M HCl (2 x 20 mL), water (20 mL) and brine (20 mL), then dried over $MgSO_4$ and solvent removed *in vacuo.* Purification via flash column chromatography (gradient of 0 – 40% DCM in hexane) yielded *tert*-butyl 4-((2-methyl-6-nitrophenyl)sulfonyl)piperazine-1-carboxylate (350 mg, 0.9 mmol, 43%) as a pale yellow oil. $R_f$ = 0.3 (100% DCM).

$^1$H NMR ($CDCl_3$, 400 MHz) δ 7.72 (1H, t, J = 7.7 Hz, Ar-*H*), 7.47 (1H, d, J = 7.9 Hz, Ar-*H*), 7.30 (1H, d, J = 7.8 Hz, Ar-*H*), 3.49 (4H, t, J = 5.0 Hz, C*H₂*NCO), 3.31 (4H, t, J = 5.4 Hz, $SO_2NCH_2$), 2.71 (3H, s, Ar-$CH_3$), 1.45 (9H, s, $C(CH_3)_3$).

$^{13}$C NMR ($CDCl_3$, 101 MHz) δ 154.4, 141.6, 135.2, 133.3, 121.8, 80.7, 45.5, 28.5, 21.9.

HRMS (ESI$^+$) $C_{16}H_{23}N_3O_6S$ $[M + H]^+$ calculated mass = 403.1651, found 403.1660 (Δ = 2.2 ppm).

**Synthesis of** *tert*-butyl 4-((2-amino-6-methylphenyl)sulfonyl)piperazine-1-carboxylate

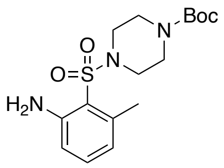


*Tert*-butyl 4-((2-methyl-6-nitrophenyl)sulfonyl)piperazine-1-carboxylate (350 mg, 0.9 mmol, 1 equiv), Fe powder (250 mg, 4.5 mmol, 5 equiv) and $NH_4Cl$ (389 mg, 7.2 mmol, 8 equiv) were combined in EtOH (3.44 mL) / $H_2O$ (0.86 mL) then stirred at 80°C for 4 h. The reaction mixture was filtered, then extracted with DCM (2 x 10 mL). The combined organic extracts were washed with water (2 x 20 mL) and brine (20 mL), then dried over $MgSO_4$, yielding *tert*-butyl 4-((2-amino-6-methylphenyl)sulfonyl)piperazine-1-carboxylate (297 mg, 0.834 mmol, 93%) as a white solid. $R_f$ = 0.35 (100% DCM).

$^1H$ NMR ($CDCl_3$, 400 MHz) δ 7.12 (1H, t, J = 7.7 Hz, Ar-*H*), 6.56 – 6.53 (2H, m, Ar-*H*), 3.46 (4H, t, J = 5.0 Hz, C*H₂*NCO), 3.16 (4H, t, J = 5.2 Hz, $SO_2NCH_2$), 2.54 (3H, s, Ar-$CH_3$), 1.44 (9H, s, $C(CH_3)_3$).
$^{13}C$ NMR ($CDCl_3$, 101 MHz) δ 154.8, 149.3, 140.7, 133.9, 121.9, 116.8, 80.8, 45.1, 28.8, 23.1.
HRMS (ESI$^+$) $C_{16}H_{26}N_3O_4S$ $[M + H]^+$ calculated mass = 356.1644, found 356.1646 (Δ = 0.6 ppm).

**Synthesis of** (*tert*-butyl 4-((2-methyl-6-(3-(quinolin-4-ylmethyl)ureido)phenyl)sulfonyl)piperazine-1-carboxylate)

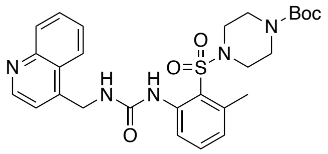


Triphosgene (64.4mg, 0.216 mmol, 0.37 equiv) was dissolved in DCM (100 μL), cooled on ice, then *tert*-butyl 4-((2-amino-6-methylphenyl)sulfonyl)piperazine-1-carboxylate (208 mg, 0.584 mmol, 1 equiv) and pyridine (47.3 μL, 46.53 mg, 0.584 mmol, 1 equiv) in DCM (2.94 mL) were added dropwise. The reaction was allowed to warm to room temperature and stirred for 2 h. The reaction was then cooled on ice, and 4-quinolinylmethyl)amine (149 mg, 0.943 mmol, 1.6 equiv) and pyridine (47.3 μL, 46.53 mg, 0.584 mmol, 1 equiv) in DCM (2.94 mL) was added dropwise. The reaction was allowed to warm to room temperature, then stirred overnight. The reaction was quenched via the addition of 1M $NaHCO_3$ solution (10 mL) which was subsequently stirred for 30 mins. The reaction was next diluted with DCM (5 mL), washed with $H_2O$ (10 mL), HCl (10 mL) and brine (20 mL), then dried over $MgSO_4$ and purified via flash column chromatography (70% EtoAc in pentane) to yield *tert*-butyl 4-((2-methyl-6-(3-(quinolin-4-ylmethyl)ureido)phenyl)sulfonyl)piperazine-1-carboxylate (304 mg, 0.563 mmol, 96%) as a colourless oil. $R_f$ = 0.6 (EtOAc).
$^1H$ NMR ($CDCl_3$, 400 MHz) δ 9.42 (1H, s, Ar-N*H*), 8.86 (1H, d, J = 4.4 Hz, Ar-*H*), 8.15 (2H, q, J = 9.0, 7.1 Hz, Ar-*H*), 8.05 (1H, d, J = 8.3 Hz, Ar-*H*), 7.73 (1H, t, J = 8.3 Hz, Ar-*H*), 7.59 (1H, t, J = 8.2 Hz, Ar-*H*), 7.41 – 7.36 (2H, m, Ar-*H*), 6.94 (1H, d, J = 7.6 Hz, Ar-*H*), 5.52 (1H, t, J = 5.9 Hz, $ArCH_2NH$), 4.95 (2H, d, J = 6.0 Hz, $ArCH_2$), 3.38 (4H, t, J = 4.8 Hz, $CH_2NCO$), 3.05 (4H, t, J = 4.5 Hz, $SO_2NCH_2$), 2.58 (3H, s, Ar-$CH_3$), 1.44 (9H, s, $C(CH_3)_3$).
$^{13}C$ NMR ($CDCl_3$, 101 MHz) δ 154.6, 154.3, 150.4, 148.3, 144.0, 140.4, 140.1, 133.6, 130.3, 129.6, 127.2, 127.1, 126.5, 123.1, 121.6, 121.1, 119.7, 80.7, 44.6, 41.2, 28.4, 23.1.
HRMS (ESI$^+$) $C_{27}H_{34}N_5O_5S$ $[M + H]^+$ calculated mass = 540.2281, found 540.2258 (Δ = 4.3 ppm).

**Synthesis of** 1-(2-((4-(2-chloroacetyl)piperazin-1-yl)sulfonyl)-3-methylphenyl)-3-(quinolin-4-ylmethyl)urea (**15**)

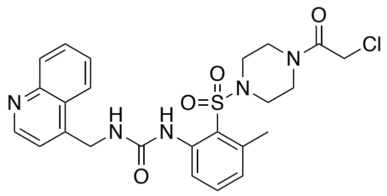

*Tert*-butyl 4-((2-methyl-6-(3-(quinolin-4-ylmethyl)ureido)phenyl)sulfonyl)piperazine-1-carboxylate (300 mg, 0.557 mmol, 1 equiv) was add to 5% TFA (*v/v*) in DCM (5.57 mL, 0.1 M), then stirred at room temperature for 4 h. Water (20 mL) was added, then the solution basified via the dropwise addition of 3 M NaOH until it reached pH 12. The free amine was then extracted with DCM (2 x 20 mL), washed with brine (50 mL), dried over $MgSO_4$ and used in the next step without characterisation.

1-(3-methyl-2-(piperazin-1-ylsulfonyl)phenyl)-3-(quinolin-4-ylmethyl)urea (110 mg, 0.234 mmol, 1 equiv) was dissolved in DCM (1.2 mL, 0.2 M) then chloroacetyl chloride added (18.6 μL, 26.3 mg, 0.192 mmol, 1 equiv) and the reaction stirred for 10 mins at room temperature. Amberlyst resin OH form (63.4 mg) was subsequently added and the reaction stirred for a further 10 mins. The reaction mixture was filtered through celite, then purified via flash column chromatography (gradient of 80 - 100% EtOAc in hexane) to yield 1-(2-((4-(2-chloroacetyl)piperazin-1-yl)sulfonyl)-3-methylphenyl)-3-(quinolin-4-ylmethyl)urea (**15**, 89.1 mg, 0.172 mmol, 74%) as a colourless oil. $R_f$ = 0.25 (100% EtOAc).

$^1$H NMR ($CDCl_3$, 400 MHz) δ 9.08 (1H, s, ArN*H*), 8.76 (1H, d, J = 4.4 Hz, Ar-*H*), 8.05 (1H, d, J = 8.5 Hz, Ar-*H*), 8.01-7.98 (2H, m, Ar-*H*), 7.69 – 7.64 (1H, m, Ar-*H*), 7.50 (1H, t, J = 8.3 Hz, Ar-*H*), 7.34 (1H, d, J = 4.5 Hz, Ar-*H*), 7.29 (1H, t, J = 7.7 Hz, Ar-*H*), 6.89 (1H, d, J = 7.2 Hz, Ar-*H*), 6.65 (1H, s, $ArCH_2N$*H*), 4.84 (2H, d, J = 5.9 Hz, ArC$H_2$NH), 3.94 (2H, s, C$H_2$Cl), 3.39 (2H, t, J = 4.6 Hz, C$H_2$NCO), 3.26 (2H, t, J = 4.8 Hz, C$H_2$NCO), 3.00 (2H, t, J = 5.6 Hz, C$H_2$$NSO_2$), 2.96 (2H, t, J = 5.2 Hz, C$H_2$$NSO_2$)2.51 (3H, s, Ar-C$H_3$).

$^{13}$C NMR ($CDCl_3$, 101 MHz) δ 165.4, 154.9, 150.3, 148.1, 144.4, 140.2, 140.0, 133.6, 130.1, 129.6, 127.3, 127.0, 126.4, 123.2, 122.0, 121.9, 119.8, 45.5, 44.4, 44.2, 41.4, 41.0, 40.8, 23.0.

HRMS ($ESI^+$) $C_{24}H_{27}N_5O_4SCl$ $[M + H]^+$ calculated mass = 516.1472, found 516.1480 (Δ = 1.5 ppm).

**Synthesis of** 1-((1*H*-indol-3-yl)methyl)-3-(2-((4-(2-chloroacetyl)piperazin-1-yl)sulfonyl)-3-methylphenyl)urea (**16**)

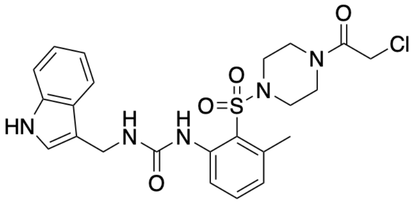

Triphosgene (61 mg, 0.206 mmol, 0.33 eq) was dissolved in DCM (1 ml), cooled on ice, then *tert*-butyl 4-((2-amino-6-methylphenyl)sulfonyl)piperazine-1-carboxylate (220 mg, 0.620 mmol, 1 eq) and pyridine (48.9 mg, 50 μL, 0.620 mmol, 1 eq) in DCM (2.435 ml) were added dropwise. The reaction was allowed to warm to room temperature and stirred for 2 h. The reaction was then cooled on ice, and (1H-indol-3-yl)methanamine (94.4 mg, 0.647 mmol, 1.05 mmol) and pyridine (48.9 mg, 50 μL, 0.620 mmol, 1 eq) in DCM (2.435 ml) was added dropwise. The reaction was allowed to warm to room temperature, then stirred overnight. The reaction was quenched via the addition of 1M $NaHCO_3$ solution (20 mL) which was subsequently stirred for 30 mins. The reaction was next diluted with DCM (10 mL), washed with $H_2O$ (20 mL) and brine (20 mL), then dried over $MgSO_4$ and purified via flash column chromatography (0 - 10% MeOH in DCM) to yield *tert*-butyl 4-((2-(3-((1H-indol-3-yl)methyl)ureido)-6-methylphenyl)sulfonyl)piperazine-1-carboxylate (253 mg, 0.417 mmol, 67%) as a pale yellow oil. $R_f$ = 0.2 (50% DCM in EtOAc). This compound was used in the next step without further characterisation.

*Tert*-butyl 4-((2-(3-((1H-indol-3-yl)methyl)ureido)-6-methylphenyl)sulfonyl)piperazine-1-carboxylate (220 mg, 0.417 mmol, 1 eq) was add to 5% TFA (*v/v*) in DCM (4.17 mL, 0.1 M), then stirred at room temperature for 4 h. Water (5 mL) was added, then the solution basified via the dropwise addition of 3 M NaOH until it reached pH 12. The free amine was then extracted with DCM (10 mL), washed with brine (10 mL), dried over $MgSO_4$ and used in the next step without characterisation.

1-((1H-indol-3-yl)methyl)-3-(2-((4-(2-chloroacetyl)piperazin-1-yl)sulfonyl)-3-methylphenyl)urea (127 mg, 0.297 mmol) was dissolved in DCM (0.86 mL, 0.35 M) then chloroacetyl chloride added (33.5 mg, 23.6 μL, 0.297 mmol, 1 eq) and the reaction stirred for 10 mins at room temperature. Amberlyst resin OH form (80.3 mg) was subsequently added and the reaction stirred for a further 10 mins. The reaction mixture was filtered through celite, then purified via flash column chromatography (0 – 10% MeOH in DCM) to yield 1-((1*H*-indol-3-yl)methyl)-3-(2-((4-(2-chloroacetyl)piperazin-1-yl)sulfonyl)-3-methylphenyl)urea (**16**, 23 mg, 0.046 mmol, 15%) as a pale yellow oil. $R_f$ = 0.4 (50% EtOAc in DCM).

$^1$H NMR ($CDCl_3$, 400 MHz) δ 8.91 (1H, s, Ar-N*H*CH), 8.46 (1H, s, CON*H*Ar), 8.07 (1H, d, Ar-*H*), 7.42 (1H, t, Ar-*H*), 7.29 (1H, d, Ar-*H*), 7.15 (1H, t, Ar-*H*), 7.11 (1H, t, Ar-*H*), 7.05 (1H, d, Ar-

*H*), 6.97 (1H, d, Ar-*H*), 6.63 (1H, s, CON*H*CH$_2$), 4.74 (2H, d, CONHC*H$_2$*), 3.92 (COC*H$_2$*Cl), 3.24 (4H, s (br), C*H$_2$*NCO), 3.04 (2H, s (br), C*H$_2$*NSO$_2$), 2.88 (2H, s (br), C*H$_2$*NSO$_2$), 2.58 (3H, s, Ar-C*H$_3$*).

$^{13}$C NMR (CDCl$_3$, 101 MHz) δ 165.2, 154.7, 140.4, 140.2, 136.1, 133.7, 130.3, 127.4, 126.7, 124.7, 122.3, 122.2, 119.5, 111.1, 100.9, 45.6, 44.5, 44.4, 43.1, 41.3, 40.8, 23.2.

HRMS (ESI$^+$) $C_{24}H_{27}N_5O_4SCl$ [M + H]$^+$ calculated mass = 504.1472, found 504.1466 (Δ = 1.2 ppm).

**Synthesis of** *tert*-butyl 4-((5-bromo-2-nitrophenyl)sulfonyl)piperazine-1-carboxylate

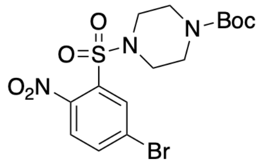


5-Bromo 2-nitro benzenesulfonyl chloride (500 mg, 1.7 mmol, 1 equiv) and DIPEA (440 μL, 326.5 mg, 2.55 mmol, 1.5 equiv) were dissolved in DCM (5.7 mL, 0.3 M) and cooled on ice. 1-boc piperazine (316.6 mg, 1.7 mmol, 1 equiv) was then added, and the reaction allowed to warm to room temperature and stirred for 30 mins. The reaction mixture was subsequently diluted with DCM, washed with 1 M HCl (2 x 20 mL), water (20 mL) and brine (20 mL), then dried over $MgSO_4$ and solvent removed *in vacuo.* Purification via flash column chromatography (50% DCM in hexane) yielded *tert*-butyl 4-((5-bromo-2-nitrophenyl)sulfonyl)piperazine-1-carboxylate (344 mg, 0.77 mmol, 45%) as a colourless oil. $R_f$ = 0.34 (100% DCM).

$^1$H NMR (CDCl$_3$, 400 MHz) δ 8.10 (1H, d, J = 2.0 Hz, Ar-*H*), 7.85 (1H, dd, J = 2.1, 8.3 Hz, Ar-*H*), 7.54 (1H, d, J = 8.4 Hz, Ar-*H*), 3.53 (4H, t, 5.0 Hz, C*H$_2$*NCO), 3.30 (4H, t, J = 5.4 Hz, C*H$_2$*NSO$_2$), 1.44 (9H, s, C(C*H$_3$*)$_3$).

$^{13}$C NMR (CDCl$_3$, 101 MHz) δ 154.5, 147.3, 137.2, 134.1, 133.4, 126.1, 81.0, 46.2, 28.9.

HRMS (ESI$^+$) $C_{15}H_{20}N_3O_6SBr$ [M + H]$^+$ calculated mass = 448.0183, found 403.0190 (Δ = 1.6 ppm).

**Synthesis of** *tert*-butyl 4-((5-ethynyl-2-nitrophenyl)sulfonyl)piperazine-1-carboxylate

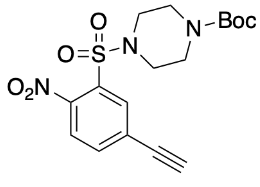

*Tert*-butyl 4-((5-bromo-2-nitrophenyl)sulfonyl)piperazine-1-carboxylate (114 mg, 0.288 mmol, 1 equiv) , TMS-acetylene (37.3 mg, 0.380 mmol, 1.3 equiv), CuI (2.4 mg, 0.012 mmol, 4 mol%), $Pd(Ph_3)_4$ (7.2 mg, 0.0062 mmol, 2 mol%) and $NEt_3$ (75.5 mg, 104 μL, 0.746 mmol, 2.6 equiv) were combined in DMF (1.25 ml, 0.23 M) under an argon atmosphere and heated to 80°C overnight. The reaction mixture was next cooled on ice, and TBAF (78.45 mg, 0.3 mmol, 1.05 equiv) in THF (1.875 ml) added dropwise. The reaction mixture was suspended in EtOAc (10 mL), then washed with water (3 x 50 mL) and brine (10 mL), then dried over $MgSO_4$ and solvent removed *in vacuo.* Purification via flash column chromatography (40% EtOAc in pentane) yielded *tert*-butyl 4-((5-ethynyl-2-nitrophenyl)sulfonyl)piperazine-1-carboxylate (62 mg, 0.16 mmol, 55%) as an orange/yellow oil. $R_f$ = 0.35 (100% DCM).

$^1$H NMR ($CDCl_3$, 400 MHz) δ 8.03 (1H, d, J = 1.7 Hz, Ar-*H*), 7.76 (1H, dd, J = 1.8, 8.2 Hz, Ar-*H*), 7.59 (1H, d, J = 8.2 Hz, Ar-*H*), 3.51 (4H, t, 5.1 Hz, C*H₂*NCO), 3.38 (1H, s, Ar-C*H*), 3.28 (4H, t, J = 5.3 Hz, $CH_2NSO_2$), 1.43 (9H, s, $C(CH_3)_3$).

$^{13}$C NMR ($CDCl_3$, 101 MHz) δ 154.3, 147.5, 137.0, 134.4, 131.9, 126.7, 124.5, 83.0, 80.7, 80.1, 46.0, 28.4.

HRMS (ESI$^+$) $C_{17}H_{21}N_3O_6S$ $[M + H]^+$ calculated mass = 395.1157, found 395.1158 (Δ = 0.3 ppm).

**Synthesis of** *tert*-butyl 4-((2-amino-5-ethynylphenyl)sulfonyl)piperazine-1-carboxylate

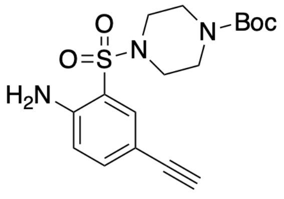


*Tert*-butyl 4-((2-amino-5-ethynylphenyl)sulfonyl)piperazine-1-carboxylate (60.3 mg, 0.15 mmol, 1 equiv), Fe powder (41.8 mg, 0.76 mmol, 5 equiv) and $NH_4Cl$ (65.2 mg, 1.22 mmol, 8 equiv) were combined in EtOH (0.6 mL) / $H_2O$ (0.15 mL) then stirred at 80°C for 4 h. The reaction mixture was filtered, then extracted with DCM (2 x 10 mL). The combined organic extracts were washed with water (2 x 20 mL) and brine (20 mL), then dried over $MgSO_4$, yielding *tert*-butyl 4-((2-amino-5-ethynylphenyl)sulfonyl)piperazine-1-carboxylate (55 mg, 0.15 mmol, 99%) as a yellow oil. $R_f$ = 0.3 (100% DCM).

$^1$H NMR ($CDCl_3$, 400 MHz) δ 7.68 (1H, s, Ar-*H*), 7.39 (1H, s, Ar-*H*), 6.66 (1H, s, Ar-*H*), 5.26 (2H, s, $NH_2$), 3.48 (4H, s, $CH_2NCO$), 3.07 (4H, s, $CH_2NSO_2$), 1.41 (9H, s, $C(CH_3)_3$).

$^{13}$C NMR ($CDCl_3$, 101 MHz) δ 154.8, 149.3, 140.7, 133.9, 121.9, 116.8, 80.8, 45.1, 28.8, 23.1.

HRMS (ESI$^+$) $C_{16}H_{26}N_3O_4S$ $[M + H]^+$ calculated mass = 356.1644, found 356.1646 (Δ = 0.6 ppm).

**Synthesis of –** *tert*-butyl 4-((2-(3-((1H-indol-3-yl)methyl)ureido)-5-ethynylphenyl)sulfonyl)piperazine-1-carboxylate

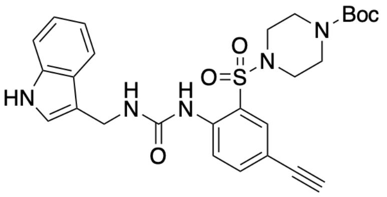

Triphosgene (16.5 mg, 0.056 mmol, 0.37 equiv) was dissolved in DCM (100 μL), cooled on ice, then *tert*-butyl 4-((2-amino-5-ethynylphenyl)sulfonyl)piperazine-1-carboxylate (55 mg, 0.15 mmol, 1 equiv) and pyridine (12.1 μL, 11.9 mg, 0.15 mmol, 1 equiv) in DCM (0.5 mL, 0.3 M) were added dropwise. The reaction was allowed to warm to room temperature and stirred for 2 h. The reaction was then cooled on ice, and (1H-indol-3-yl)methanamine (35.1 mg, 0.24 mmol, 1.6 equiv) and pyridine (12.1 μL, 11.9 mg, 0.15 mmol, 1 equiv) in DCM (0.5 mL) was added dropwise. The reaction was allowed to warm to room temperature, then stirred overnight. The reaction was quenched via the addition of 1M $NaHCO_3$ solution (10 mL) which was subsequently stirred for 30 mins. The reaction was next diluted with DCM (5 mL), washed with $H_2O$ (10 mL), HCl (10 mL) and brine (20 mL), then dried over $MgSO_4$ and purified via flash column chromatography (0-30% gradient of EtoAc in DCM) to yield *tert*-butyl 4-((2-(3-((1H-indol-3-yl)methyl)ureido)-5-ethynylphenyl)sulfonyl)piperazine-1-carboxylate (35.7 mg, 0.066 mmol, 44%) as a white solid. $R_f$ = 0.2 (10% EtOAc in DCM).

$^1$H NMR ($CDCl_3$, 400 MHz) δ 8.68 (1H, s (br), Ar-N*H*), 8.46 (1H, s (br), Ar-N*H*CH), 8.38 (1H, d, J = 8.8 Hz, Ar-*H*), 7.76 (1H, s, Ar-*H*), 7.60 (1H, dd, J = 8.8, 2.1 Hz, Ar-*H*), 7.34 (1H, d, J = 8.0 Hz, Ar-*H*), 7.22 (1H, t, J = 3.2 Hz, Ar-*H*), 7.14 (1H, t, J = 7.2 Hz, Ar-*H*), 7.04 (1H, J = 7.2 Hz, Ar-*H*), 6.60 (1H, s (br), Ar-$CH_2$N*H*), 5.30 (1H, s, ArNHC*H*), 4.75 (2H, d, J = 5.5 Hz, CONHC$H_2$), 3.34 (4H, s (br), C$H_2$NCO), 3.10 (1H, s, Ar-C*H*), 2.87 (4H, s (br), C$H_2$N$SO_2$), 1.44 (9H, s, C(C$H_3$)$_3$).

$^{13}$C NMR ($CDCl_3$, 101 MHz) δ 153.8, 153.5, 138.4, 137.5, 135.8, 133.0, 126.3, 124.4, 121.9, 121.8, 121.5, 119.0, 115.7, 110.9, 100.4, 81.6, 80.4, 76.5, 45.3, 42.8, 28.1.

HRMS (ESI$^+$) $C_{27}H_{32}N_5O_5S$ $[M + H]^+$ calculated mass = 538.2124, found 538.2119 (Δ = 0.9 ppm).

**Synthesis of** 1-((1*H*-indol-3-yl)methyl)-3-(2-((4-(2-chloroacetyl)piperazin-1-yl)sulfonyl)-4-ethynylphenyl)urea (**17**)

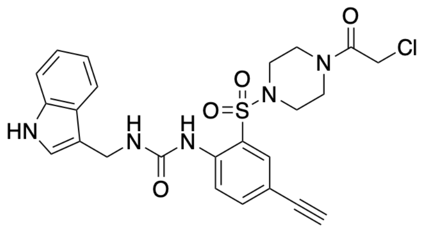

4-((2-(3-((1H-indol-3-yl)methyl)ureido)-5-ethynylphenyl)sulfonyl)piperazine-1-carboxylate (35.7 mg, 0.066 mmol, 1 equiv) was added to 5% TFA (*v/v*) in DCM (2 mL, 0.033 M), then stirred at room temperature for 3 h, in which time the reaction changed from colourless to dark red. Water (10 mL) was added, then the solution basified via the dropwise addition of 3 M NaOH until it reached pH 12. The free amine was then extracted with DCM (2 x 10 mL), washed with brine (10 mL), dried over $MgSO_4$ and used in the next step without characterisation.

1-((1H-indol-3-yl)methyl)-3-(4-ethynyl-2-(piperazin-1-ylsulfonyl)phenyl)urea (24.4 mg, 0.055 mmol, 1 equiv) was dissolved in DCM (0.279 mL, 0.2 M) then chloroacetyl chloride added (4.4 mL, 4.31 mg, 0.055 mmol, 1 equiv) and the reaction stirred for 10 mins at room temperature. Amberlyst resin OH form (15 mg) was subsequently added and the reaction stirred for a further 10 mins. The reaction mixture was filtered through celite, then purified via flash column chromatography (50% EtOAc in hexane) to yield 1-((1H-indol-3-yl)methyl)-3-(2-((4-(2-chloroacetyl)piperazin-1-yl)sulfonyl)-4-ethynylphenyl)urea (2.5 mg, 0.0049 mmol, 7%) as a colourless oil. $R_f$ = 0.2 (20% EtOAc in DCM).

$^1$H NMR ($CDCl_3$, 400 MHz) δ 8.54 (1H, s (br), ArN*H*), 8.39 (1H, s (br), Ar-N*H*CH), 8.33 (1H, d, J = 8.8 Hz, Ar-*H*), 7.79 (1H, d, Ar-*H*), 7.64 (1H, dd, J = 8.7, 2.1 Hz, Ar-*H*), 7.37 (1H, d, J = 7.9 Hz, Ar-*H*), 7.24 (1H, t, J = 3.3 Hz, Ar-*H*), 7.17 (1H, t, J = 7.1 Hz, Ar-*H*), 7.06 (1H, J = 7.0 Hz, Ar-*H*), 6.63 (1H, s (br), Ar-$CH_2$N*H*), 5.23 (1H, s, ArNHC*H*), 4.75 (2H, d, J = 5.8 Hz, CONHC$H_2$), 3.94 (2H, s, COC$H_2$Cl), 3.40 (4H, m, C$H_2$NCO), 3.11 (1H, s, Ar-C*H*), 2.93 (4H, m, C$H_2$$NSO_2$).

$^{13}$C NMR ($CDCl_3$, 101 MHz) δ 165.1, 153.7, 138.4, 137.8, 136.0, 133.2, 129.1, 128.2, 125.3, 124.7, 122.9, 122.4, 122.1, 119.4, 116.4, 111.1, 100.7, 81.6, 78.4, 77.3, 77.0, 76.8, 45.3, 43.1, 41.2, 40.6, 21.5.

HRMS (ESI$^+$) $C_{24}H_{27}N_5O_4SCl$ $[M + H]^+$ calculated mass = 514.1316, found 514.1326 (Δ = 1.9 ppm).

#### 2.4 Analytical Data for Reported Compounds

##### $^1$H-NMR (400 MHz, $CDCl_3$)

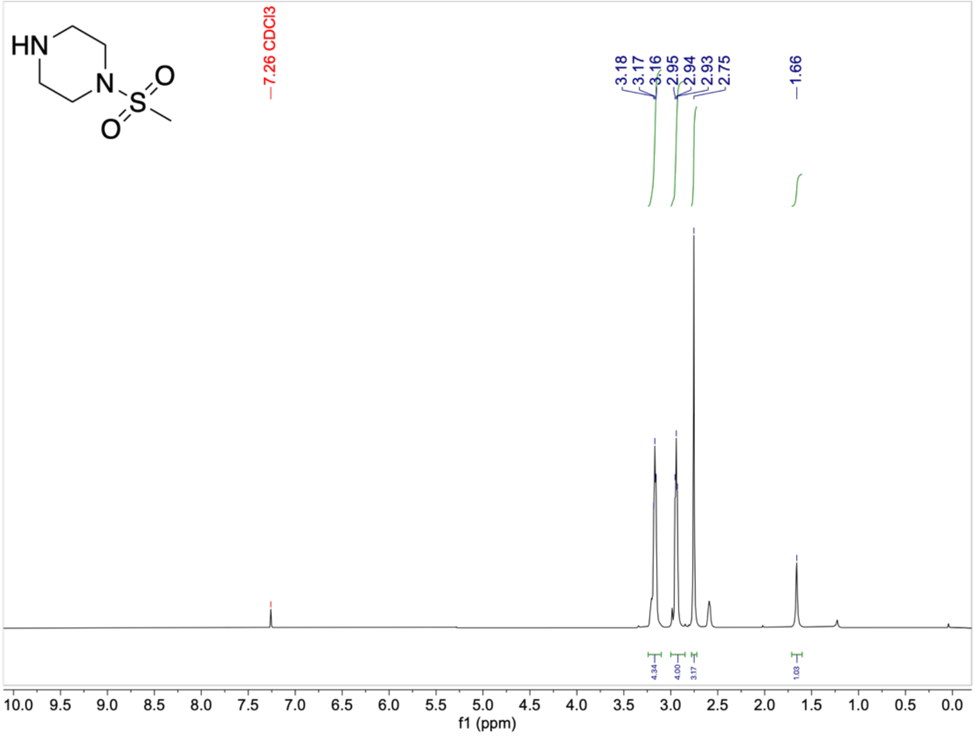


##### $^{13}$C-NMR (101 MHz, $CDCl_3$)

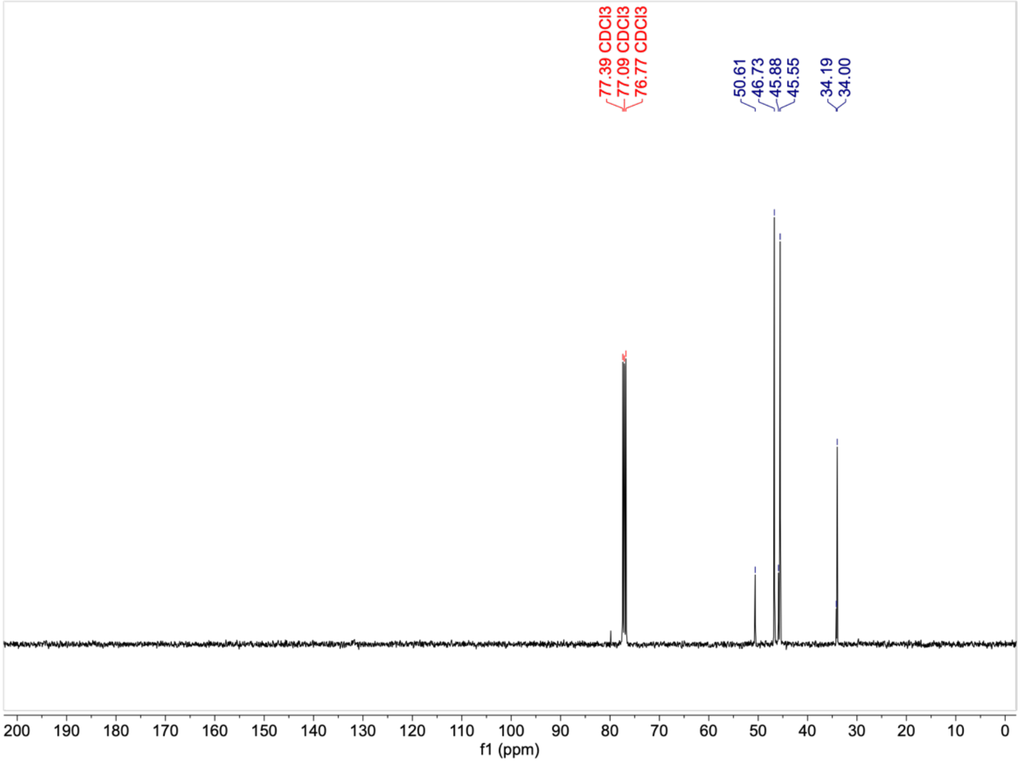

#### $^{1}$H-NMR (400 MHz, $CDCl_3$)

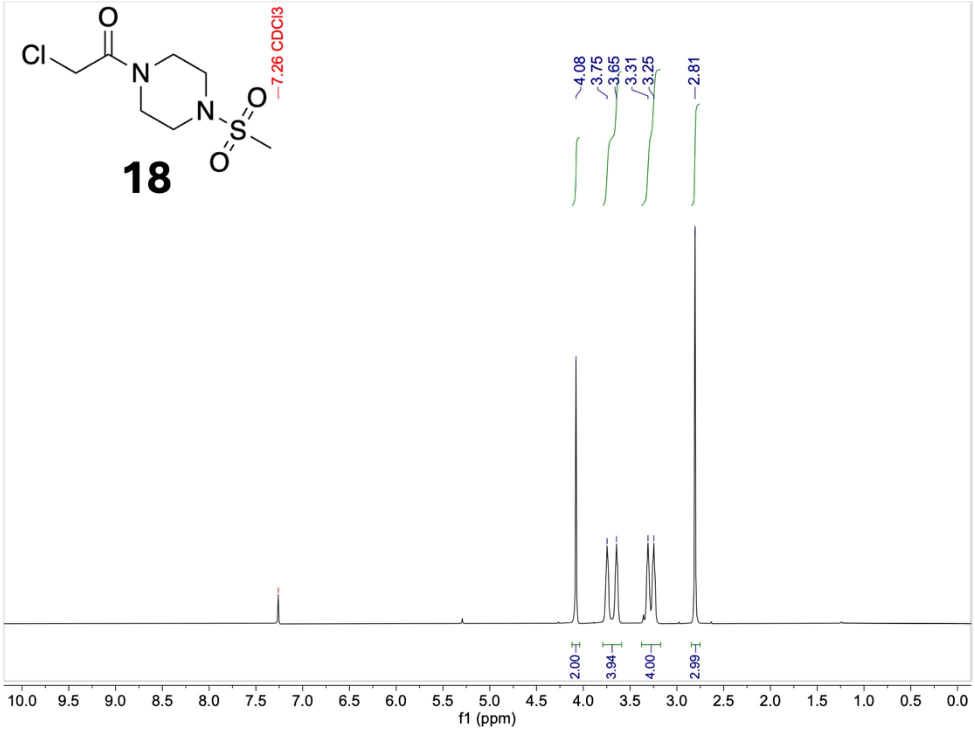


#### $^{13}$C-NMR (101 MHz, $CDCl_3$)

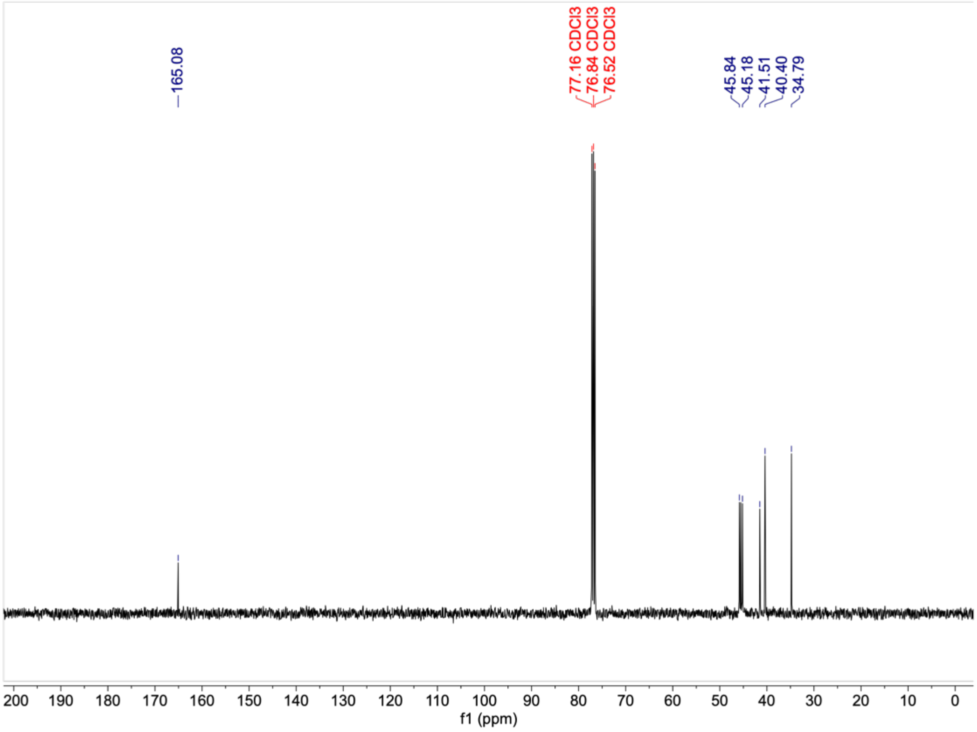

#### $^{1}$H-NMR (400 MHz, $CDCl_3$)

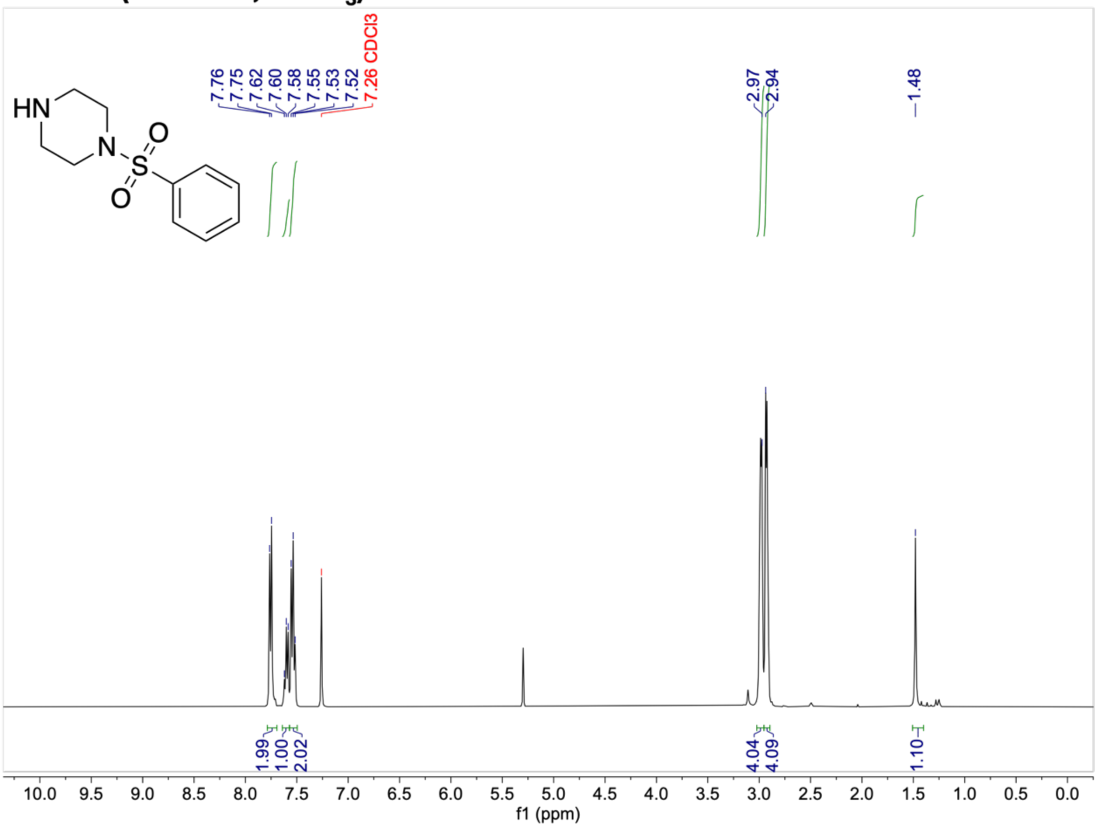


#### $^{13}$C-NMR (101 MHz, $CDCl_3$)

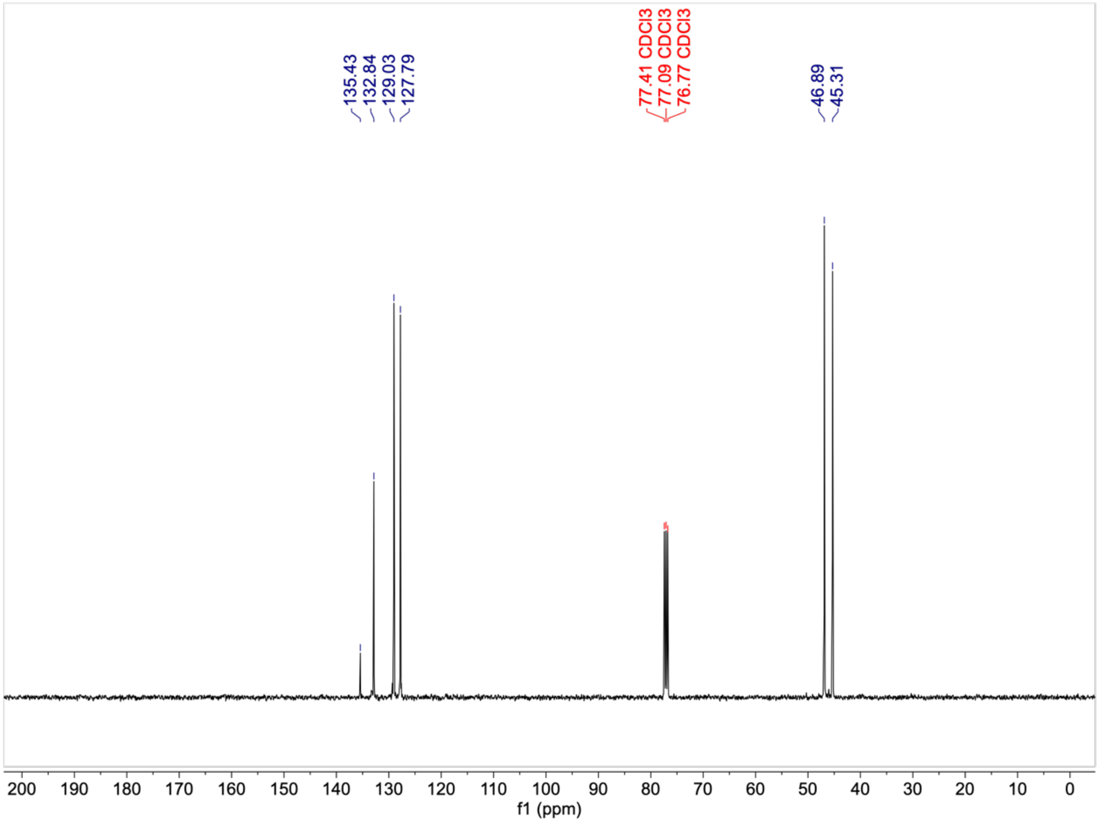

#### $^{1}$H-NMR (400 MHz, $CDCl_3$)

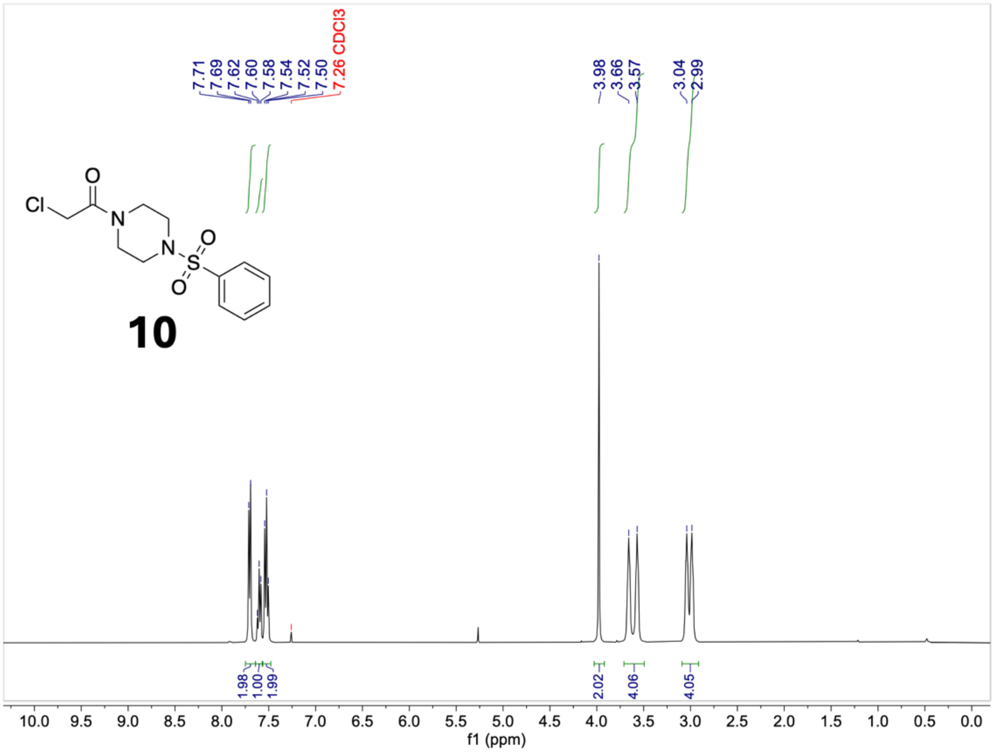


#### $^{13}$C-NMR (101 MHz, $CDCl_3$)

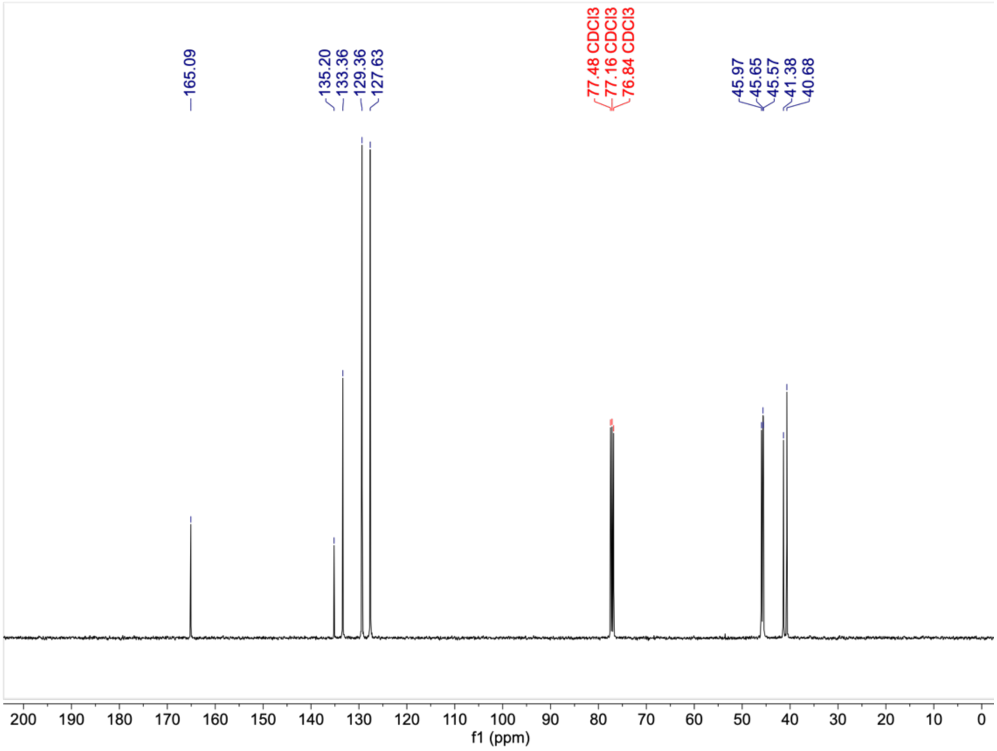

**[1]H-NMR (400 MHz, $CDCl_3$)**

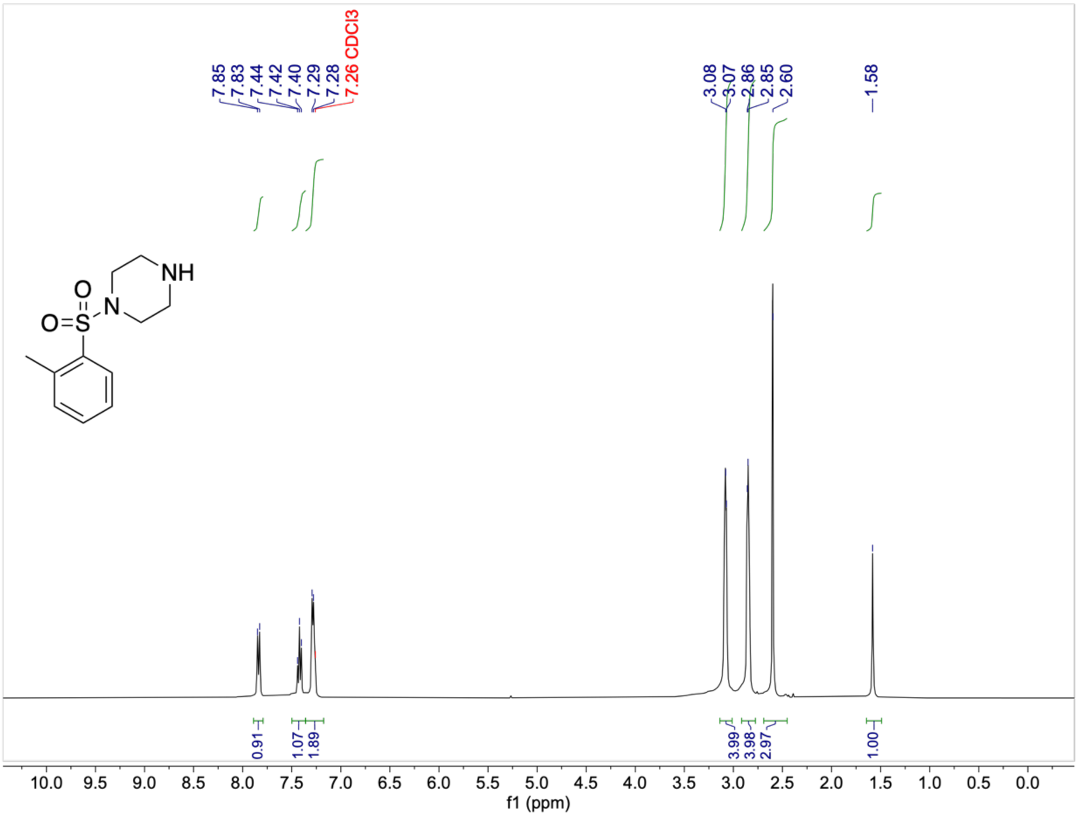


**[13]C-NMR (101 MHz, $CDCl_3$)**

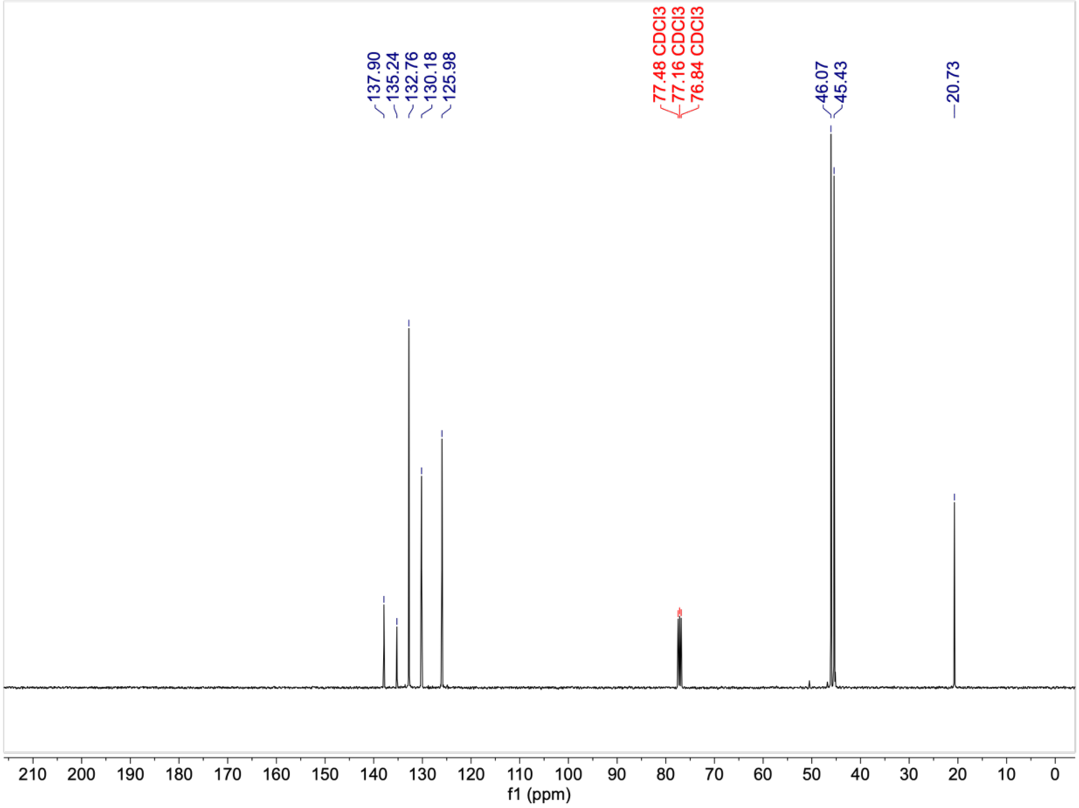

#### $^{1}H$-NMR (400 MHz, $CDCl_3$)

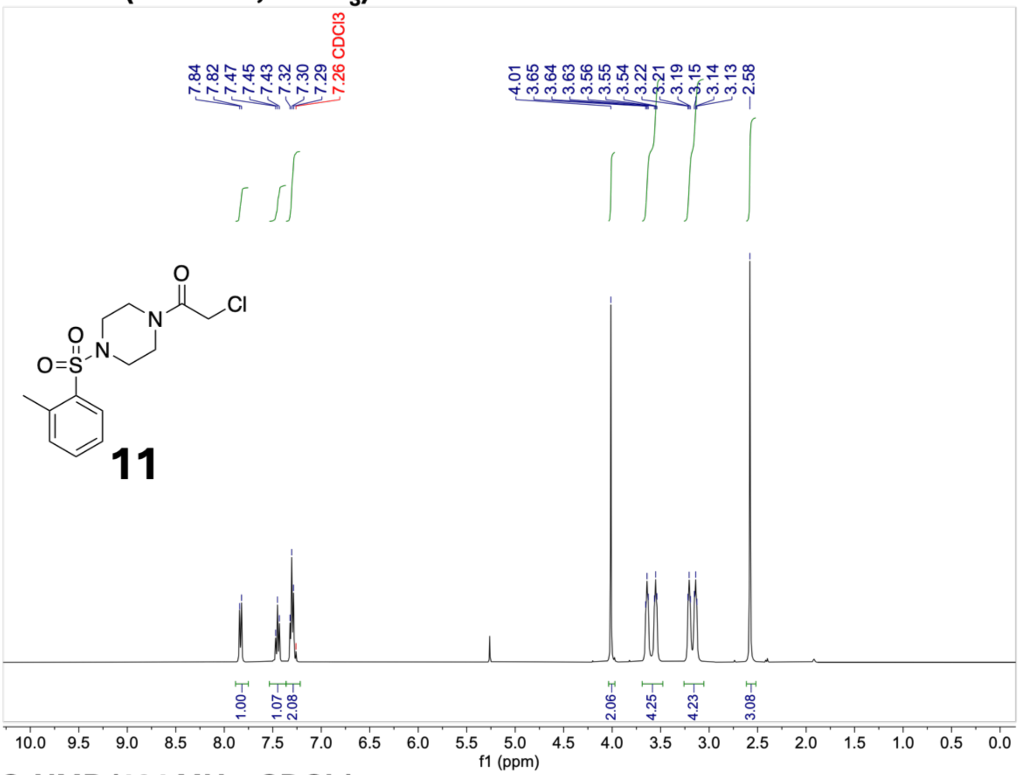


#### $^{13}C$-NMR (101 MHz, $CDCl_3$)

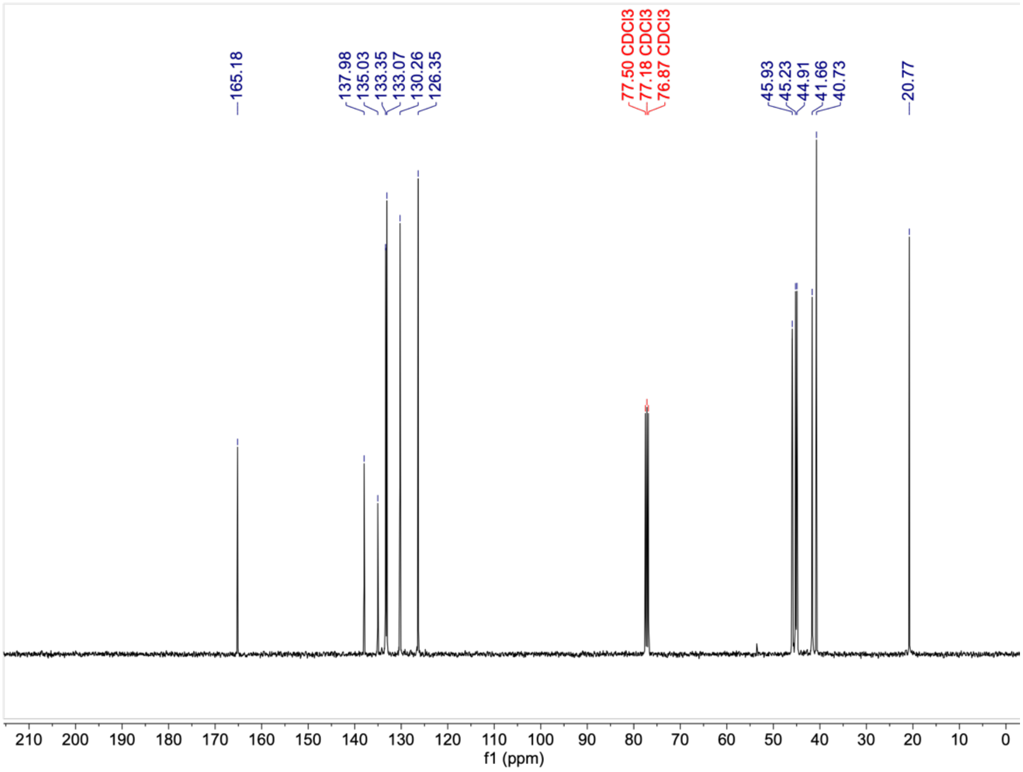

#### $^1$H-NMR (400 MHz, $CDCl_3$)

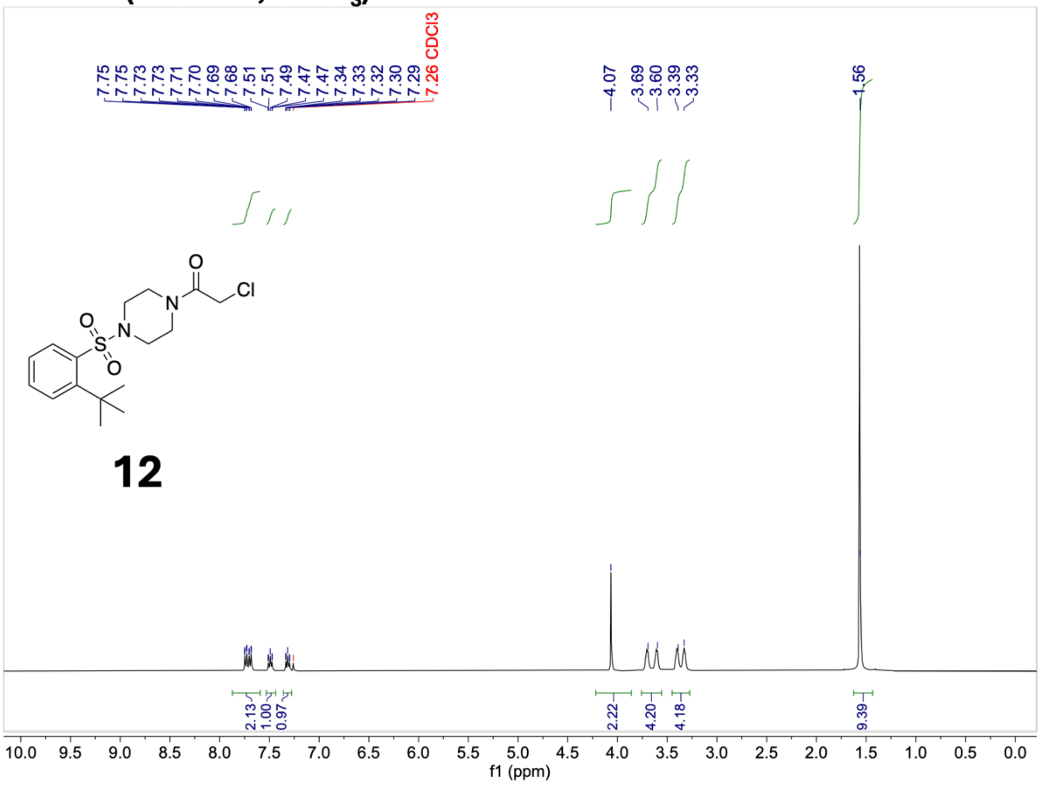


#### $^{13}$C-NMR (101 MHz, $CDCl_3$)

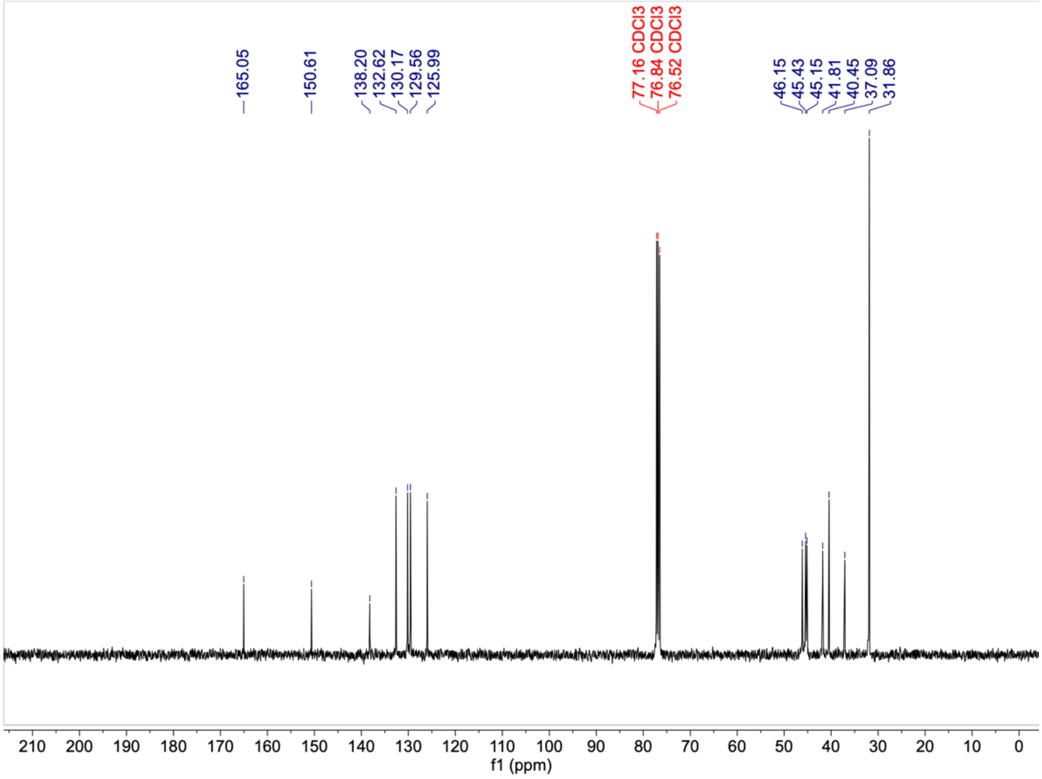

#### $^{1}$H-NMR (400 MHz, $CDCl_3$)

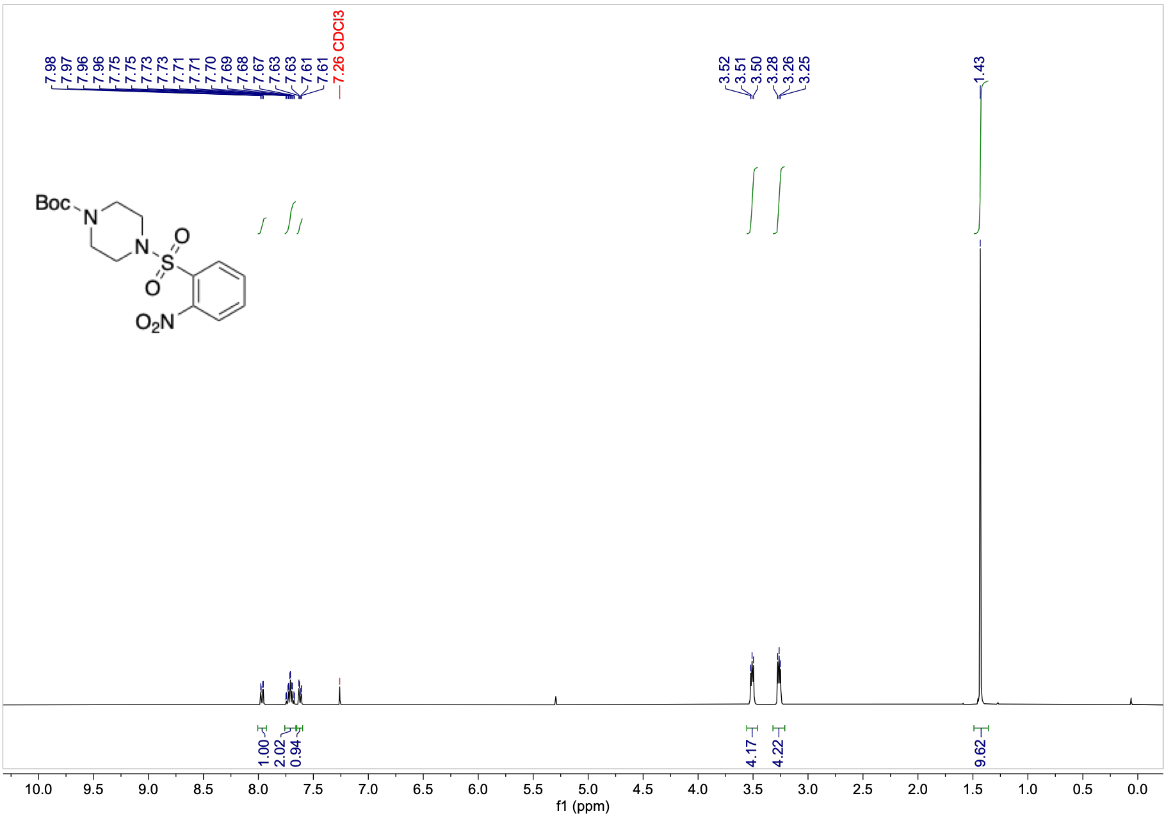


#### $^{13}$C-NMR (101 MHz, $CDCl_3$)

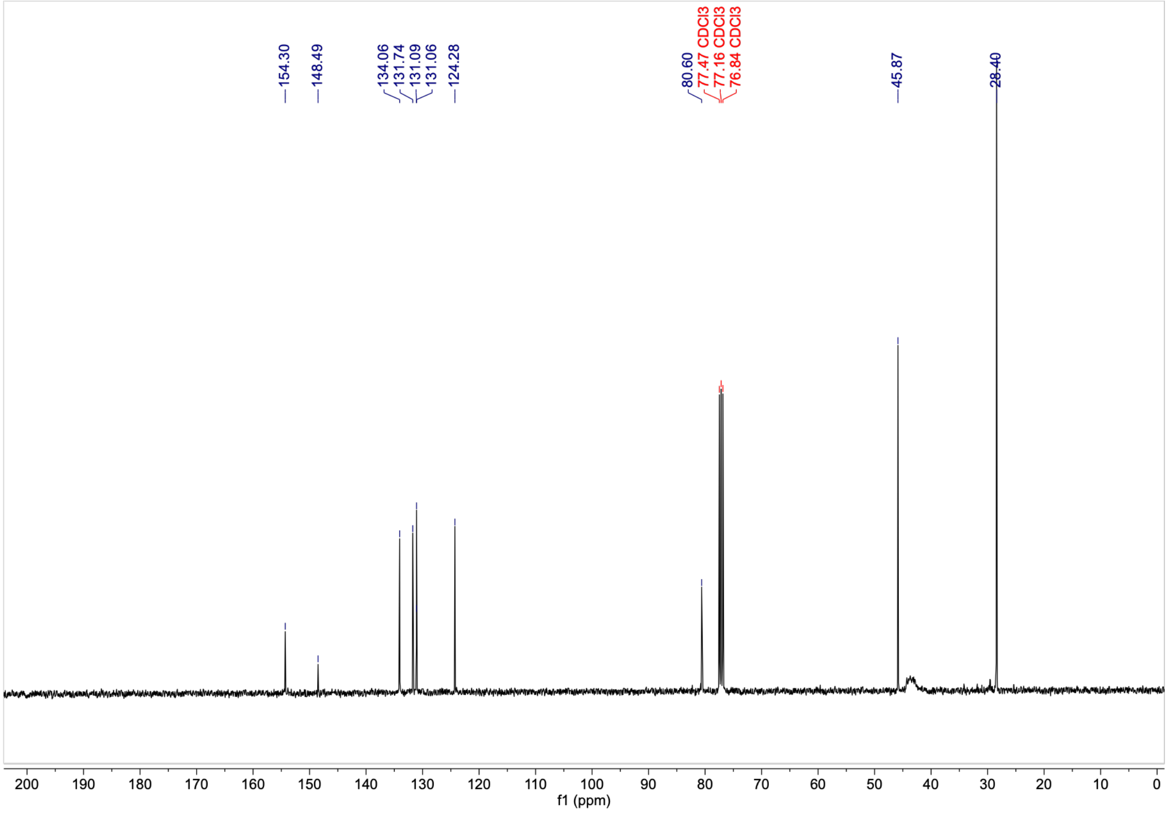

#### $^{1}$H-NMR (400 MHz, $CDCl_3$)

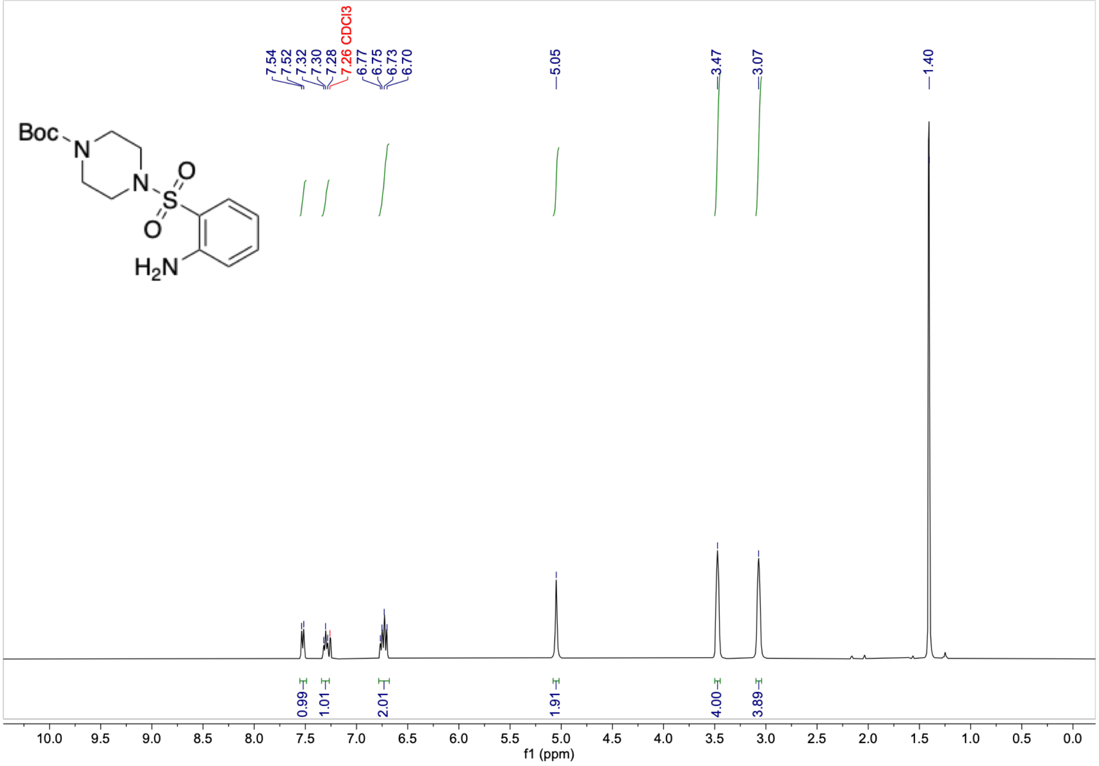


#### $^{13}$C-NMR (101 MHz, $CDCl_3$)

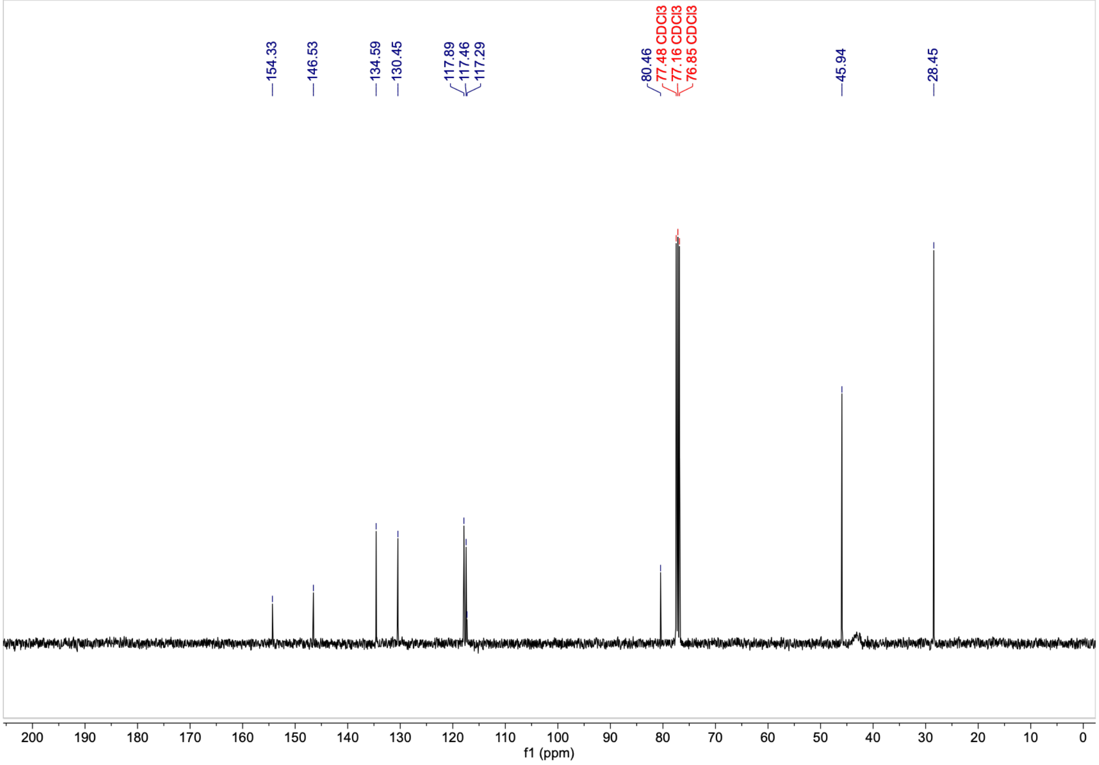

#### $^1$H-NMR (400 MHz, $CDCl_3$)

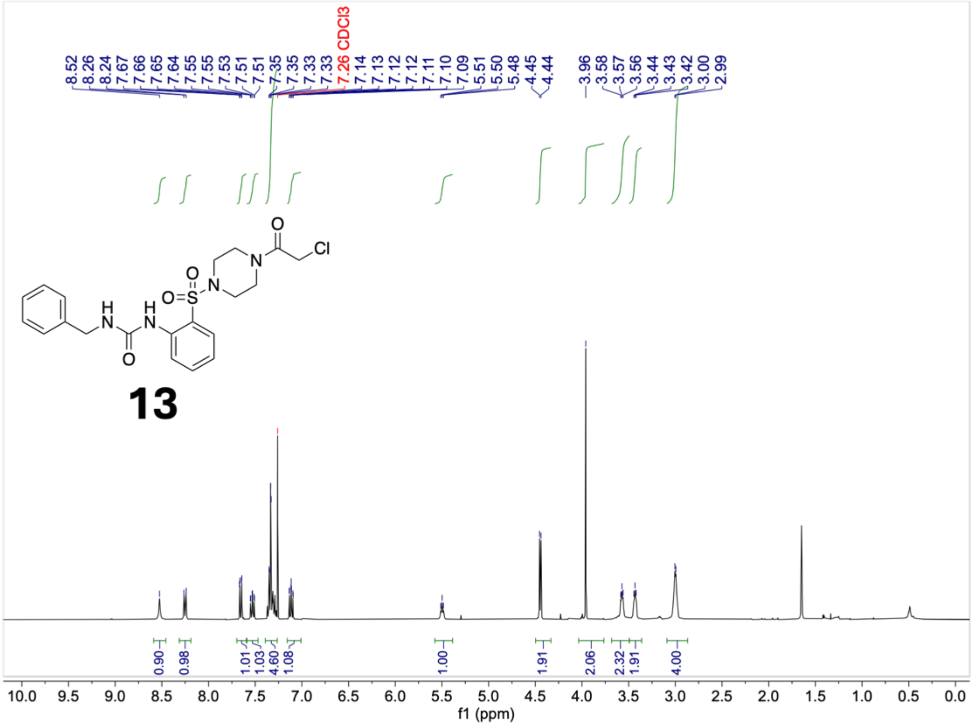


#### $^{13}$C-NMR (101 MHz, $CDCl_3$)

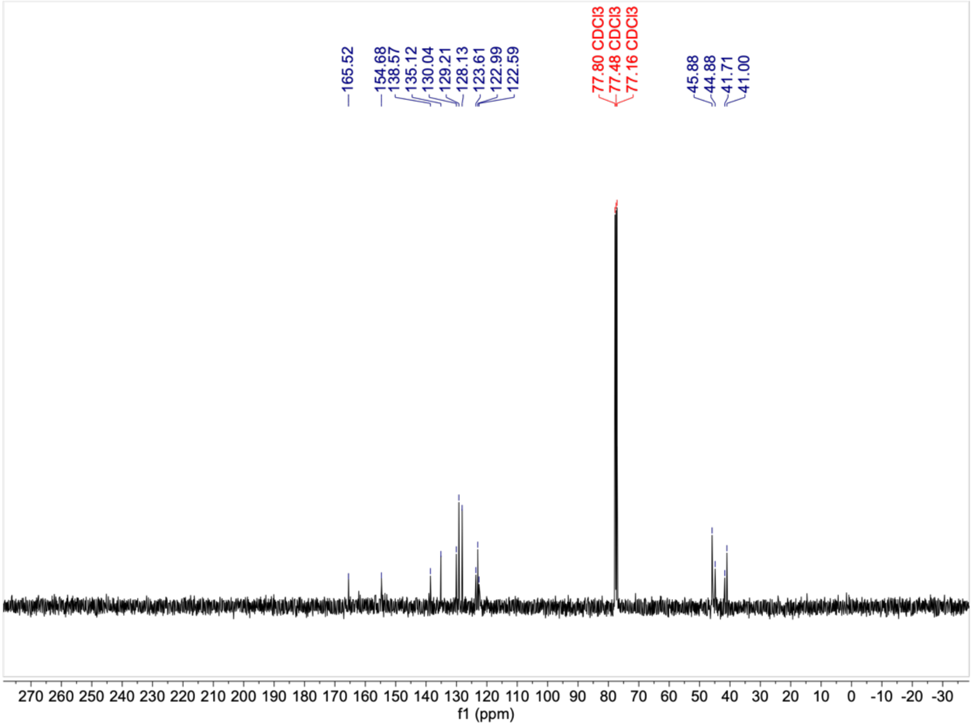

#### $^1$H-NMR (400 MHz, $CDCl_3$)

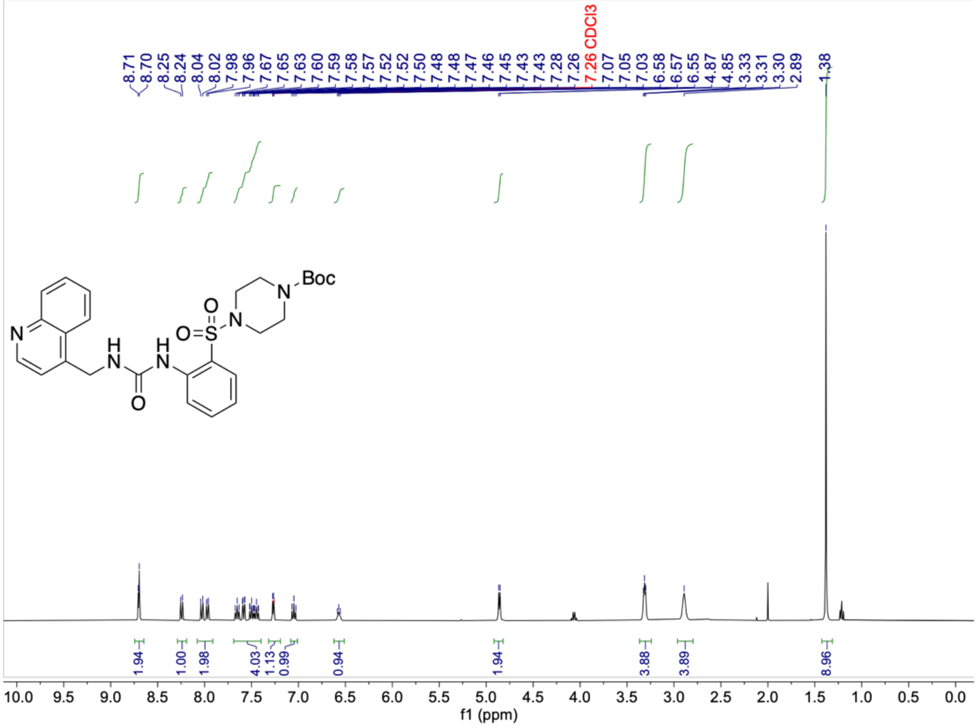

#### $^{13}$C-NMR (101 MHz, $CDCl_3$)

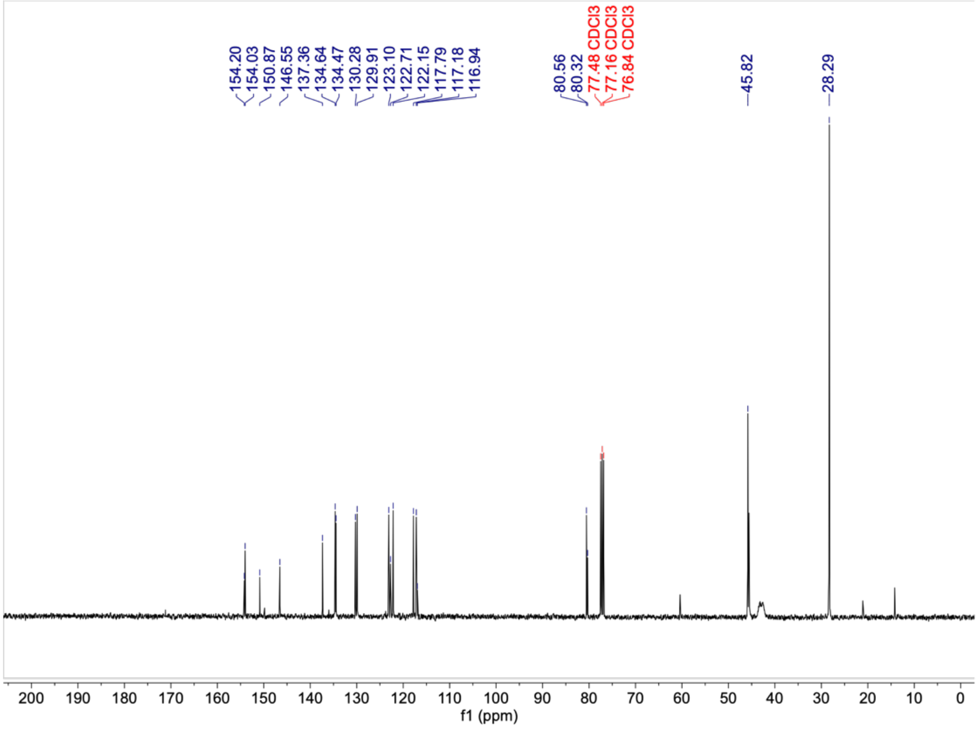

#### $^1$H-NMR (400 MHz, $CDCl_3$)

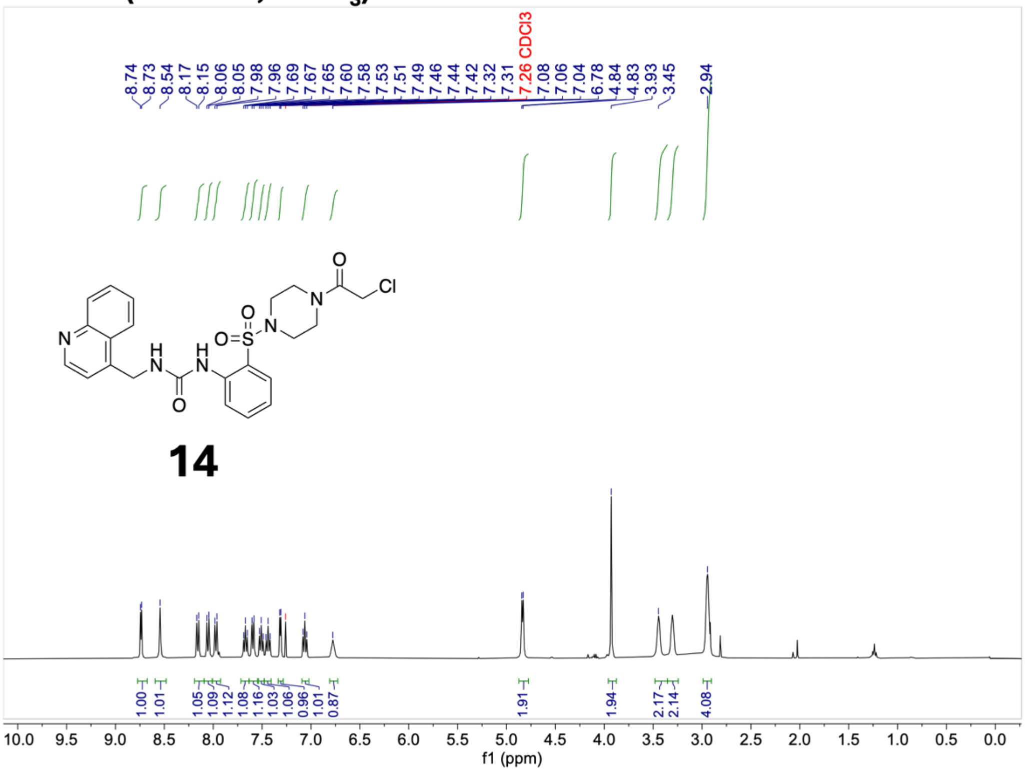


#### $^{13}$C-NMR (101 MHz, $CDCl_3$)

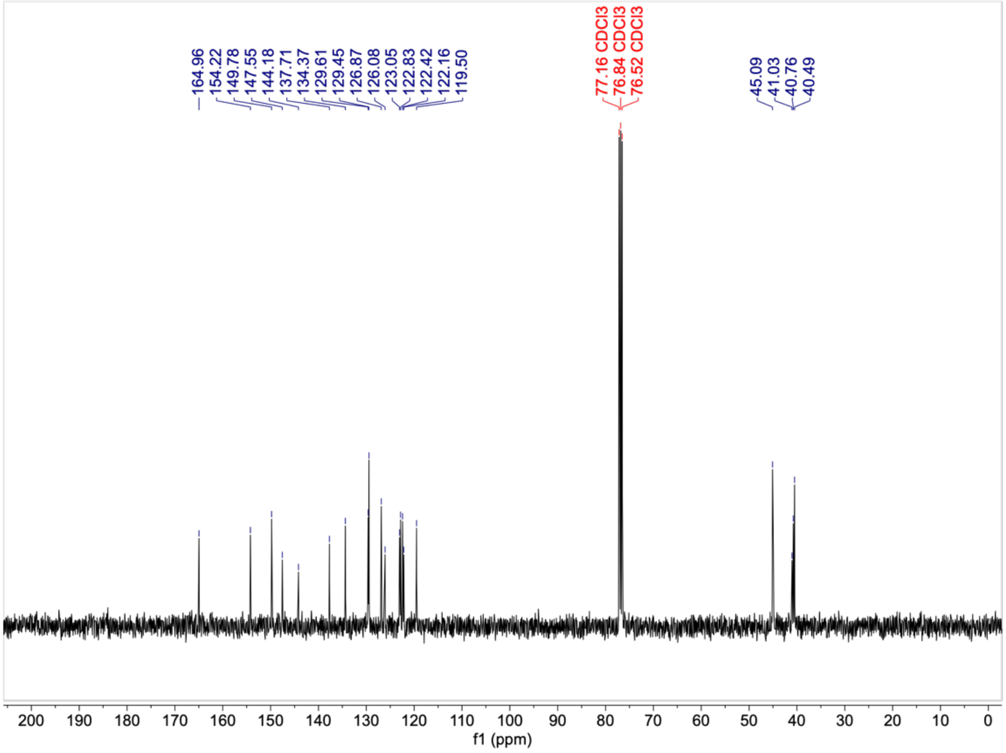

#### $^{1}$H-NMR (400 MHz, $CDCl_3$)

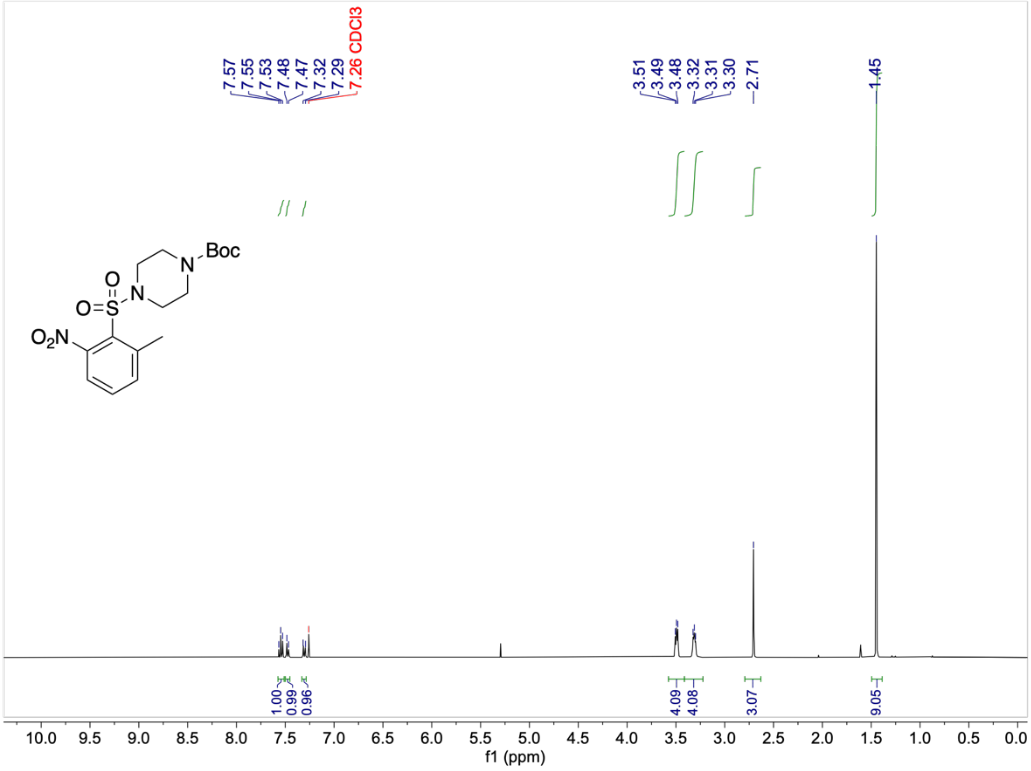


#### $^{13}$C-NMR (101 MHz, $CDCl_3$)

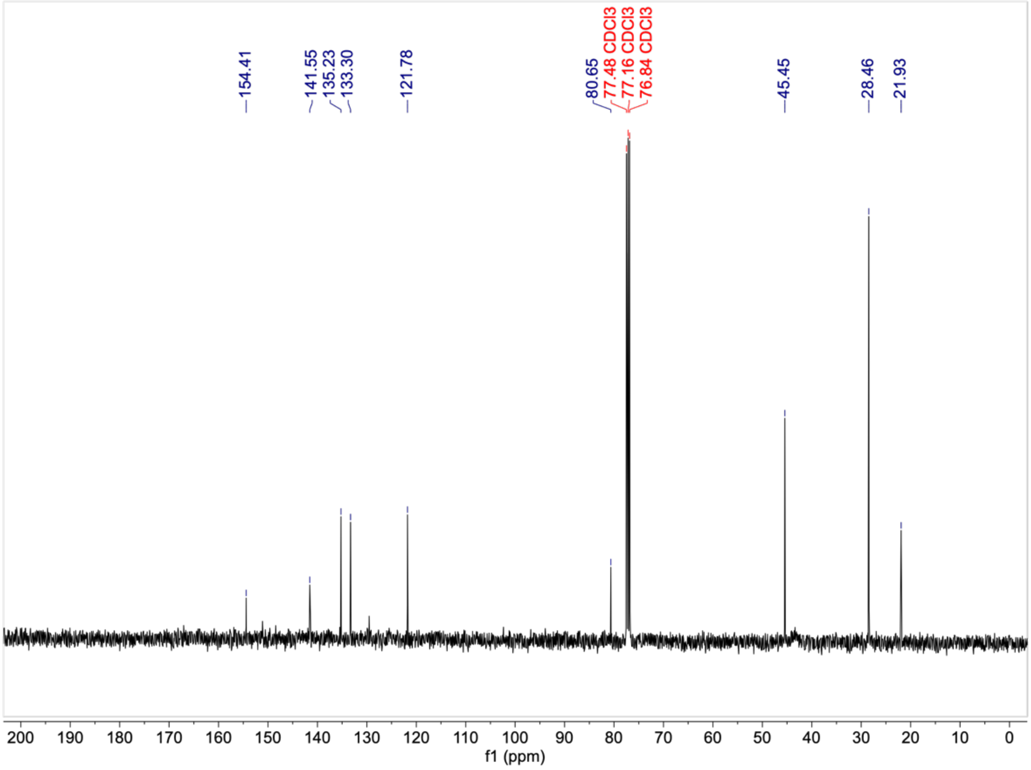

**$^1$H-NMR (400 MHz, CDCl$_3$)**

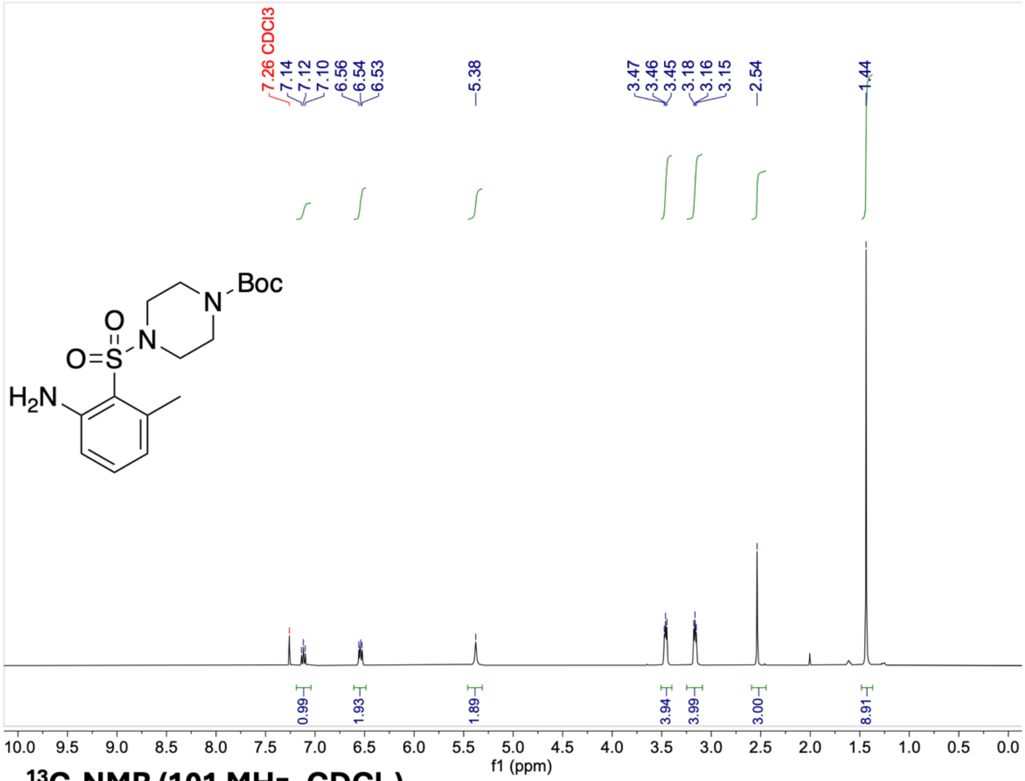


**$^{13}$C-NMR (101 MHz, CDCl$_3$)**

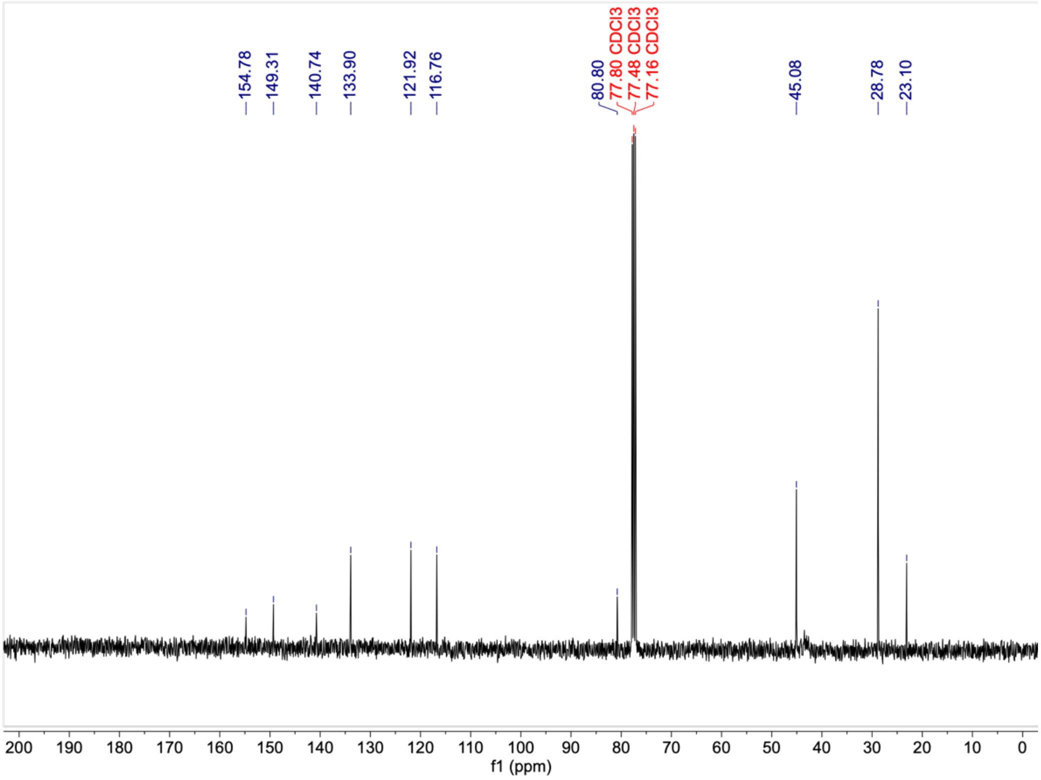

#### $^1$H-NMR (400 MHz, $CDCl_3$)

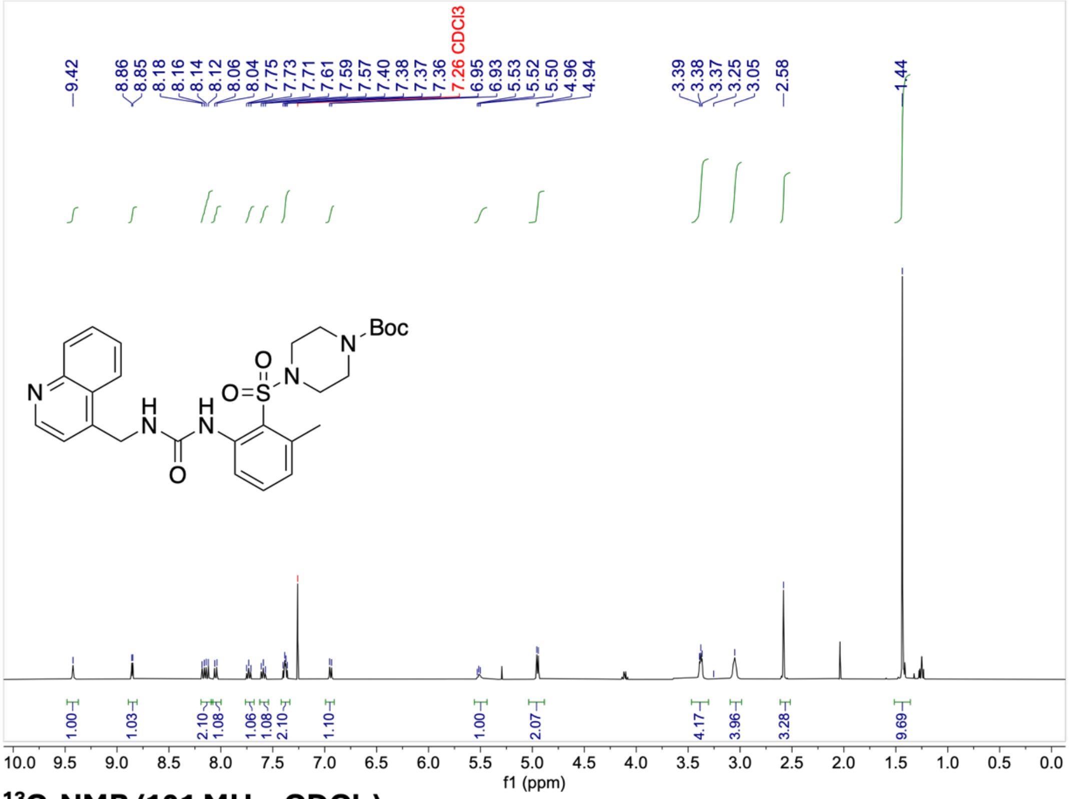

#### $^{13}$C-NMR (101 MHz, $CDCl_3$)

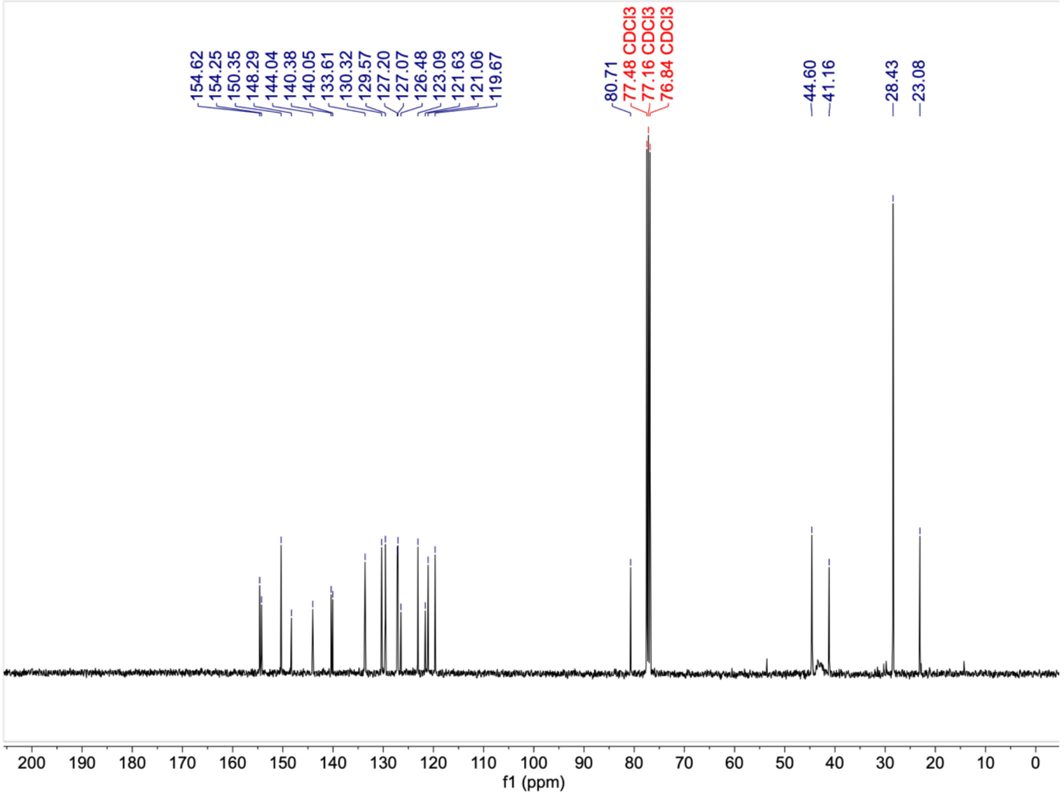

#### $^{1}$H-NMR (400 MHz, $CDCl_3$)

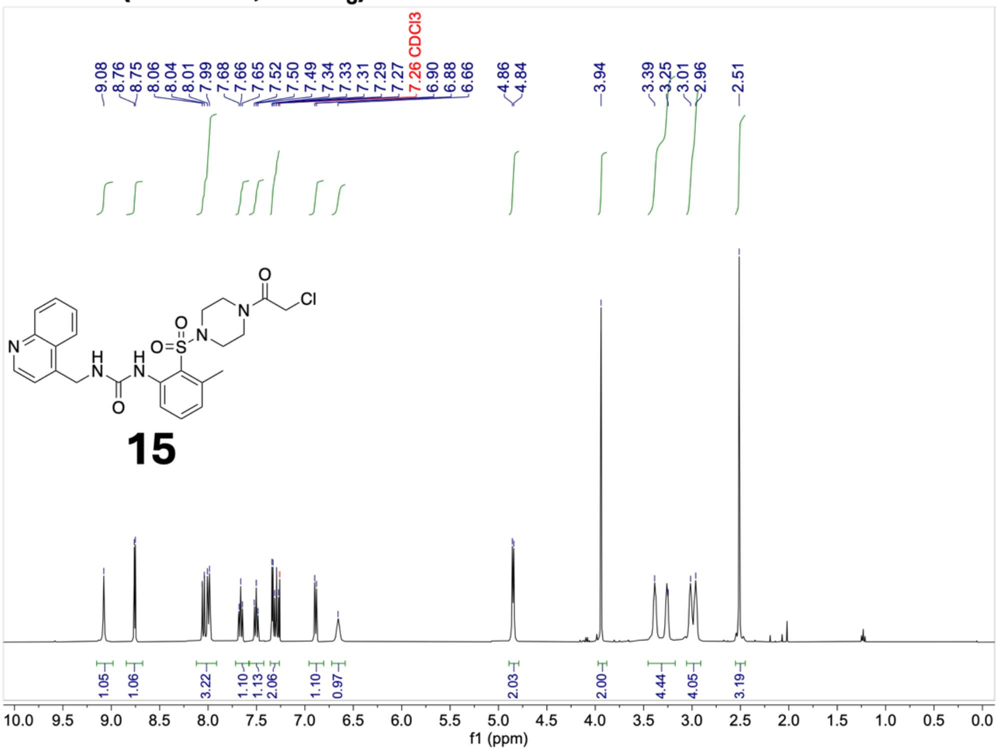


#### $^{13}$C-NMR (101 MHz, $CDCl_3$)

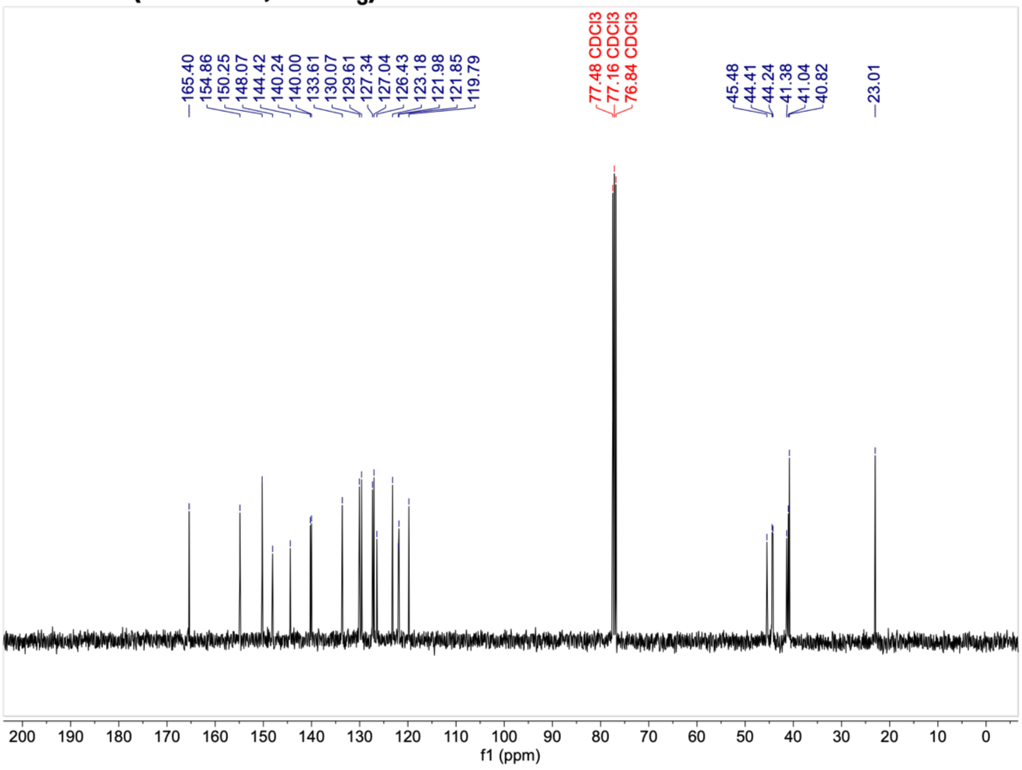

$^{1}$H-NMR (400 MHz, $CDCl_3$)

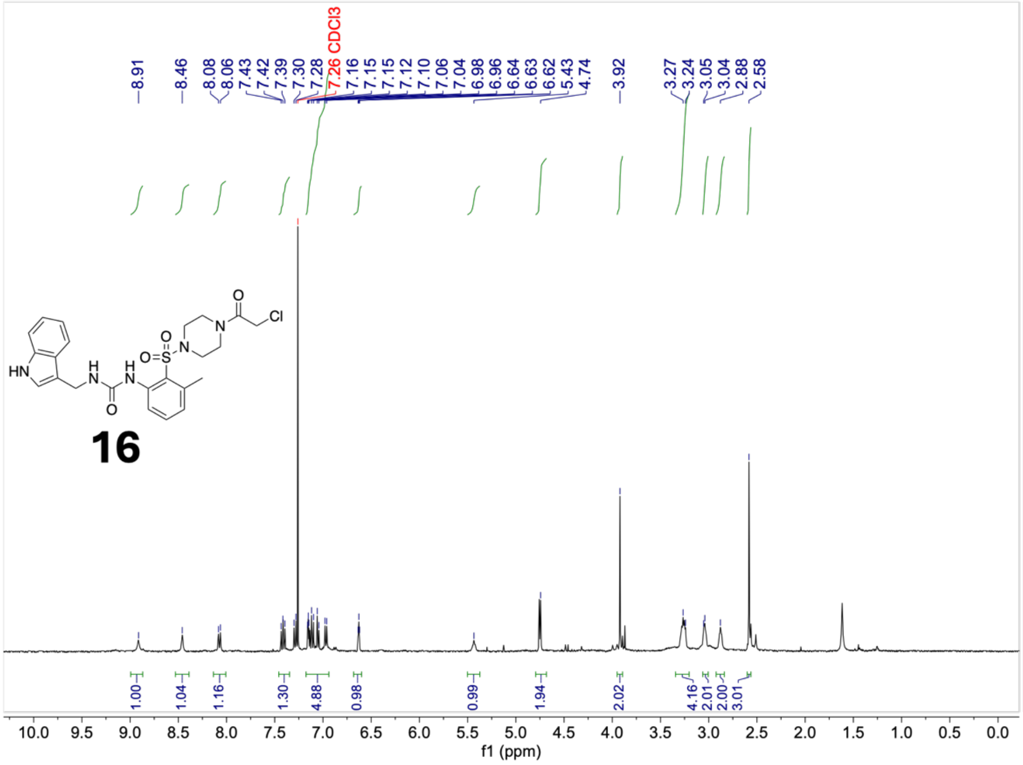


$^{13}$C-NMR (101 MHz, $CDCl_3$)

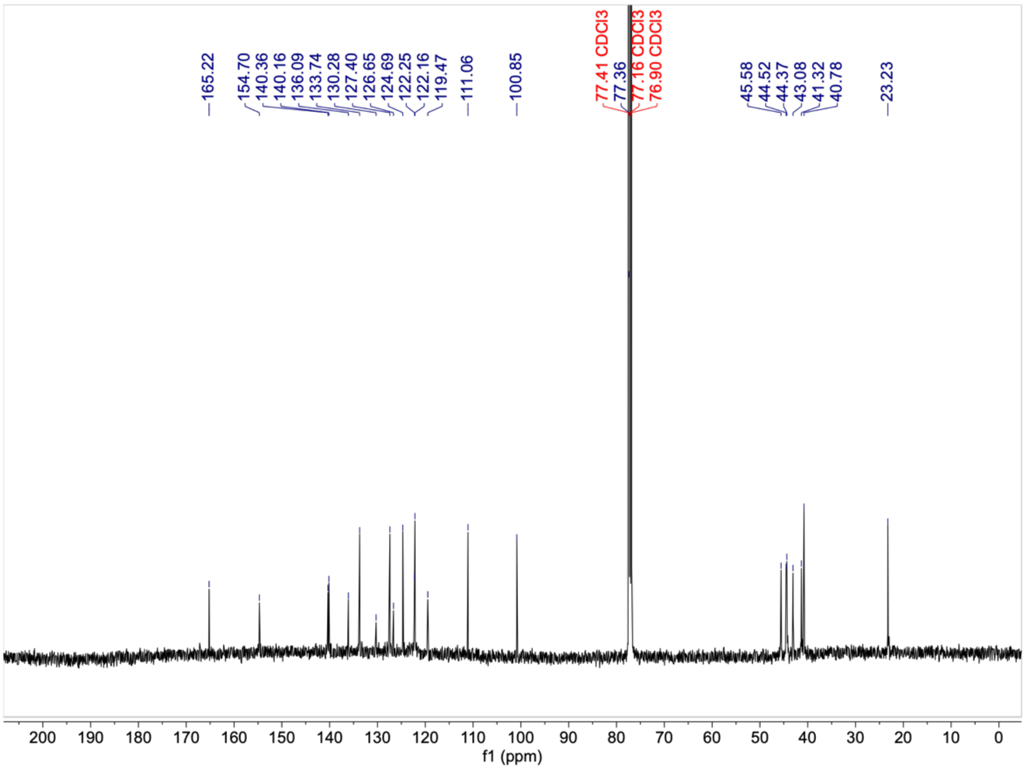

#### $^{1}H$-NMR (400 MHz, $CDCl_3$)

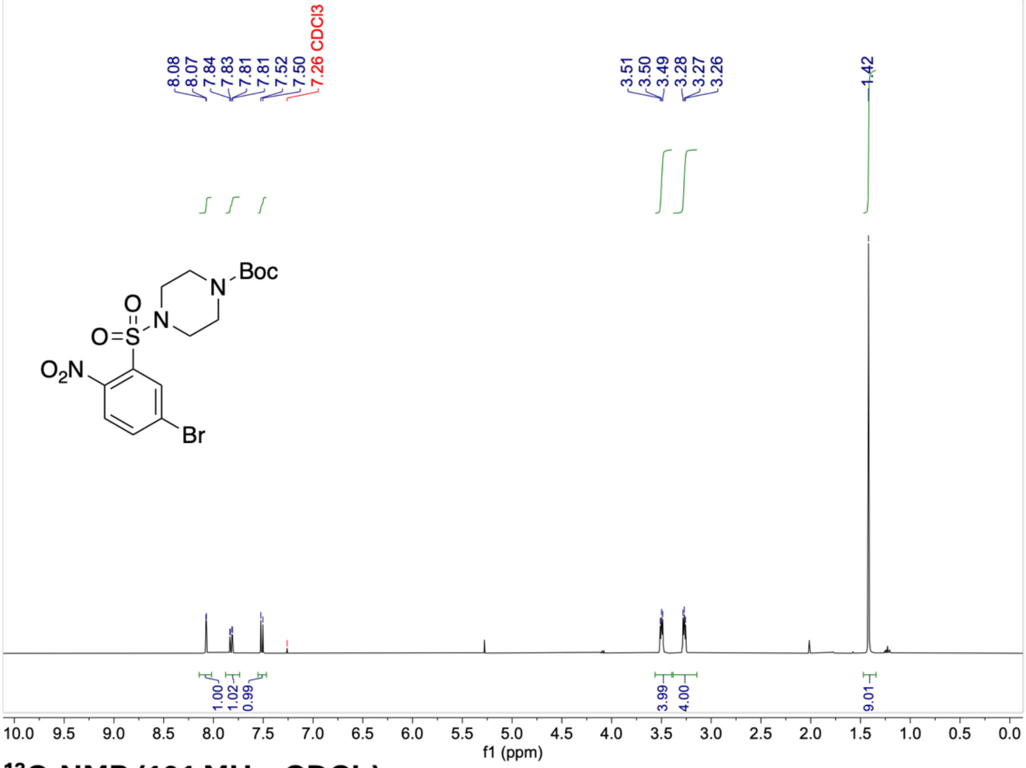


#### $^{13}C$-NMR (101 MHz, $CDCl_3$)

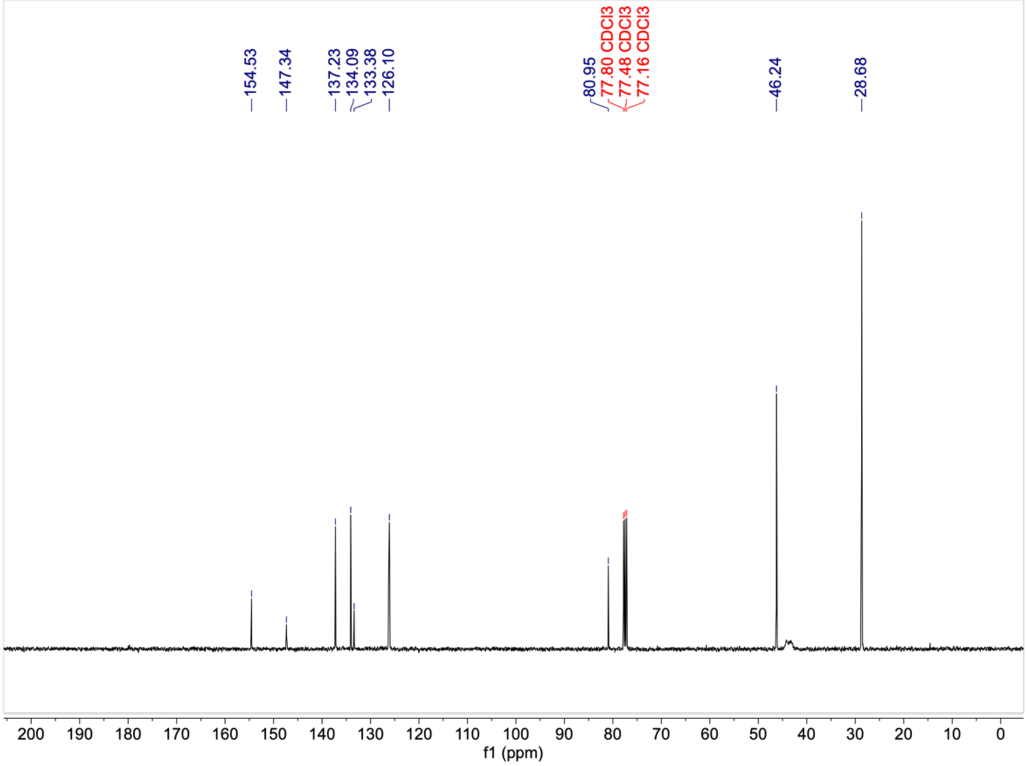

[1]H-NMR (400 MHz, $CDCl_3$)

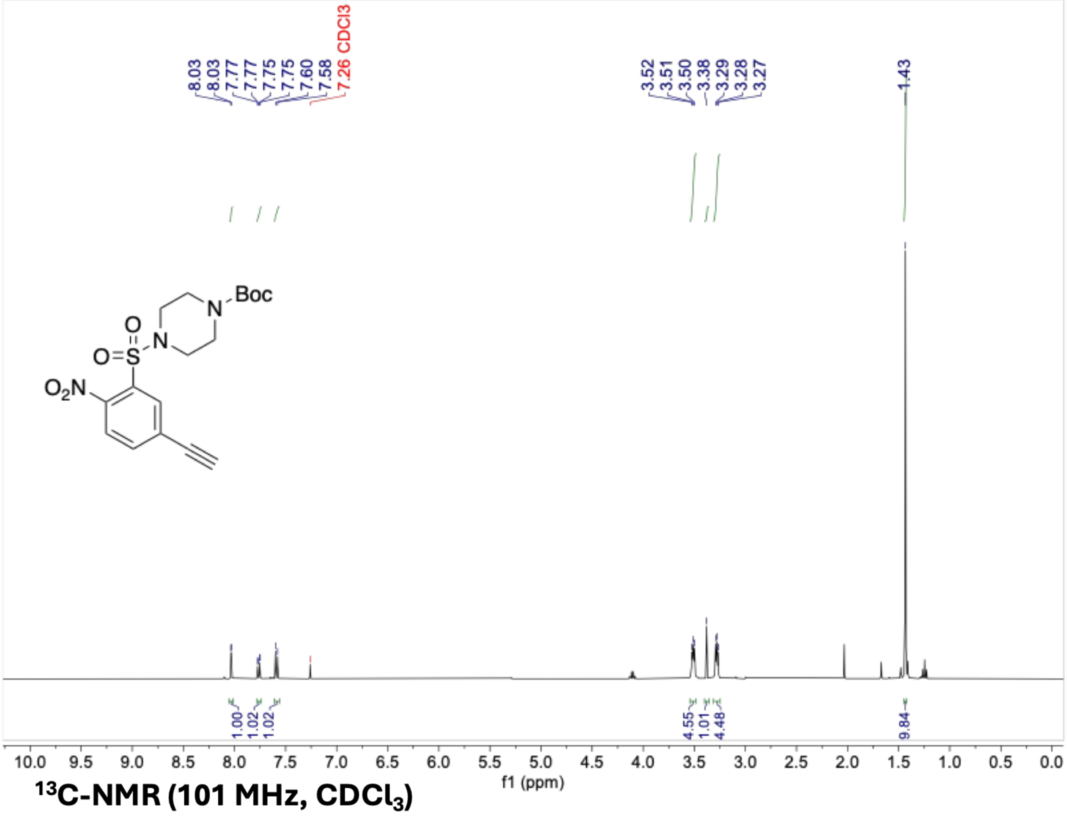

[13]C-NMR (101 MHz, $CDCl_3$)

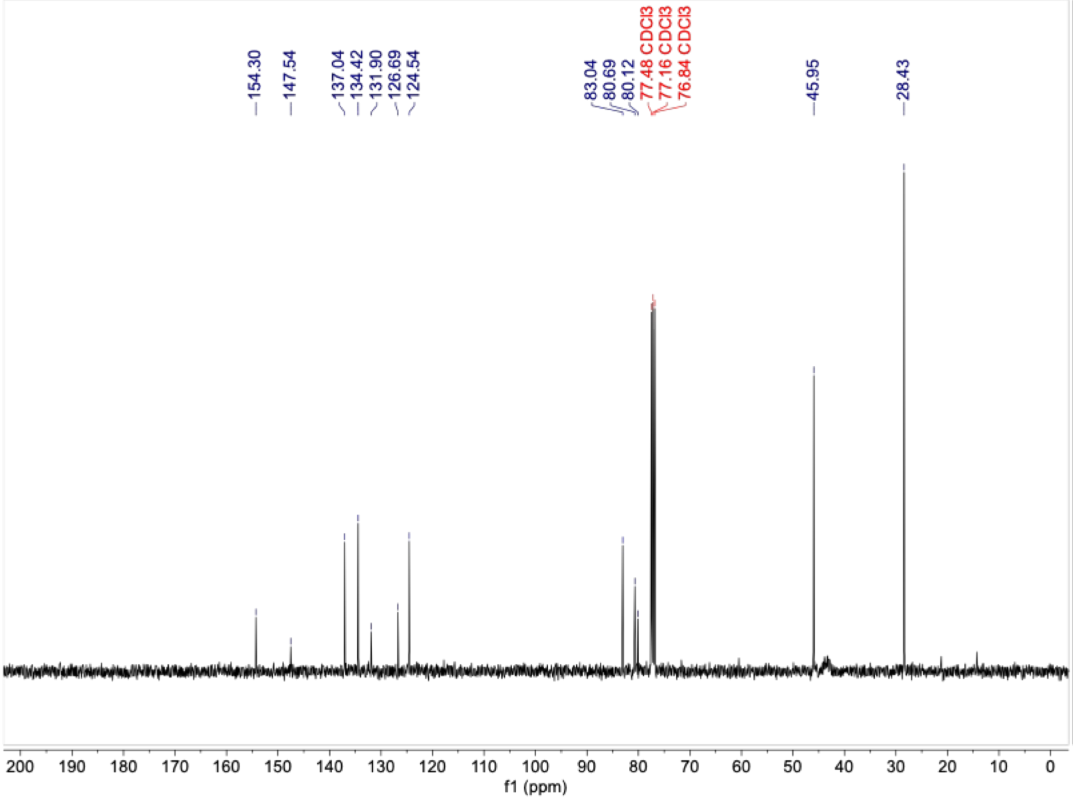

[1]H-NMR (400 MHz, $CDCl_3$)

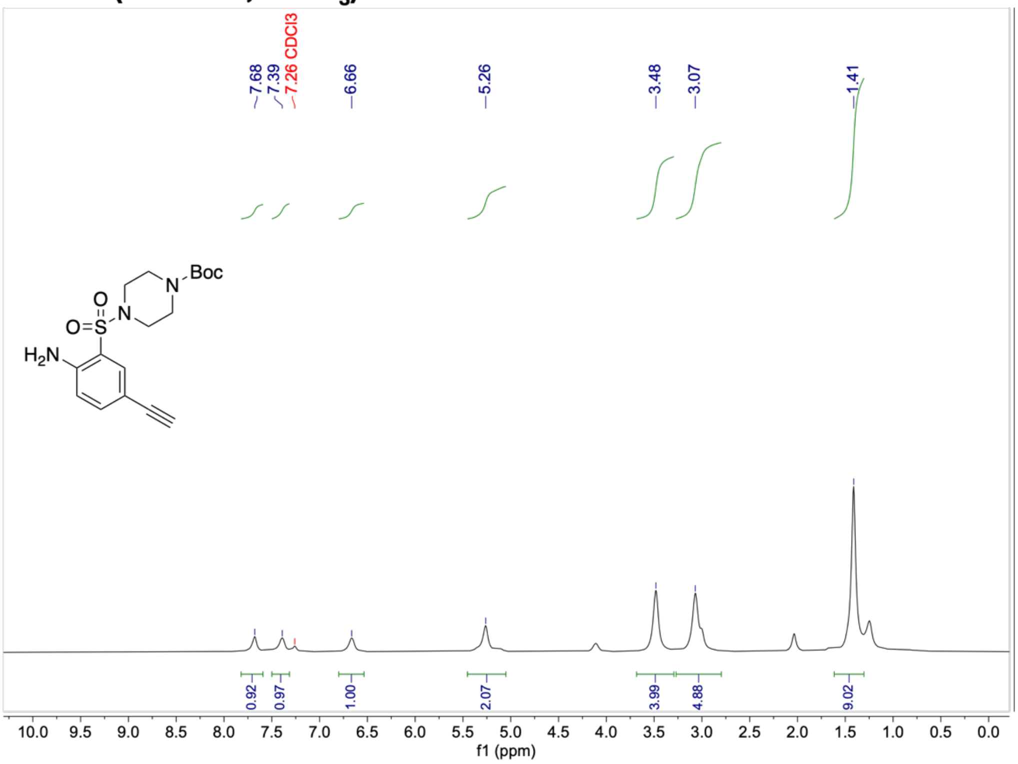


[13]C-NMR (101 MHz, $CDCl_3$)

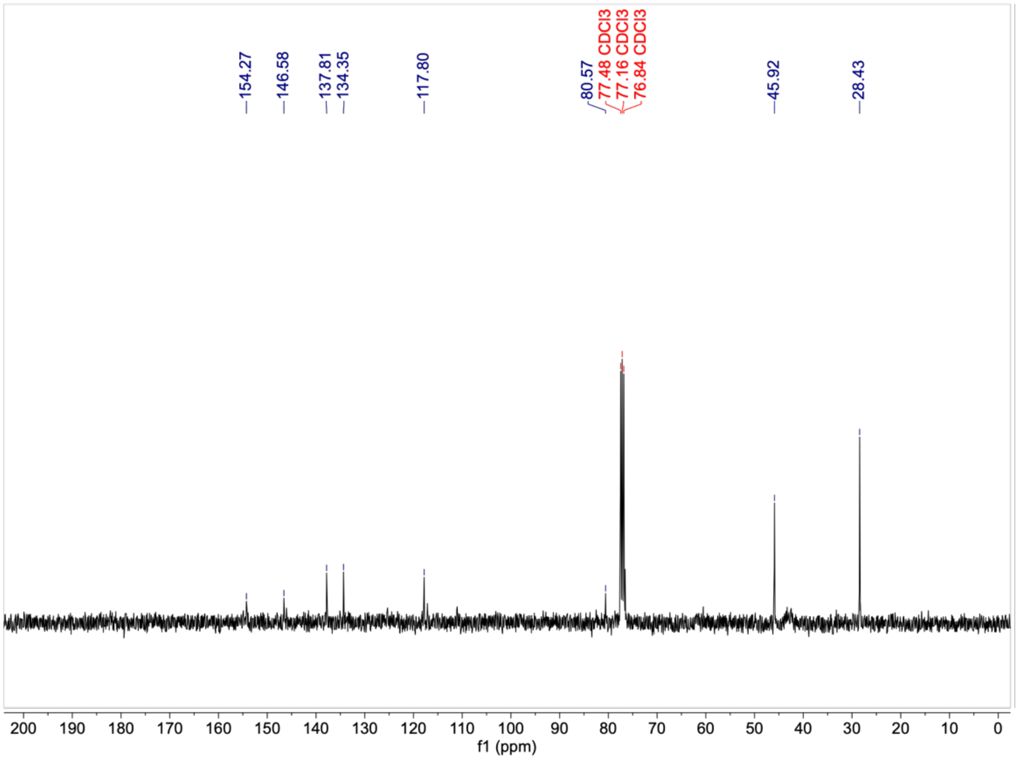

$^{1}H$-NMR (400 MHz, $CDCl_3$)

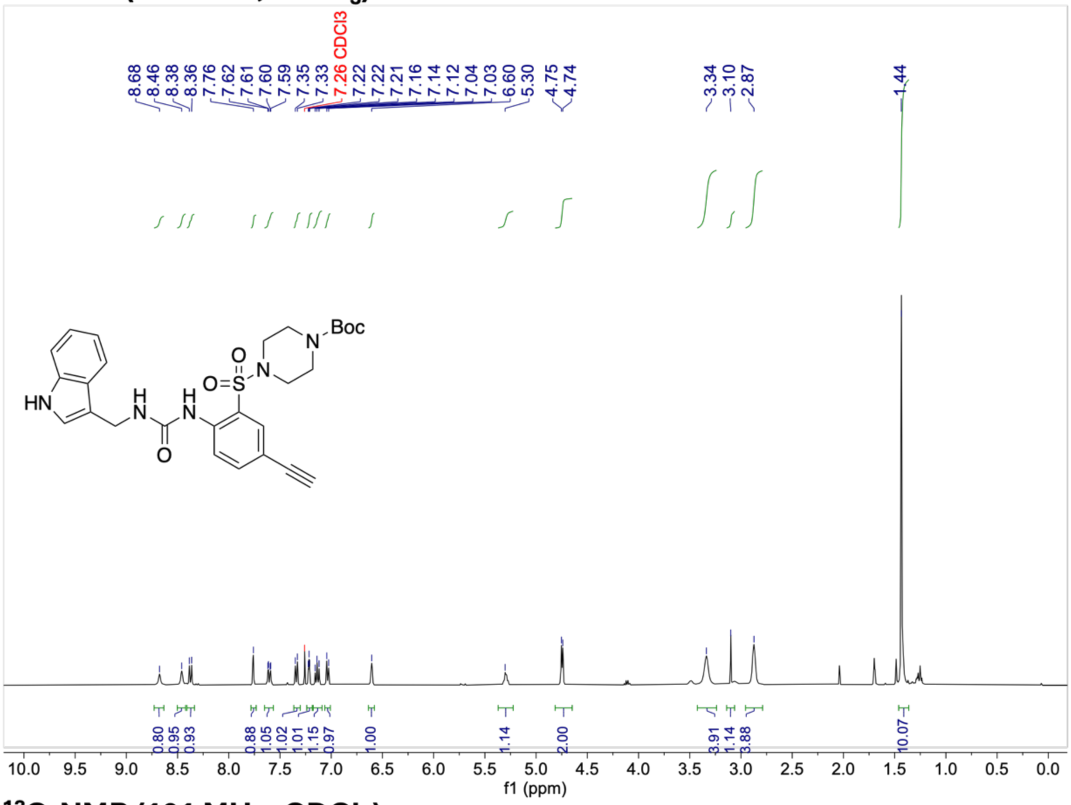

$^{13}C$-NMR (101 MHz, $CDCl_3$)

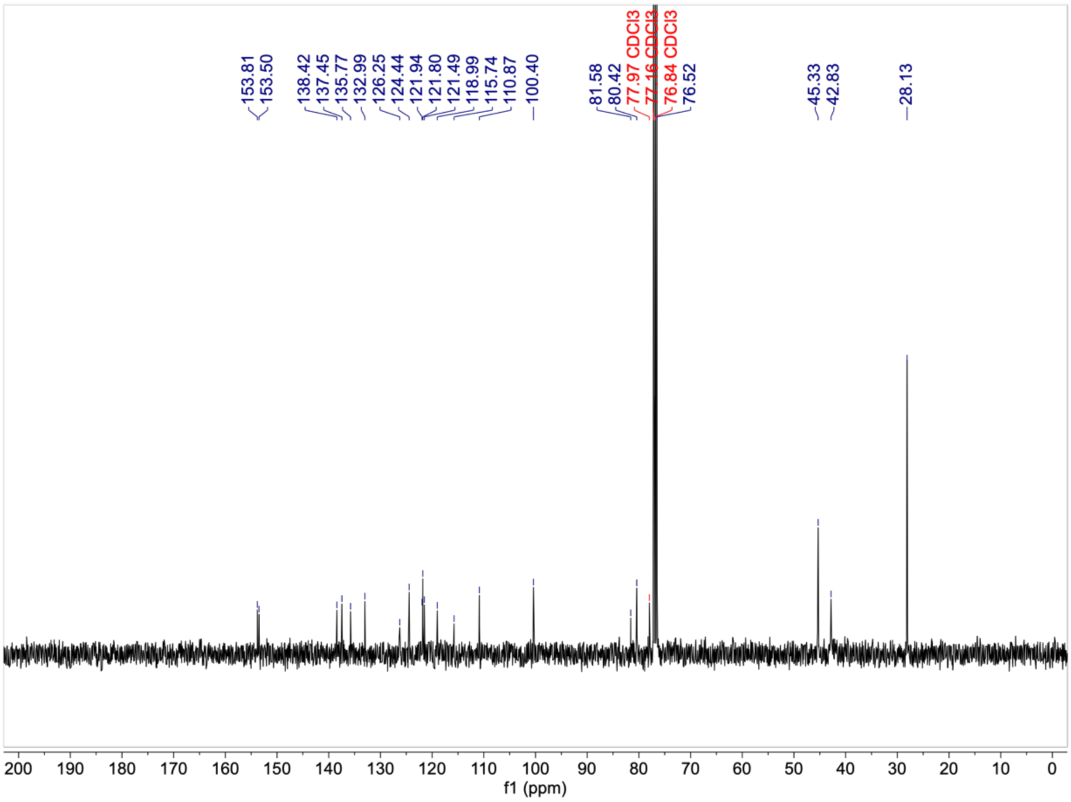

#### [1]H-NMR (400 MHz, $CDCl_3$)

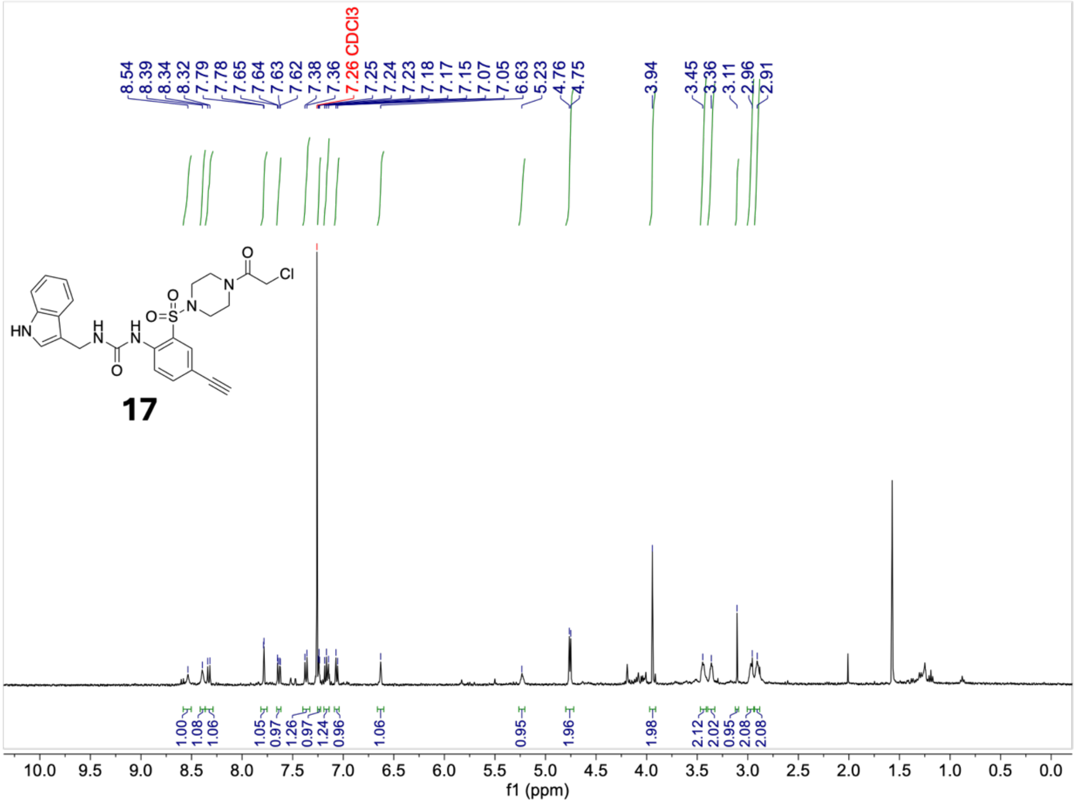

#### [13]C-NMR (101 MHz, $CDCl_3$)

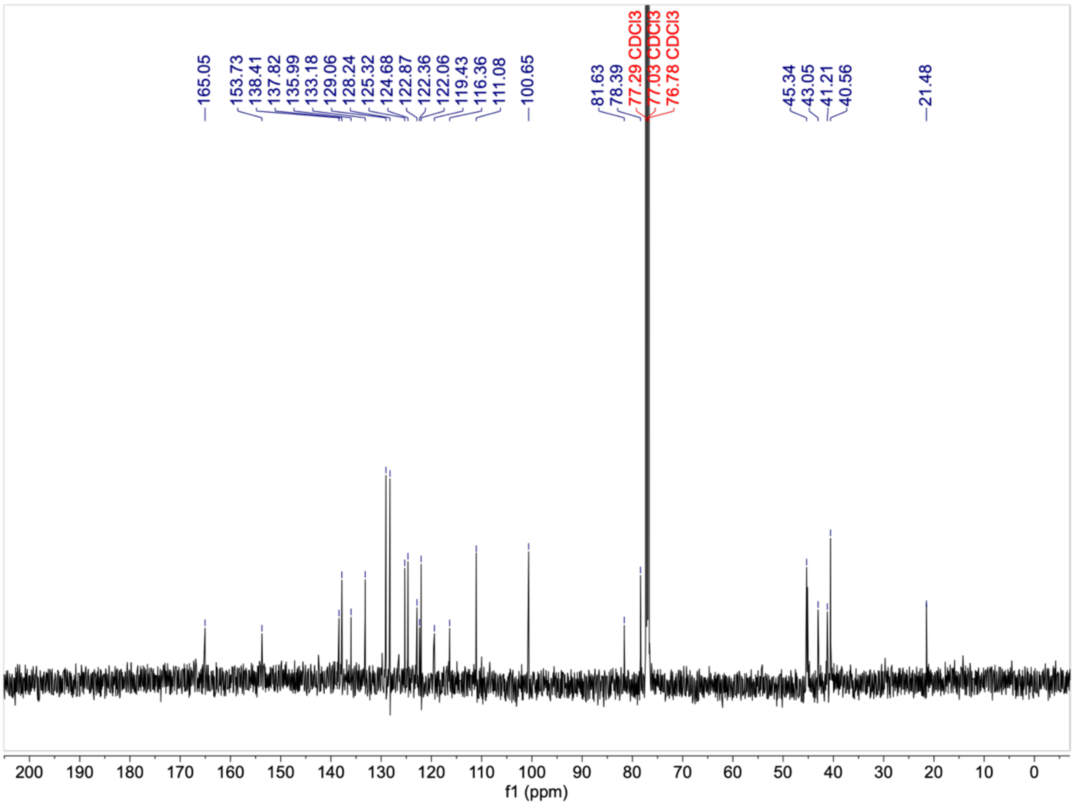
